# Material-driven fibronectin assembly enables low-dose BMP-2 to regenerate structurally mature bone in critical-sized load-bearing defects

**DOI:** 10.64898/2026.10.28.761843

**Authors:** Oihane Mitxelena-Iribarren, Emma Muiños-López, Leire Iturriaga, Irati Aiestaran-Zelaia, Daniel Werner, Jonathan A. Williams, Vineetha Jayawarna, Penelope M. Tsimbouri, Tania López-Martínez, Rafael Llombart-Blanco, Oliver Spaeker, Laura Zorzetto, Wolfgang Wagermaier, Sophie Anuth, Jussi-Petteri Suuronen, Timm Weitkamp, Bernhard Hesse, Peter G. Childs, Stuart Reid, Eva González-Lavado, Diana Martinez-Redondo, Matthew J. Dalby, Froilán Granero-Moltó, Manuel Salmeron-Sanchez, Amaia Cipitria

**Affiliations:** Group of Bioengineering in Regeneration and Cancer, Biogipuzkoa Health Research Institute, San Sebastian, Spain; Cell Therapy Area, Clínica Universidad de Navarra, Pamplona, Spain; Department of Orthopedic Surgery and Traumatology, Clínica Universidad de Navarra, Pamplona, Spain; Instituto de Investigaciones Sanitarias de Navarra (IdiSNA), Pamplona, Spain; Max Planck Institute of Colloids and Interfaces, Department of Biomaterials, Potsdam Science Park, Am Mühlenberg 1, 14476 Potsdam, Germany; Centre for the Cellular Microenvironment, Department of Biomedical Engineering, University of Strathclyde, Glasgow, United Kingdom; Centre for the Cellular Microenvironment, School of Engineering, University of Glasgow, Glasgow, United Kingdom; ioviso GmbH, Am Mühlenberg 11, 14476, Potsdam, Germany; Xploraytion GmbH, Tieckstrasse 29, 10115, Berlin, Germany; Department of Cell Biology and Histology, Faculty of Medicine and Nursing, University of the Basque Country EHU, Leioa, Spain; Synchrotron SOLEIL, Saint Aubin, France; Histocell S.L., Bizkaia Technology Park, Derio, Spain; Institute for Bioengineering of Catalonia (IBEC), The Barcelona Institute for Science and Technology (BIST), Barcelona, Spain; Institució Catalana de Recerca i Estudis Avançats (ICREA), Barcelona, Spain; IKERBASQUE, Basque Foundation for Science, Bilbao, Spain

**Keywords:** Critical-sized bone defect, multiscale characterization, monetite, poly(ethyl acrylate) (PEA), recombinant human bone morphogenetic protein-2 (rhBMP-2), fibronectin, bone regeneration

## Abstract

Bone grafts for critical-sized load-bearing defects are usually judged by how much bone forms, and rarely by whether that bone is structurally and mechanically competent. Here, degradable ceramic monetites coated with poly(ethyl acrylate) (PEA) and functionalised with fibronectin (FN) and rhBMP-2 at 10-25 µg/mL, 60-150-fold below the approved clinical concentration, are assessed across length scales. MicroCT showed that only monetites functionalised with rhBMP-2 bridged the defect, with a significant increase in new bone volume in both the monetite vicinity and the whole defect volume. In contrast, platelet-rich plasma (PRP) functionalisation significantly reduced bone formation relative to FN, showing that the identity of the adhesion cue, and not merely its presence, determines the outcome. Monetite degradation and increased interconnected porosity favoured bone ingrowth and produced a continuous interface between monetite and new bone. The multiscale analyses of quantitative polarised light microscopy, SAXS and nanoindentation show that PEA-coated monetites functionalised with FN and 25 µg/mL rhBMP-2 promoted structurally and mechanically more mature bone, with a hierarchical organisation and local mechanical properties approaching those of native cortical tissue. In the bridged region this group recapitulated the preferential collagen fibre orientation of native cortical bone and reached an elastic modulus and hardness that were not statistically distinguishable from it, although mineral particles throughout the regenerated tissue remained thicker and less aligned than in cortical bone. Synchrotron microCT in an ovine model extended these findings to a clinically relevant scale, supporting the translational potential of the system. Taken together, these findings position PEA-coated monetites functionalised with FN and low-dose rhBMP-2 as a strategy for the structural and functional regeneration of large bone defects.

## 1. Introduction

Critical-sized bone defects in load-bearing sites remain a major clinical challenge in orthopaedic and reconstructive surgery, particularly when spontaneous healing is insufficient ^[1,2]^. Successful regeneration requires re-establishing bone as a hierarchical, mechanically competent tissue, not merely filling a defect radiographically ^[3–5]^. Native bone function arises from structural features spanning multiple length scales, from collagen–mineral organization to tissue architecture, porosity, vascular spaces, and macroscopic load transfer ^[3,5–7]^. Accordingly, bone graft performance should be evaluated not only by the amount of newly formed bone, but also by its spatial distribution and integration with the implanted material.

This perspective is particularly important for large segmental defects with scaffold-guided bone regeneration, where the implanted material acts not merely as a passive filler but as a physical and biological template for tissue formation ^[8,9]^. Pore architecture and scaffold geometry have been shown to direct tissue infiltration and pattern soft and mineralized tissue formation in vivo ^[4]^. In translational load-bearing models, composite scaffolds combined with osteogenic signals have achieved defect bridging and mechanical recovery in large-animal segmental defects, underscoring the clinical relevance of engineered graft strategies ^[10]^. Importantly, growth-factor delivery may enhance the amount and distribution of regenerated tissue without necessarily modifying its intrinsic microstructure ^[11]^. Together, these studies indicate that regeneration should be assessed by where it forms, how it is organized, how it interfaces with the material, and how closely its local structure– property relationships approach those of native bone.

This is equally true of biomaterials based on calcium phosphates, which are widely used for their osteoconductivity and clinical practicality ^[2,12–14]^. However, most granular substitutes remain largely passive, relying predominantly on host-mediated repair, which can be inadequate in critical-sized or biologically compromised defects ^[15,16]^. In this context, monetite, the anhydrous form of dicalcium phosphate, is of particular interest because its resorption can be coupled to new bone formation, but its performance cannot be captured by material degradation or bone volume alone ^[13,17,18]^. Instead, it is necessary to determine how ceramic resorption and bone formation and integration evolve together ^[19,20]^.

A major strategy for enhancing the performance of bone graft substitutes has been incorporating osteoinductive growth factors, particularly bone morphogenetic protein-2 (BMP-2) ^[21,22]^. BMP-2 is a potent regulator of osteogenesis, but its therapeutic application remains constrained by short biological half-life, poor local retention, burst release from conventional carriers, and the need for high or supraphysiological doses to achieve consistent efficacy ^[23,24]^. In fact, the EMA and FDA-approved dosage of the rhBMP-2 product is 1.5 mg/mL — a figure based on nonhuman primate data (where 0.75 mg/mL was considered a low-dose, 2.0 mg/mL a high-dose) and adopted directly for human use ^[21]^ . These limitations are clinically important because excessive BMP-2 exposure has been associated with adverse effects including ectopic bone formation, inflammation, osteolysis, and cancer ^[25]^. Thus, the challenge is not simply to load BMP-2 into a graft material, but to present it at the material interface in a controlled manner, and at a low, biologically efficient dose.

One promising route to achieving this is surface engineering of the graft, rather than bulk growth-factor incorporation. Poly(ethyl acrylate) (PEA) is particularly relevant because it promotes the spontaneous organization of fibronectin (FN) into fibrillar nanonetworks after adsorption ^[26–28]^. This organization exposes integrin- and growth-factor-binding domains, enabling synergistic signalling between cell adhesion receptors and osteogenic pathways ^[26,27,29]^. PEA-based interfaces can support highly efficient growth-factor presentation and osteogenic signalling, including BMP-2-driven bone regeneration at doses substantially lower than conventional soluble delivery approaches ^[30–33]^. However, the biological value of such surface-engineered ceramics cannot be established by bone volume alone, but requires analysis across length scales, from interfacial organization and tissue infiltration to collagen alignment and mechanical competence ^[34]^.

Here, we examined degradable PEA-coated monetites functionalized with a FN+rhBMP-2 coating as a bioactive bone graft substitute, and asked how the newly formed bone develops relative to non-treated controls across length scales. We hypothesized that the PEA coating would organize FN into a biologically active fibrillar network, capable of presenting low-dose rhBMP-2 in a synergistic way, thereby enhancing the osteoinductive performance of the monetites and providing the newly formed tissue with properties approaching native bone. In vivo bone formation was assessed in a critical-sized load-bearing 5 mm femoral defect in a rat model. Our ex vivo multiscale material characterization includes: 3D analysis of the amount of new bone formation and monetite degradation by high-resolution laboratory microCT; detailed analysis of in vivo monetite degradation, new bone–monetite integration and bone infiltration by back-scattered electron microscopy (BSE); evaluation of new mineralized and non-mineralized tissue formation by histology; collagen fibre alignment by polarized light microscopy, complemented by small angle X-ray scattering (SAXS) analysis of nm-scale mineral particle organization; and, finally, quantification of new bone stiffness and hardness by nanoindentation. In addition, synchrotron microCT imaging was applied to an ovine bone-defect model to corroborate the results. More broadly, by integrating 3D morphometry, ceramic degradation analysis, histology, electron microscopy, collagen/mineral organization, and local mechanics, this work aims to establish a surface-engineering strategy for converting monetites from osteoconductive fillers into bioactive materials serving as medical devices for bone regeneration, in which biological performance and tissue maturation can be evaluated across the same hierarchical length scales that define native bone function.

## 2. Results

### 2.1. PEA-coated monetites promote protein adsorption and osteogenic differentiation while preserving cytocompatibility in vitro

Prior to in vivo testing, surface functionalisation of monetites with plasma-polymerised PEA was confirmed by X-ray photoelectron spectroscopy (XPS) (Figure S1a). PEA functionalisation significantly enhanced adsorption of both FN and rhBMP-2 compared with uncoated monetites (Figure S1b,c). Cytocompatibility of the functionalised monetites was evaluated by LIVE/DEAD staining after 3 and 10 days of culture. Viable human adipose stromal cells (hASCs) extensively covered the monetite surfaces, with very few dead cells observed throughout (Figure S1d), indicating that plasma polymerisation and subsequent protein functionalisation did not adversely affect cell viability. The osteogenic potential of the monetites was assessed by quantitative real-time PCR after 28 days of culture. PEA-coated monetites functionalised with FN and rhBMP-2 exhibited significantly higher expression of the osteogenic markers osteocalcin (*BGLAP*, Figure S1e) and osteonectin (*SPARC*, Figure S1f) than the corresponding uncoated monetites, indicating that PEA-mediated protein presentation promoted osteogenic differentiation. As expected, the osteogenic medium (OM) positive control produced the highest overall gene expression, although statistically significant differences relative to the PEA+FN+rhBMP-2 group were observed only for *BGLAP*.

### 2.2. Multiple monetite implantation increases rat critical-sized bone defect regeneration

Once the coating was validated in vitro, we compared new bone formation after 10 weeks following implantation of a single monetite with 5 mm diameter and 3 mm height (Figure S2a) versus two monetites with 2.5 mm diameter and 1.5 mm height (Figure 1a, S2b). The former represents the material intended for human applications; the latter is a smaller version adapted to the rat defect model. Animals received either one (“1M”) or two monetites (“2M”), plain or coated with PEA and functionalised with FN and 10 µg/mL rhBMP-2 (“PFB10”, Figure S2c).

**Figure 1.**
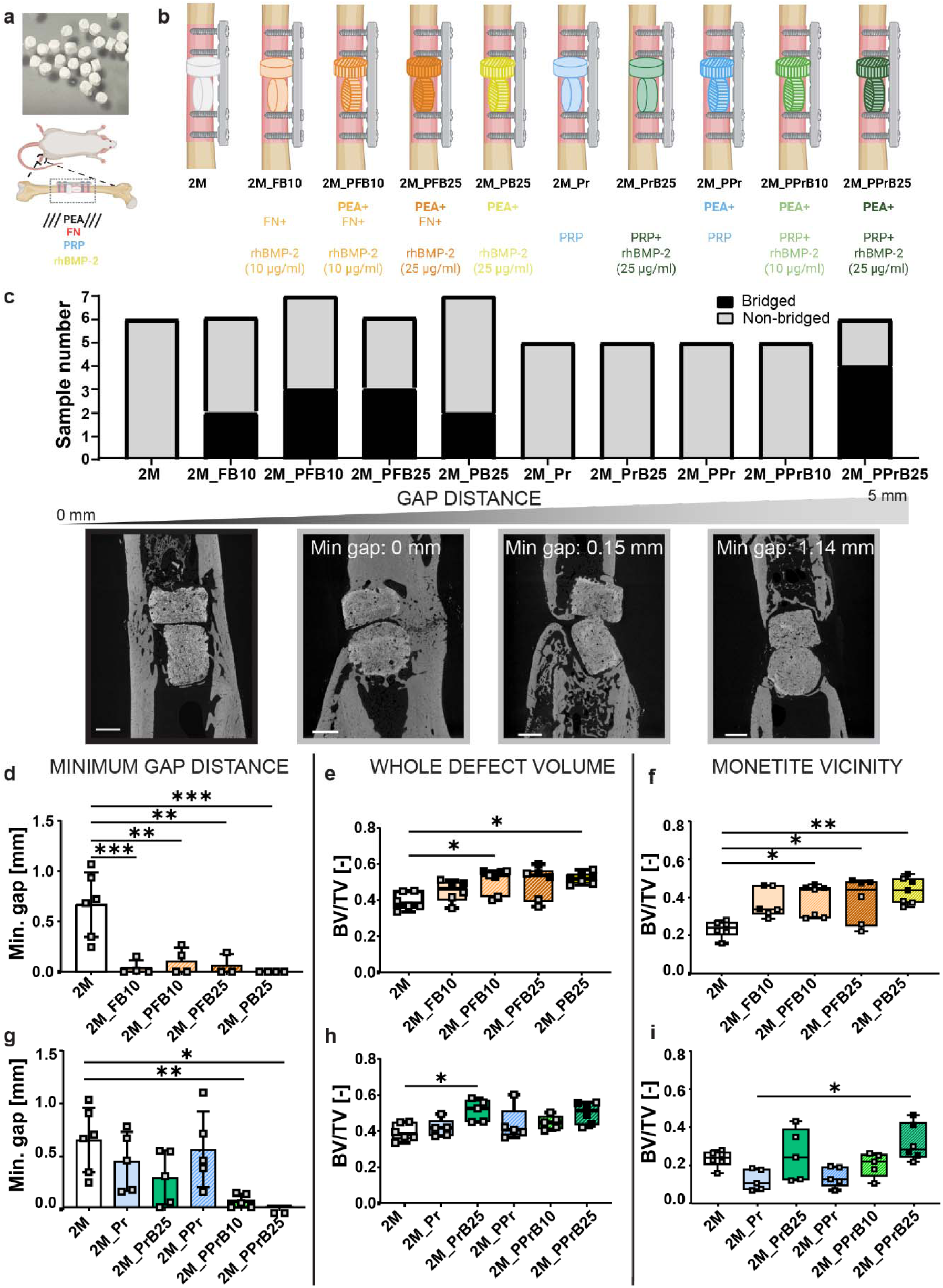
Bone formation after monetite implantation in a rat model. a) Picture of small monetites tested in rat femora: placing two monetites with 2.5 mm diameter and 1.5 mm height held with a plate and four screws. Figure created with BioRender.com. b) Different combinations of the coating and functionalization used in the study, represented each by a different colour or pattern: PEA (P, stripes), FN (F, red), PRP (Pr, blue) and rhBMP-2 (B, yellow) in two different concentrations (B10 and B25, darker when higher concentration). c) Summary of bone bridging analysis for the rats containing two monetites, showing fully bridged samples in black and non-bridged samples in grey. Images of exemplary non-bridged samples show their partial bridging with the measured minimum gap distance between the distal and the proximal bone. Scale bar: 1 mm. Dragonfly analysis results of the high-resolution laboratory microCT images quantifying the minimum gap distance between proximal and distal bone (d, g), the bone volume per total volume in the whole defect (e, h) and in the vicinity of the monetite (f, i). Colour and pattern of each bar plot and boxplot is determined by the mixture of coatings in each experimental group. Fully bridged samples appear as black squares, while grey ones represent non-bridged animals in all graphs. Bar plots show the mean ± standard deviation per group, while boxplots show the median and interquartile range. Comparisons between groups were performed using one-way ANOVA, with Bonferroni post-hoc correction (parametric cases shown with bar plots) or its equivalent non-parametric Kruskal-Wallis test (boxplots). Statistical significance is indicated with * p<0.05, **p<0.01, ***p<0.001. Figures created with BioRender.com.

Defects were classified as fully bridged when new cortical bone with no visible monetite spanned the whole defect region (Figure S2d, black), or non-bridged when a gap remained between the distal and proximal cortical bone (Figure S2d, grey). None of the plain monetites (1M or 2M) resulted in defect bridging, while similar bridging was observed with the coated monetites (2/6 for 1M_PFB10 and 3/7 for 2M_PFB10, Figure S2e). Two monetites significantly increased new bone volume, in both the whole defect (Figure S2f) and the monetite vicinity (defined as the monetite volume dilated by 10%; Figure S2g). In addition, within the two-monetite configuration, coated monetites showed significantly higher new bone volume than plain ones, both in the whole defect and in the monetite vicinity, demonstrating the osteogenic effect of the coating. Thus, for subsequent analyses of the rat critical-sized defect, we used two small monetites per defect across the different coating conditions.

### 2.3. PEA coating and low-dose rhBMP-2 increase 3D bone bridging and regeneration with and without FN functionalization

Next, we analysed the effect of two small monetites per defect (Figure 1a) with and without PEA coating (“P”) and functionalisation with selected combinations of FN (“F”) and/or rhBMP-2 at 10 µg/mL (“B10”) or 25 µg/mL (“B25”) (Figure 1b).

rhBMP-2 at either concentration, alone or combined with FN and/or PEA resulted in full bridging in nearly half of the cases (Figure 1c). For non-bridged defects, the minimum gap distance between proximal and distal bone was measured as previously described ^[35]^. Partial bridging was confirmed by the significant difference in minimum gap distance across all experimental groups compared to control (Figure 1d).

In the whole defect volume, BV/TV was higher than the plain monetite in the four functionalised experimental groups (2M_FB10, 2M_PFB10, 2M_PFB25, and 2M_PB25), reaching statistical significance for 2M_PFB10 and 2M_PB25 (Figure 1e). In addition, BV/TV in the vicinity of the monetite increased significantly in all groups containing PEA-coated monetites (2M_PFB10, 2M_PFB25 and 2M_PB25), confirming the coating’s efficacy for new bone formation (Figure 1f). Together, these results confirm the efficacy of PEA coating and rhBMP-2 functionalization, at both concentrations, alone or together with FN, for bone regeneration in a critical-sized defect.

### 2.4. PRP coating decreases 3D bone bridging and new bone formation if not combined with PEA and rhBMP-2

As GMP-grade recombinant FN suitable for clinical manufacture is not readily available, we evaluated platelet-rich plasma (PRP) as a clinically relevant alternative. While FN is an extracellular matrix protein that supports cell adhesion and migration through integrin-mediated signalling ^[36]^, human PRP could offer a more complex autologous biological environment to promote cell attachment, proliferation, angiogenesis, extracellular matrix remodelling, and tissue repair ^[37–39]^. In vitro, PRP-functionalized surfaces also exhibited a network-like protein organisation similar to that observed with FN functionalization (Figure S3a). Despite donor variability, PRP from four of five donors retained rhBMP-2 at levels comparable to monetites functionalized with FN (Figure S3b). In addition, *BGLAP* expression in human adipose-derived stem cells (hASCs) cultured on PRP-functionalized monetites varied between PRP donors (Figure S3c), reflecting the biological variability of autologous PRP. Nevertheless, monetites functionalized with PRP and rhBMP-2 from two donors induced *BGLAP* expression comparable to the FN+rhBMP-2 combination, whereas monetites functionalized with PRP alone produced lower osteogenic responses. Therefore, we evaluated PRP as an alternative to FN for monetite functionalisation in the rat model.

In vivo, however, PRP-based functionalisation did not perform comparably to FN in terms of defect bridging and new bone formation. No bridging was observed in most PRP-functionalized groups, except for the combination of PRP (“Pr”) with PEA and 25 µg/ml rhBMP-2 (Figure 1c). In addition, only groups combining PRP with PEA and rhBMP-2 showed a significant difference in minimum gap distance (Figure 1g), confirming that PRP alone lacks efficacy.

Similarly, in the whole defect volume, only group 2M_PrB25 showed significantly higher BV/TV than the control (Figure 1h), reflecting lateral bone growth rather than bridging. In the vicinity of the monetite, BV/TV decreased in all groups, except for 2M_PPrB25, which was significantly higher than 2M_Pr (Figure 1i).

Based on bridging outcomes and statistically significant bone volume increases relative to the plain monetite group, five groups were selected for the multiscale tissue characterization: 2M_FB10, 2M_PFB10, 2M_PFB25, 2M_PB25, and 2M_PPrB25.

### 2.5. Monetite degradation and increased interconnected porosity favour bone-monetite integration and bone infiltrations in PEA-coated groups

Next, we evaluated monetite degradation in vivo after 10 weeks of implantation in the critical-sized femoral defect. To do so, high-resolution laboratory microCT scans (2 μm voxel) were performed before implantation (“pre-implant”) and compared with explants after 10 weeks of implantation (Figure 2a, b, e, f). Monetite volume (excluding porosity) (Figure 2c, g), overall porosity, closed and interconnected porosity (Figure 2d, h) were quantified with self-developed deep learning models in ORS Dragonfly software for both plain and PEA-coated monetites. Monetite volume decreased significantly in both plain and PEA-coated groups after 10 weeks in vivo, with a drop of nearly 20% (Figure 2c, g). In contrast, overall porosity increased significantly only in PEA-coated monetites, by 25%, with no change in plain monetites (Figure 2d, h). This increase in overall porosity resulted from a roughly twofold decrease in closed porosity and a twofold increase in the interconnected porosity (Figure 2h), favouring new bone ingrowth.

**Figure 2.**
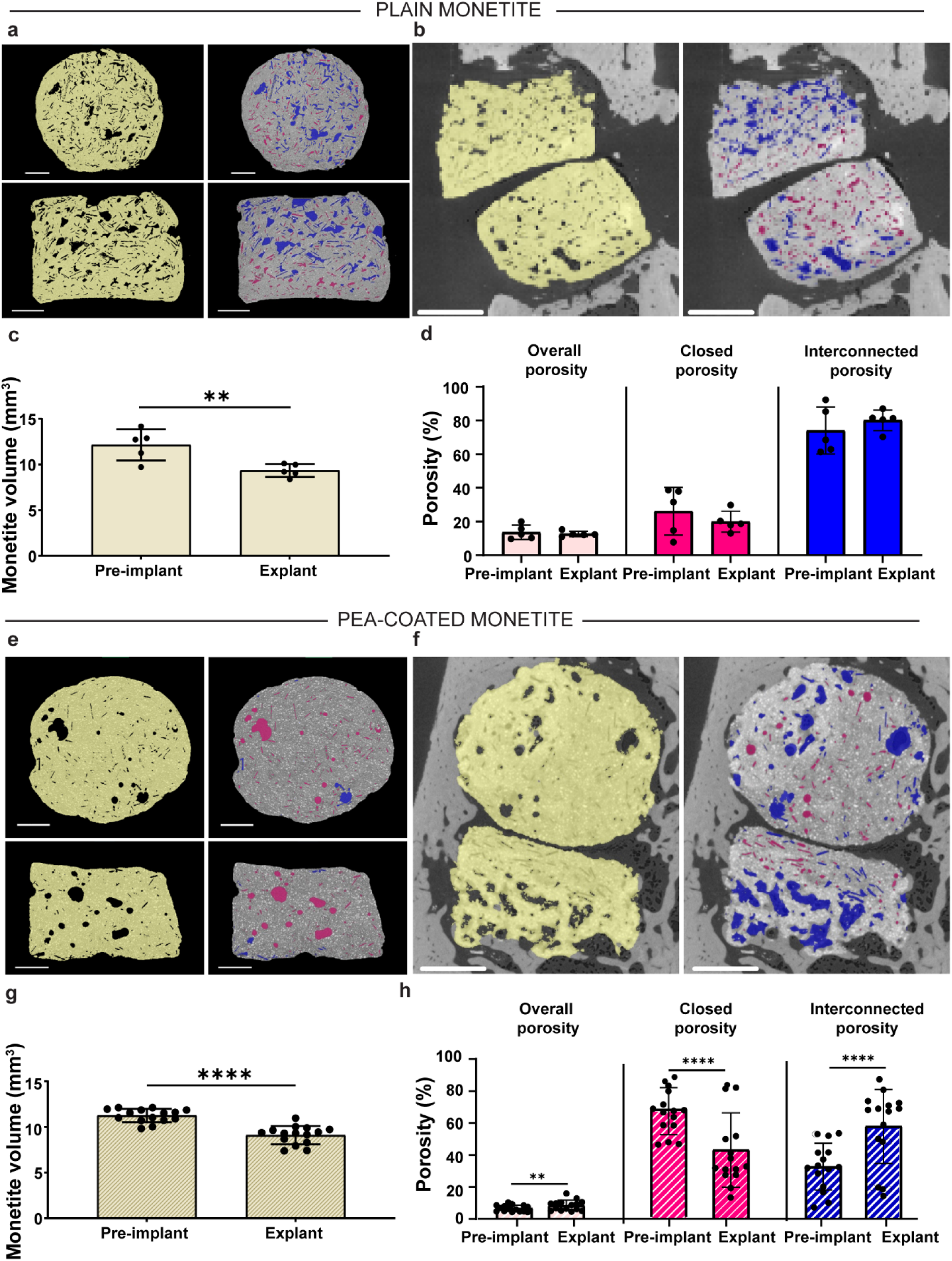
Monetite in vivo degradation analysis. High resolution laboratory microCT images of plain monetites before implantation (pre-implant) (a) and in the explants after 10 weeks in vivo (explant) (b), showing the monetite volume in yellow, the closed porosity in pink and interconnected porosity in blue. Quantification of plain monetite volume (c) and porosity percentage (d), including overall, closed and interconnected porosity. Similarly, high resolution microCT images of a PEA-coated monetites before implantation (e) and in the explants (f), showing the monetite volume in yellow, the closed porosity in pink and interconnected porosity in blue. Quantification of PEA-coated monetite volume (g) and porosity percentage (h), including overall, closed and interconnected porosity. Bar plots show the mean and standard deviation per group, and dots represent each analysed monetite, being n = 5 in non-coated monetites and n = 15 in PEA-coated monetites. Pair-wise comparisons of the monetite before and after implantation were performed with t-test. Statistical significance is indicated with **p<0.01, ****p<0.0001. Scale bar: 0.5 mm.

New bone integration adjacent to and within monetites was further quantified in BSE images from three representative explants per group (Figure 3a, S4, S5). Good bone-to-monetite surface (%) was observed in 2M_FB10, 2M_PFB10, 2M_PFB25, and 2M_PB25, with a higher trend than the control, indicating a seamless interface and good bone-monetite integration (Figure 3b, S6). This was not the case for group 2M_PPrB25, which showed greater variability, consistent with its poorer bone bridging results. When measuring the number and area of bone infiltrations within monetites, we found more and larger infiltrations in all functionalized groups, with very few in plain monetites (Figure 3c). Both the size and number of infiltrations were significantly higher in the 2M_PFB10 group than in the control. Osteocyte lacunae were clearly identified within these new bone infiltrations, demonstrating new bone ingrowth through the ceramic (Figure S6).

**Figure 3.**
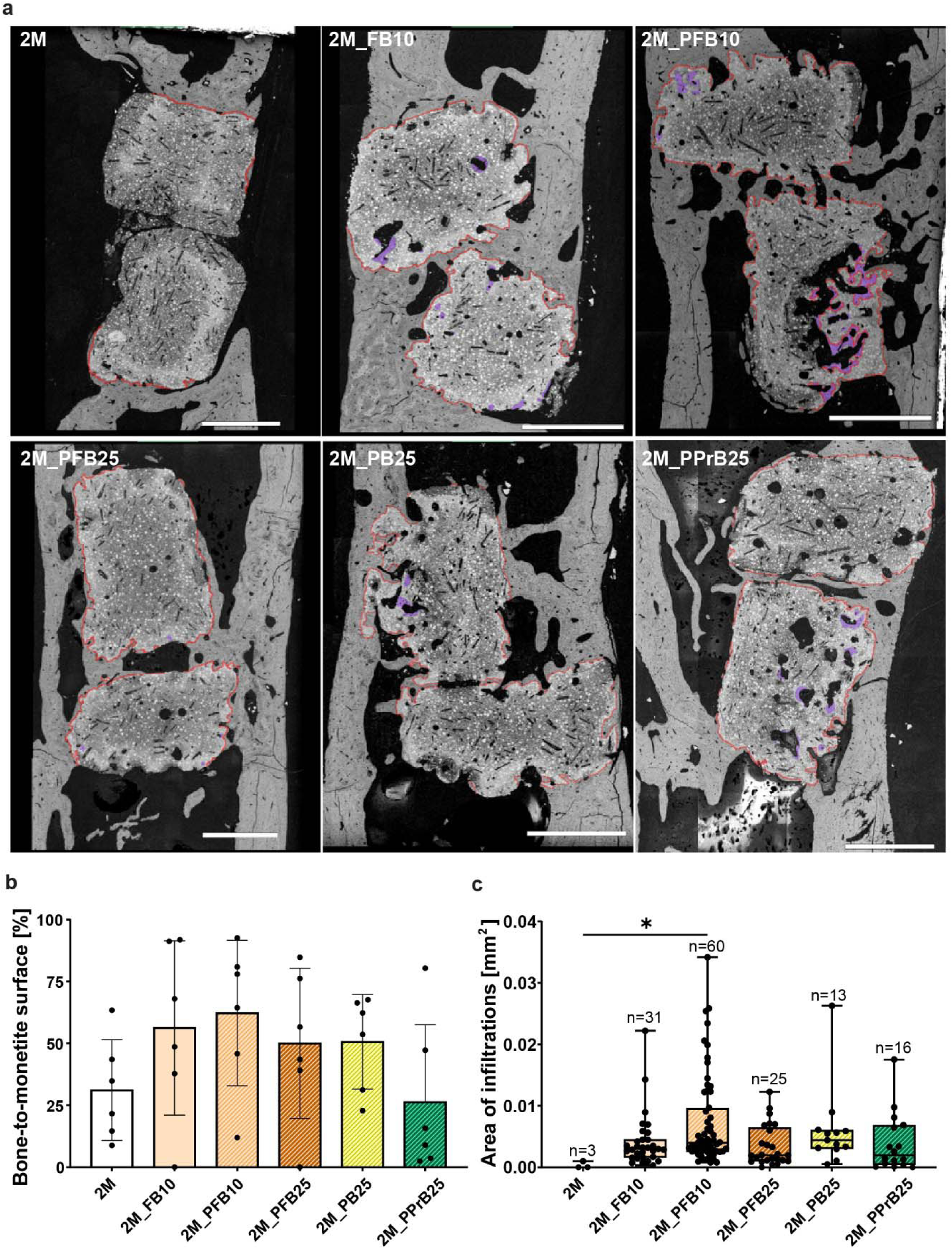
BSE images analysis of new bone integration and bone infiltrations. a) BSE images of one representative sample of the selected experimental groups, including plain monetite (2M), FN+rhBMP-2 10 μg/ml (2M_FB10), PEA+FN+rhBMP-2 10 μg/ml (2M_PFB10), PEA+FN+rhBMP-2 25 μg/ml (2M_PFB25), PEA+rhBMP-2 25 μg/ml (2M_PB25) and PEA+PRP+rhBMP-2 25 μg/ml (2M_PPrB25) groups, showing the monetite in direct contact with new bone (red line) and the bone infiltrations into the monetites (purple). Scale bar: 1mm. b) Quantification of the bone-to-monetite surface (%). Bar plots show the mean and standard deviation per group. Comparisons between groups were done using one-way ANOVA, with Bonferroni post-hoc correction. n = 3 animals per group, with two monetites per sample. c) Area of all infiltrations counted in each experimental group, where each dot represents one infiltration. Total number of infiltrations counted per group are indicated on top of each box. Boxplots show the median and interquartile range per group. Comparisons between groups were done using the non-parametric Kruskal-Wallis test. Statistical significance is indicated with *p<0.05.

### 2.6. PEA-coated and functionalized samples show more non-mineralized matrix and more osteoclasts by 2D histomorphometry

Histological sections obtained from each sample of the six selected groups were used to assess the mineralized and soft tissue surrounding the monetites (n = 5-7 per group). Serial histological sections of non-demineralized samples were obtained using a controlled-angle cutting procedure to reach the region of interest (sagittal plane) ^[40]^, in which monetites surrounded by newly formed tissue were identified using different stains.

Goldner’s trichrome staining was first performed to evaluate bone morphology, remodelling and formation (Figure S7). All samples from the selected groups showed collagen in green, both in the native cortical bone and in the new bone.

Next, von Kossa+Movat pentachrome and tartrate-resistant acid phosphatase (TRAP) stains were performed to assess new mineralized tissue formation (dark brown-black) and osteoclast presence (pink-purple), respectively (Figure 4a). A Fiji macro was used to obtain the ROIs described in Figure S8 (available on Zenodo: 10.5281/zenodo.23324947). New bone area increased slightly in 2D (Figure 4b), although the 3D microCT analysis above is the more sensitive measure. Monetite volume decreased in all groups compared to the control, with a significant drop for the 2M_PB25 group, confirming the results obtained with 3D microCT. Non-mineralized collagenous tissue (soft tissue) increased in all groups compared to the control, with a significant increase for the 2M_PB25 group.

**Figure 4.**
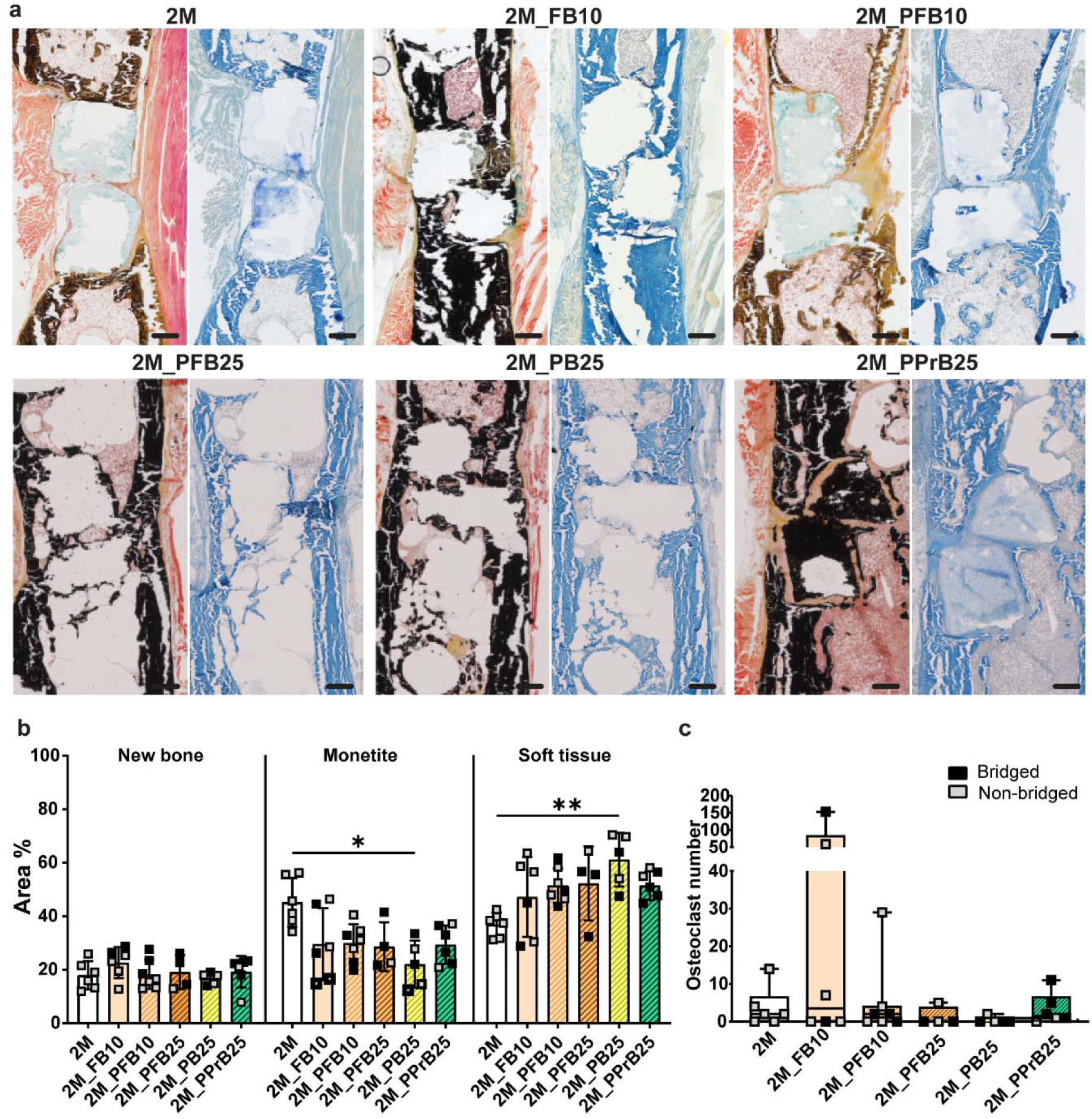
Histological stains for bone mineralisation and histomorphometry measurements. a) Von Kossa+Movat pentachrome (left) and TRAP (right) staining of one representative sample of the selected experimental groups, including plain monetite (2M), FN+rhBMP-2 10 μg/ml (2M_FB10), PEA+FN+rhBMP-2 10 μg/ml (2M_PFB10), PEA+FN+rhBMP-2 25 μg/ml (2M_PFB25), PEA+rhBMP-2 25 μg/ml (2M_PB25) and PEA+PRP+rhBMP-2 25 μg/ml (2M_PPrB25) groups. Scale bar: 500 μm. b) Quantification of the area % of new bone, monetite and soft tissue in each experimental group, where each square represents a fully bridged (black) or partially/non-bridged (grey) defect. Bar plots show the mean and standard deviation per group. Comparisons between groups were done using one-way ANOVA, with Bonferroni post-hoc correction. c) Osteoclast number quantification per group, where boxplots show the median value and interquartile range per group. Each square represents the number of osteoclasts found in a fully bridged (black) or partially/non-bridged (grey) defect. Statistical significance is indicated with *p<0.05, **p<0.01. n = 4-7 per group.

TRAP staining revealed osteoclasts surrounding the monetites in all experimental groups (Figure S9). Osteoclast numbers were higher in non-bridged samples from the 10 μg/ml rhBMP-2 groups (specially 2M_FB10), suggesting greater bone remodelling activity (Figure 4c).

### 2.7. PEA-coated monetites functionalized with FN and low-dose rhBMP-2 promote collagen fibre orientation comparable to cortical bone in newly formed bridged bone

Based on these results, we performed multiscale tissue organisation analysis, from micro-to nanometre scale, in the groups coated with PEA and functionalized with FN and rhBMP-2 at both concentrations (2M_PFB10 and 2M_PFB25), as well as the group with PEA alone with 25 μg/ml rhBMP-2 (2M_PB25). Animals implanted with plain monetites (2M) were used as controls. Alcian blue and picrosirius red staining of serial histological sections, combined with polarized light imaging, showed that groups with PEA+FN coating had higher collagen content. Across all groups, regions surrounding the monetites showed more newly formed collagen (Figure 5a). The analysed ROIs were the cortical region (native cortical bone), bridged region (new bone in the bridged area), interphase region (new bone adjacent to the monetite surface), and intermonetite region (new bone between monetites), as defined in the experimental section (Figure S10).

**Figure 5.**
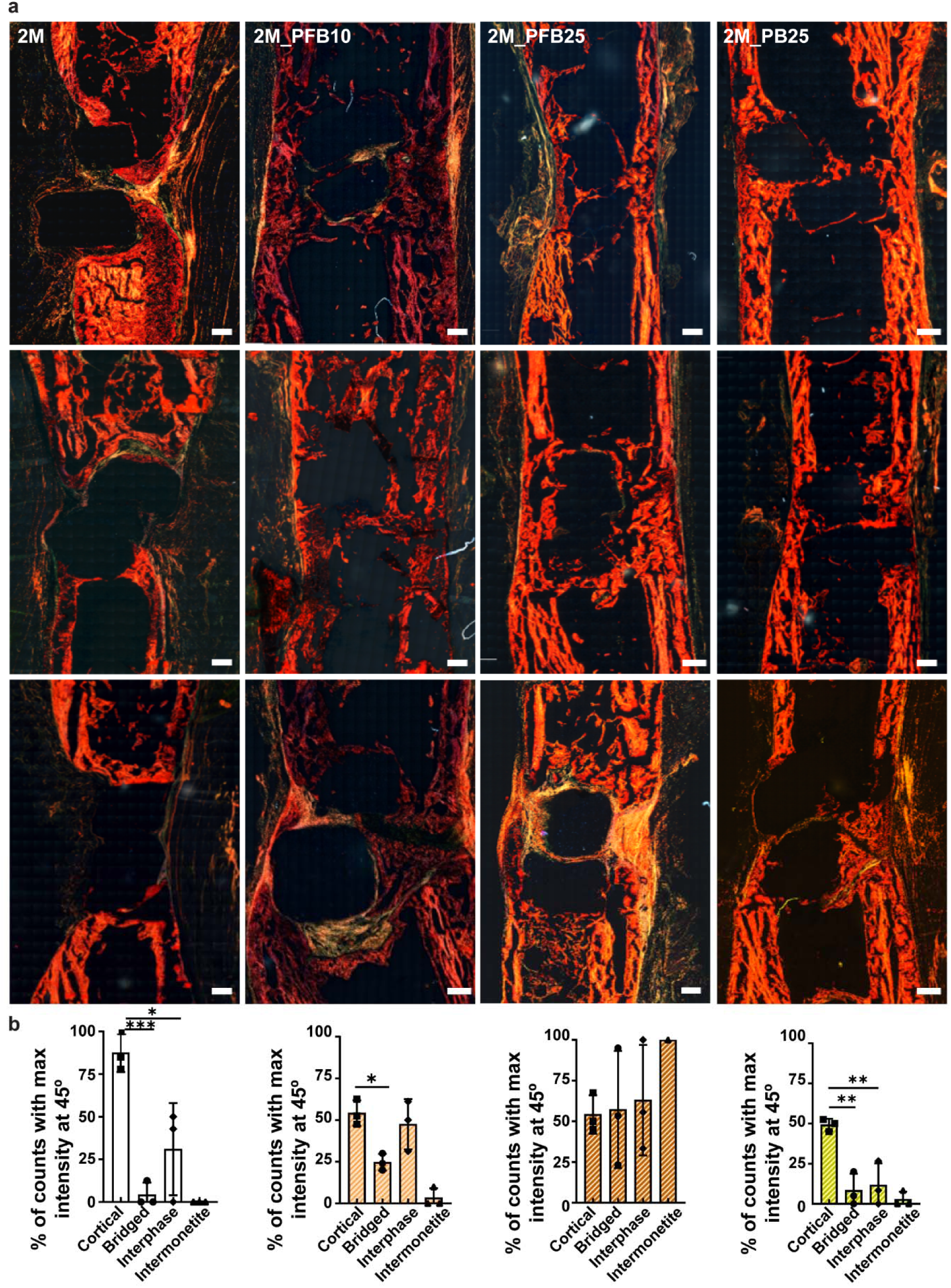
Collagen fibre characterization in the newly formed bone. a) Collagen fibre organization in the critical-sized bone defects shown by alcian blue and picrosirius red staining imaged with polarized light microscopy. Images represent merge of six polarizer-analyser angle combinations (polarizer-analyser: 0-45°, 15-60°, 30-75°, 45-90°, 60-105° and 75-120°), for n = 3 samples in each group. Scale bar: 500 μm. b) Preferential orientation of collagen fibres indicated by the percentage of measurements (counts) with maximum intensity at polarization light angle of 45°, for n = 3 samples per group, four ROIs (cortical, bridged, interphase, intermonetite) and n > 6 measurements per ROI. Symbols represent the mean value of all the measurements per region and the bars, the standard deviation. One-way ANOVA with Bonferroni post-hoc test was performed between regions per group. Statistical significance is indicated with *p<0.05, **p<0.01, ***p<0.001.

The quantitative analysis of collagen fibre orientation was performed as previously described ^[11]^ and is summarized in Figure 5b. The maximum intensity of collagen fibres in the cortical region of all samples occurred when the polarizer was set at 45° and the analyser at 90°, indicating that collagen fibres in native cortical bone are predominantly oriented parallel to the main bone axis (Figure S11–S16). Next, we quantified collagen fibre alignment in the bridged region and compared it with the cortical region. The 2M_PFB25 group exhibited comparable collagen fibre orientation between the cortical and bridged regions (Figure 5b, S14–S16). However, in the groups 2M_PFB10 and 2M_PB25, no predominant collagen fibre orientation parallel to the main bone axis was observed. This reflects statistically significant differences in collagen organization between the native cortical bone and bridged bone in these groups, with the highest level of maturation in the 2M_PFB25 group.

Regarding the interphase region, both the 2M_PFB10 and 2M_PFB25 groups exhibited collagen fibre orientation comparable to cortical bone. In contrast, the 2M_PB25 group showed statistically significant differences between the interphase and cortical regions (Figure 5b). This reflects a higher level of maturation in the 2M_PFB10 and 2M_PFB25 groups for the bone grown in direct contact with the monetite. Intermonetite regions were measured in too few cases to reach the minimum sample size for statistical analysis in all groups.

### 2.8. PEA-coated monetites functionalized with FN and low-dose rhBMP-2 show new bone with a maturation quality trending toward that of cortical bone

SAXS measurements provided additional information on the sub-micron bone organisation, providing values for the average mineral particle thickness (T) within the volume of the X-ray beam, average predominant orientation (direction of the lines in Figure 6a), and degree of alignment (ρ) (length of the lines), where ρ = 1 represents perfect alignment and ρ = 0, random orientation of mineral particles, as outlined by others^[41,42]^ . Results were analysed in the same ROIs as those defined for the polarized light analysis in Figure 5, adding one further region: the bone infiltrations inside the monetite (Figure S10).

**Figure 6.**
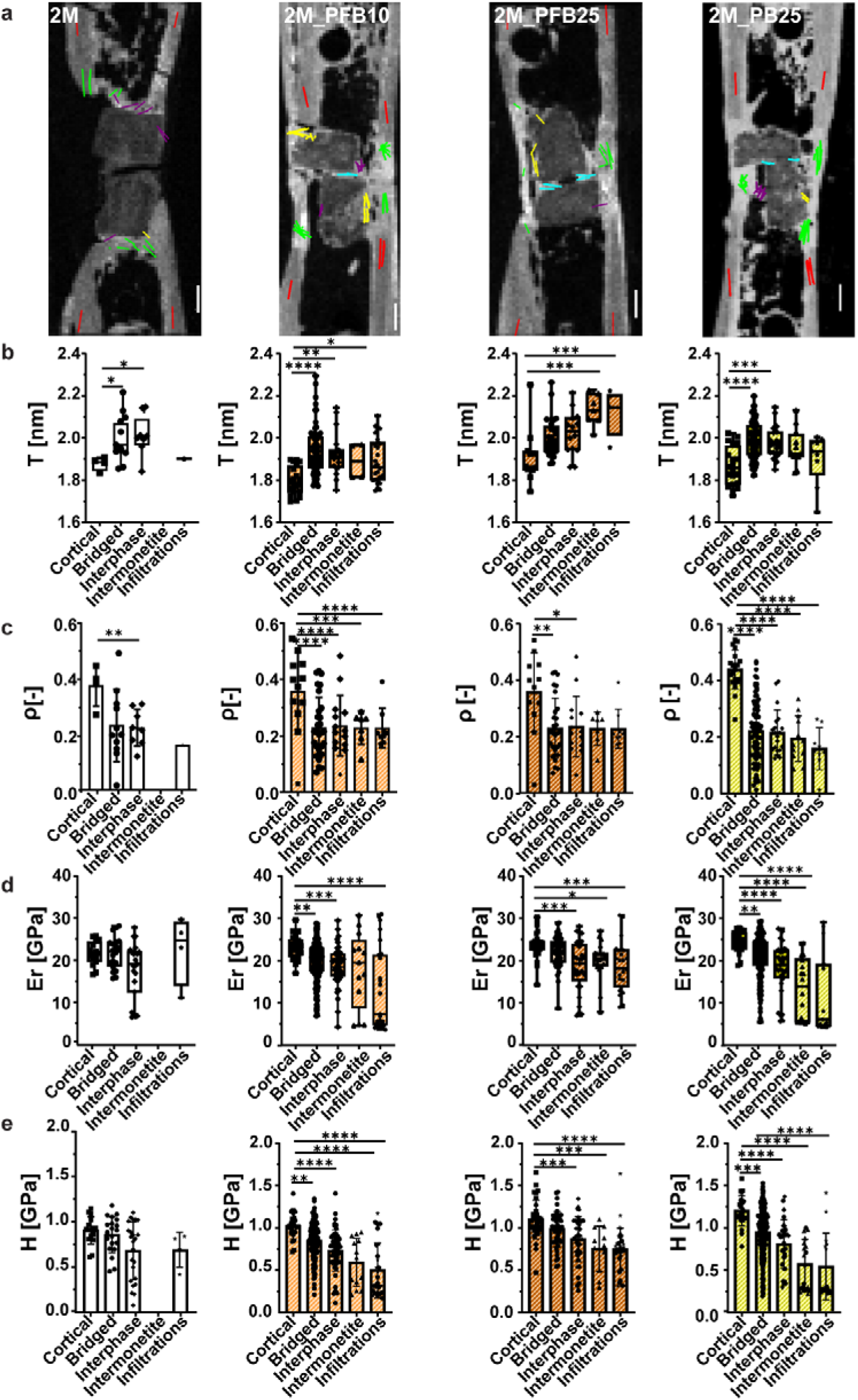
Bone mineral organisationa) SAXS results overlaid on nanography images of one representative sample per group, including 2M, 2M_PFB10, 2M_PFB25 and 2M_PB25. The colour of the lines represents the analysed ROI (cortical = red, bridged = green, interphase = purple, intermonetite = cyan and infiltrations = yellow). The direction of the lines depicts the average predominant orientation of mineral particles and the length of the lines shows the degree of alignment, ρ. Scale bar: ρ = 0.5. Quantified values of (b) mineral particle thickness, T, (c) degree of alignment, ρ, (d) local elastic indentation modulus, E_r_, and (e) hardness, H, shown per region and experimental group. Bar plots indicate the mean and standard deviation of all measurements per region, where every dot represents one SAXS or nanoindentation measurement. Boxplots show the median value and the interquartile range per region. One-way ANOVA with Bonferroni post-hoc test (bar plots) or its equivalent non-parametric Kruskal-Wallis test (boxplots) was performed to compare ROIs of each group. Statistical significance is shown with *p<0.05, **p<0.01, ***p<0.001, ****p<0.0001.

SAXS results were displayed graphically, overlaid on transmission maps. The line colour represents the region analysed (cortical = red, bridged = green, interphase = purple, intermonetite = cyan, and infiltrations = yellow). All analysed samples are shown in Figure S17–S19. The mineral particle thickness in the bridged region of the 2M_PFB25 group did not differ significantly from the native cortical bone, suggesting a more advanced state of new bone maturation (Figure 6b). However, all other measurements in new bone regions showed statistically significantly higher particle thickness (T) than the native cortical bone control, suggesting a less mature mineralized tissue (Figure 6b). On the other hand, the degree of alignment of the particles (ρ) was always lower in new bone regions (Figure 6c), indicating less organised new bone tissue and corroborating the results obtained with the picrosirius red staining and quantitative polarized light imaging. Sample to sample variability is shown in Figure S17 and S19.

### 2.9. PEA-coated monetites functionalized with FN and low-dose rhBMP-2 show elastic indentation modulus and hardness comparable to cortical bone in newly formed bridged bone regions

Nanoindentation measurements were performed in the same selected groups, samples and ROIs. Nanoindentation measurements (Figure 6 d, e) provided the local elastic indentation modulus (Er) and the hardness (H) of each region. Both the elastic modulus and hardness of the bone in the bridged region of the 2M_PFB25 group were not statistically distinguishable from native cortical bone. However, in all other measured groups the elastic indentation modulus and hardness were lower in all regions compared to the original cortical bone. This confirms the results of higher bone maturation quality of 2M_PFB25 shown with quantitative polarized light imaging and SAXS. Sample to sample variability is shown in Figure S20 and S21.

### 2.10. PEA-coated monetites functionalized with FN and low-dose rhBMP-2 favour monetite degradation and bone integration in an ovine defect model

Finally, the efficacy of the selected 2M_PFB25 group, which showed the most advanced tissue maturation in the rat model, was assessed in a large-animal model. We evaluated in vivo monetite degradation and new bone formation in the vicinity of the monetite using synchrotron phase-contrast microCT at SOLEIL (Anatomix beamline). The coherent radiation of synchrotron microCT allows the acquisition of high resolution (5.4 μm voxel size) scans of large sheep bone explants, with high signal-to-noise ratio and improved contrast in grey values, needed to separate the ceramic biomaterial from the new bone. Two experimental groups were compared: the previously mentioned PFB25 and PEA+FN without rhBMP-2 (named PF) as control (Figure 7a), as its efficacy was demonstrated previously ^[26]^. The ovine defect model comprised an 8 mm diameter monocortical defect in the knee of the sheep (Figure 7b,c), into which eight monetites were implanted randomly (Figure 7d-f). Image segmentation showed the monetites located in different anatomical locations, which we defined as periosteal region (periost), cortical, endosteal (endost) and bone marrow (marrow) (Figure 7g,h). Therefore, all quantitative analyses of the monetite volume (Figure 7i), the bone volume (Figure 7j) located in the region of interest (BV/TV, Figure 7k) and bone infiltrations (Figure 7l) were performed taking into account not only the experimental group but also the anatomical location. Only monetites defined as periost, cortical and endost, which had mineralized bone in their surroundings were considered. The monetite classification performed by anatomical location in every sample can be found in Figures S22–S32.

**Figure 7.**
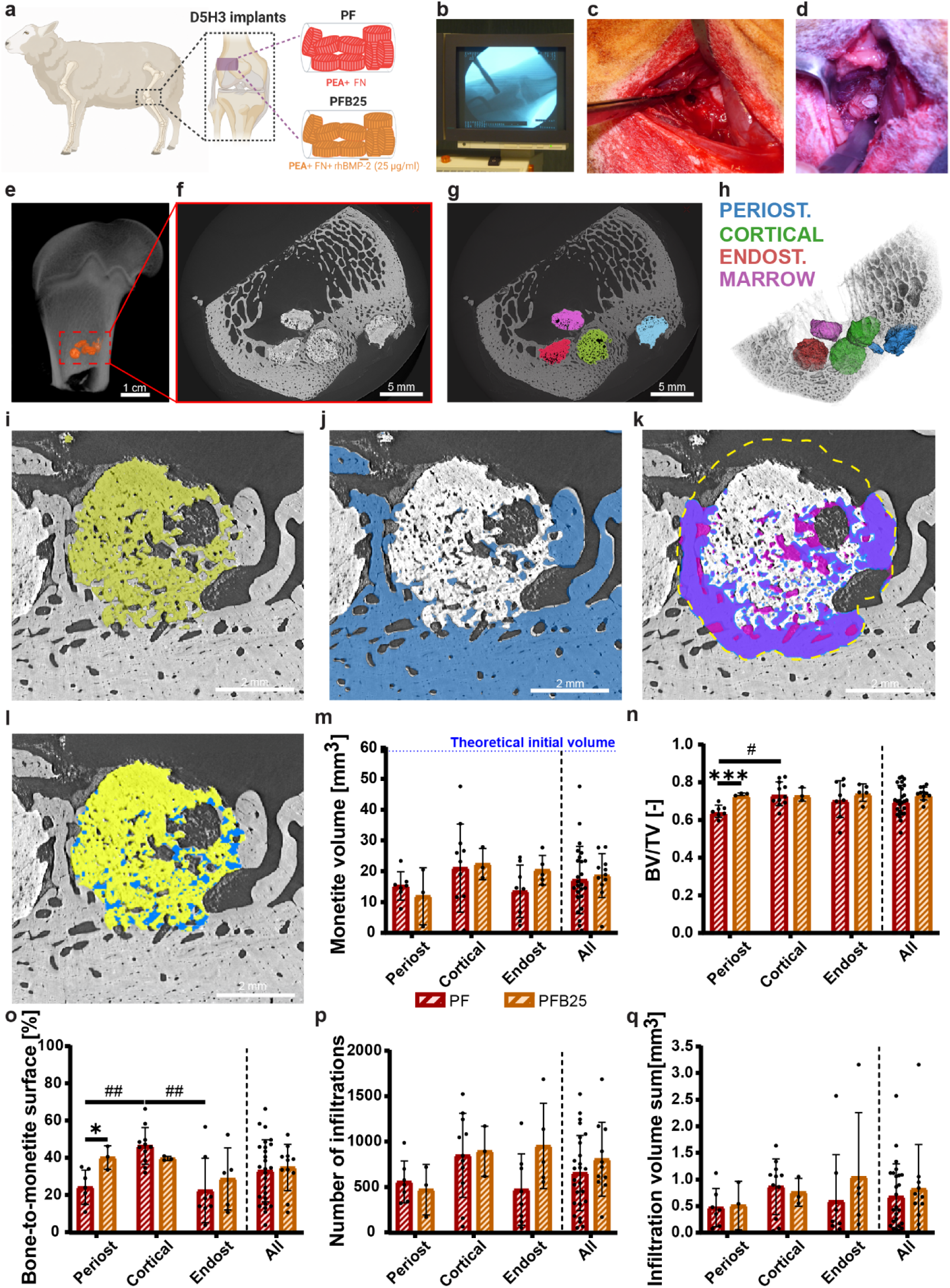
Ovine bone defect model. a) Bone defect performed in a large-animal ovine model, with a cylinder drilled in the knee and filled with eight monetites; two different monetite combinations used, PF and PFB25. Figure created with BioRender.com. Monetites were implanted following the next steps: b) X-ray images of the sheep knee to locate the defect; c) generating the defect in the sheep knee with the drill and removing debris from the defect and finally d) placing the monetites in the defect. After 3 months, sheep were sacrificed and explants were analysed. e) 3D reconstruction of X-ray images of the whole femoral head of a sheep to show the anatomical location of all monetites. Scale bar: 1 cm. f) Synchrotron phase-contrast microCT image with g) segmentation of the monetites by anatomical location. Scale bar: 5 mm. h) Colour code used to classify monetites per anatomical location in periost (blue), cortical (green), endost (red) and marrow (pink). Synchrotron phase-contrast microCT image showing i) monetite volume segmentation, j) bone volume segmentation, k) bone volume (purple) per total volume analysed (pink) in the monetite dilated region (dashed yellow line) and l) monetite volume in yellow with bone infiltrations in blue. Scale bar: 2 mm. Quantification per group and anatomical location of m) monetite volume, n) bone volume per total volume, o) bone-to-monetite surface percentage, p) number of infiltrations and q) sum of the volume of all infiltrations per monetite. Bar plots indicate the mean and standard deviation of all measurements per group and region, where every dot represents one monetite. Comparisons between groups were performed with t-test (statistical significance shown with *p<0.05, ***p<0.001). Comparisons between different anatomical locations of each group were performed with one-way ANOVA with Bonferroni post-hoc test (statistical significance shown with #p<0.05, ##p<0.01). n = 7 animals in PF (7 monetites in periosteal region, 11 monetites in cortical region and 8 monetites in endosteal region) and n = 4 animals in PFB25 (3 monetites in periosteal region, 3 monetites in cortical region and 5 monetites in endosteal region).

After successful segmentation of the ceramic monetite and the new bone with a custom-made 3D U-Net model (available on Zenodo: 10.5281/zenodo.23324947, Supplementary Video S1), the monetite volume was quantified in each region and experimental group (Figure 7m). Considering a theoretical initial monetite volume of around 59 mm^3^, all monetites lost at least 50% of their volume, with no statistically significant differences between both experimental groups or anatomical locations, suggesting similar remodelling in the large-animal model, which is greater than that observed in the small-animal model.

Next, the bone volume in the vicinity of the monetite was quantified. For this analysis, the monetite volume was dilated by 500 μm, as described in the Methods. BV/TV was slightly higher in the PFB25 group at all anatomical locations, and the difference reached statistical significance in the periosteal region (Figure 7n). In addition, monetites without rhBMP-2 were impacted more by anatomical location than those functionalised with the growth factor.

The influence of anatomical location was likewise apparent in the PF group when new bone integration adjacent to the monetites was analysed (percentage of monetite surface covered by bone), but not in the PFB25 group (Figure 7o). Bone-to-monetite contact was generally higher in the PFB25 group, significantly in the periosteal region: these monetites were in direct contact with bone over nearly 40% of their surface, indicating that a low-dose of rhBMP-2 is sufficient to promote integration.

Finally, bone integration within the monetites was assessed from the number and volume of bone infiltrations (Figure 7p,q). Both the total number of infiltrations through the monetites and their combined volume were higher in the PFB25 group than in the PF group, indicating increased bone ingrowth with a low-dose of rhBMP-2 added to the implant.

## 3. Discussion

In this study, we investigated PEA-coated degradable monetites functionalized with FN and low-dose rhBMP-2 as a bioactive bone graft substitute, demonstrating its efficacy through a multiscale and multimodal characterization. Our results show that the monetites serve as a resorbable, osteoconductive scaffold that supports coupled osteoclastic resorption and new bone formation, enabling regeneration in critical-sized bone defect models. While monetite alone provides a permissive substrate for bone ingrowth, coating and functionalizing it with a bioactive nanoscale interface such as PEA+FN+rhBMP-2 substantially increases both the extent and the structural maturity of the regenerated bone.

A central finding is that monetite behaves as a highly resorbable calcium phosphate whose degradation appears closely linked to cellular remodelling as supported by the presence of osteoclasts and coupled bone ingrowth supporting a cellular contribution alongside material chemical dissolution ^[13,43–48]^. This is consistent with previous reports of monetite as a resorbable calcium phosphate that supports coupled osteoclastic degradation and new bone formation in preclinical and clinical settings, making it an attractive base material for bone regeneration. In our study, this intrinsic resorbability provided a permissive base material, but PEA coating combined with FN+rhBMP-2 functionalization clearly increased the extent of regeneration, indicating that the coating does not replace the osteogenic potential of monetite; instead, it amplifies it.

In addition, our data demonstrate that PEA+FN+rhBMP-2 coating not only enhances the regenerative process, but also enables robust bone repair at low BMP-2 doses, which is a major translational advantage given the known limitations of high-dose BMP therapies ^[26,27,49,50]^. This extends those earlier works by showing that a bioactive coating improves the structural quality of the regenerated bone, and that the benefit is retained in a large-animal model. In fact, the nanoscale organization of FN and BMP-2 on PEA enables exposure of integrin- and growth-factor-binding domains, fostering synergistic signalling through integrins and BMP receptors ^[29,30,33]^. This mechanistic synergy likely lowers the dose of BMP-2 required for effective bone repair while maintaining, or even enhancing, the quality of bone formed.

PRP was evaluated as a clinically relevant, autologous alternative to recombinant FN because its complex mixture of adhesive proteins and growth factors can support cell attachment, proliferation, angiogenesis and tissue repair ^[37–39]^. Although PRP formed a network-like organization on monetites, retained rhBMP-2, and showed osteogenic activity at levels comparable to FN in vitro, PRP-based functionalization generally failed to promote defect bridging and significantly reduced new bone formation in vivo. Only the combination of PRP, PEA and 25 μg/ml rhBMP-2 improved bridging and local bone formation, suggesting that PRP alone did not provide sufficiently stable or bioactive rhBMP-2 presentation. Consequently, PRP was not considered a reliable substitute for FN and was excluded from the subsequent multiscale tissue evaluation.

Our histological and structural findings indicate that the coating also influences how new matrix is organized. Collagen architecture strongly determines mineral deposition, matrix anisotropy, and ultimately the mechanical performance of bone, because the hierarchical arrangement of collagen fibrils templates apatite deposition and contributes to osseointegration ^[51–53]^. In this context, our observation that bridged regions in the PEA-coated group functionalized with FN+rhBMP-2 exhibited a more organized collagen network comparable to native cortical bone is particularly relevant: it indicates not just more bone, but bone that is structurally more mature and more closely resembles native cortical bone. This is important because bone quality depends not only on quantity, but also on the alignment of collagen and mineral at the nanoscale ^[7,54]^.

SAXS analysis strengthens this conclusion by providing direct ultrastructural indicators of maturation. SAXS provides complementary nanoscale information on the mineral phase, resolving nanoscale features of mineralized collagen, including fibril orientation and mineral crystal organization, and therefore adds a level of structural insight that conventional histology cannot provide. Animals implanted with monetites coated with PEA+FN+25 µg/ml rhBMP-2 showed no statistically significant difference between the mineral particle thickness of native cortical bone and that measured in the bridged and interface regions. This, together with the observations of collagen organisation suggested a more advanced state of new bone maturation in this experimental group. The presence of thicker mineral particles but lower degree of alignment than native cortical bone in all regions is consistent with the expected developmental sequence of bone mineralization ^[55,56]^. However, the slightly higher degree of mineral alignment observed in PEA+FN+rhBMP-2 compared to other groups suggests more advanced progression toward an organized lamellar matrix, rather than simply inducing rapid but disordered mineral deposition ^[7,54,57,58]^. This distinction matters because biomaterials that favour fast but poorly organized mineralization may fail mechanically over time, whereas materials that support hierarchical maturation are more likely to produce robust bone repair ^[59]^.

In this context, our mechanical testing provides further evidence of more advanced maturation. Nanoindentation studies of bone have repeatedly shown that local stiffness and hardness correlate with mineral content and microstructural organization, while newly formed bone generally exhibits lower values until it becomes more mineralized and better aligned ^[7,54,59,60]^. Accordingly, the improved nanoindentation response in the PEA+FN+25 µg/ml rhBMP-2 group indicates that the regenerated tissue was not only more abundant, but also mechanically more competent at the 10-week endpoint: both elastic indentation modulus and hardness of the newly formed bridged region statistically resembled the ones of native cortical bone. This is a key translational point, because defect repair must ultimately restore load-bearing function, not merely fill space with mineralized tissue ^[61–63]^.

From a translational perspective, these results position PEA-coated monetites functionalized with FN and rhBMP-2 as a platform that combines three properties: osteoconductive resorption, nanoscale presentation of the growth factor, and matrix-guided regeneration. The sheep study reported here shows that these monetites support bone formation in large defects while themselves being resorbed. Synchrotron phase-contrast microCT allowed ceramic biomaterial degradation and new bone ingrowth to be assessed at a resolution that, to our knowledge, has not previously been applied to a large-animal defect model, and at length scales relevant to the clinical setting. Taken together, these data indicate that the PEA+FN+rhBMP-2 coating promotes regeneration and also influences how the newly formed tissue matures.

Overall, our findings show that monetite is a favourable resorbable calcium phosphate, and that its regenerative performance is substantially improved by a PEA+FN+rhBMP-2 coating that organises growth-factor presentation and guides maturation of the mineralised matrix. The combination of improved collagen alignment, more mature mineral organisation and near-physiological local stiffness suggests that the platform improves not only the extent of repair but also the quality of the bone formed. The large-animal data support further development of the approach for large bone defects and represent a step towards clinical application.

## 4. Conclusions

This work establishes the efficacy of degradable PEA-coated ceramic monetites functionalised with FN and low-dose rhBMP-2 for bone regeneration based on a multiscale analysis using microcomputed tomography, backscattered electron microscopy, histomorphometry, quantitative polarised light microscopy, SAXS and nanoindentation, and synchrotron phase-contrast microCT,. MicroCT and 3D reconstruction showed that only monetites functionalised with PEA and rhBMP-2 bridged the defect, with a significant increase in new bone volume in both the monetite vicinity and the whole defect volume. In contrast, PRP functionalisation significantly reduced bone formation in both regions, showing that the choice of osteogenic cue matters. Monetite degradation and the associated increase in interconnected porosity favoured bone ingrowth and produced a continuous interface between monetite and new bone. The multiscale analyses show that new bone in the 2M_PFB25 group — monetites coated with PEA and functionalised with FN and 25 μg/ml of rhBMP-2 — was structurally and mechanically more mature. Quantitative polarised light microscopy showed that only this group recapitulated the preferential collagen fibre orientation of the original cortical bone across the bridged and interphase regions, indicating better guidance of matrix organisation at the tissue level. SAXS measurements indicated improved mineral alignment, and therefore a more advanced state of hierarchical organisation, consistent with the collagen data — although all regenerated regions still contained thicker and less aligned mineral particles than cortical bone. Finally, nanoindentation showed that the elastic modulus of the bridged region was indistinguishable from that of native cortical bone. Together, these findings show the formation not only of more bone, but of more mature bone, with a hierarchical organisation and local mechanical properties approaching those of native cortical tissue. Synchrotron microCT analysis in an ovine model confirmed these findings at a clinically relevant scale, supporting the translational potential of the system. Taken together, these findings position PEA-coated monetites functionalised with FN and low-dose rhBMP-2 as a strategy for the structural and functional regeneration of large bone defects.

## 5. Experimental section

### Fabrication, functionalization and characterization of monetites

Monetites were produced using an established manufacturing process developed by Histocell S.L. Briefly, brushite cement was first generated by combining β-tricalcium phosphate (β-TCP), monocalcium phosphate monohydrate, calcium carbonate and sodium pyrophosphate in an acidic aqueous solution undergoing the following reaction:

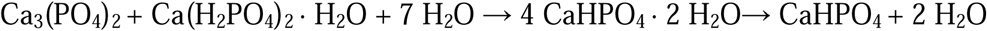

During the setting reaction in a defined silicone mould, the carbonate reacts with the hydrogen ions in the medium, decomposing into carbon dioxide, forming bubbles which, after setting, generate a spongy and porous brushite matrix. After hardening in the moulds with the desired geometry (5x3mm or 2.5x1.5mm, diameter-height), the brushite granules were neutralized by washing with purified water until neutral pH was reached. Conversion of brushite to monetite was subsequently carried out by dry thermal treatment at 121 °C in an autoclave. The resulting monetites exhibited an amorphous matrix with limited macroporosity and an irregular pore distribution.

For the PEA-coated experimental groups, monetites were functionalised by plasma polymerisation deposition of ethyl acrylate (EA) monomers onto the monetite. Polymerisation was carried out following a protocol defined by the Centre for the Cellular Microenvironment of the University of Glasgow and Histocell S.L. using a commercial custom designed PECVD system (PICO, Diener, Germany), applying 21 W of power and 0.2 mbar of pressure. To ensure homogeneous coating, the monetites were processed in a rotating vessel during polymerisation. Following coating, the PEA-functionalised monetites were sterilised using ethylene oxide before use in both in vitro and in vivo studies.

Surface chemical composition of the monetites was analysed by XPS using a Thermo Scientific Nexsa XPS system equipped with a monochromated Al Kα X-ray source (1486.7 eV). Spectra were acquired from the top, bottom and curved surfaces of the monetites to assess the uniformity of the PEA coating. Survey spectra were analysed using CasaXPS software (v2.3.19PR1.0). The presence of the PEA coating was confirmed by changes in the surface elemental composition, with particular attention given to carbon (C 1s), oxygen (O 1s), calcium (Ca 2p) and phosphorus (P 2p) signals.

Finally, prior in vivo implantation, monetite shape and porosity were characterized using a micro-computed tomography (microCT) RX Solutions X-ray microCT Easy Tom 160 system equipped with a nano-focus X-ray tube and a horizontal flat-panel detector. An aluminium filter was applied during image acquisition. Scans were acquired in helicoidal mode (8 turns) at 80 kV and 69 µA (5.5 W) in large focal spot mode, at a voxel size of 2 µm resolution. Images were reconstructed and 3D renderings generated using ORS Dragonfly version 2022.2 software (Object Research Systems Inc, Montreal, CA) for quantification of monetite volume and porosity.

Following PEA coating, monetites were functionalised with FN (R&D Systems) and rhBMP-2 (R&D Systems). Monetites were first immersed in FN solution (20 µg/mL) under vacuum for 1 h at room temperature to promote infiltration of the protein solution throughout the monetite structure. After FN adsorption, monetites were washed with phosphate-buffered saline (PBS) and subsequently incubated with rhBMP-2 solution (100 ng/mL) using the same adsorption procedure. The amount of adsorbed FN was determined indirectly by measuring the non-adsorbed FN remaining in the supernatant using a micro bicinchoninic acid (BCA) protein assay. Absorbance was measured at 562 nm and FN concentration was calculated from a standard curve prepared using known FN concentrations. rhBMP-2 adsorption was quantified by measuring the residual rhBMP-2 concentration in the supernatant using a sandwich ELISA kit (R&D Systems), according to the manufacturer’s instructions. Absorbance was measured at 450 nm with wavelength correction at 570 nm, and rhBMP-2 concentration was calculated using a four-parameter logistic standard curve. For both FN and rhBMP-2, the amount of adsorbed protein was calculated by subtracting the amount remaining in the supernatant from the initial amount added to each monetite.

PRP was prepared from four healthy donors using the Endoret® kit (BTI, Vitoria, Spain). Liquid PRP was activated with 10% calcium chloride (CaCl ), frozen at −80 °C and thawed prior to use. Samples were subsequently centrifuged at 16,000 x g to remove platelet debris. Monetites were then incubated with PRP following the same protocol used for fibronectin coating. Depending on the experimental group, PRP-functionalised monetites were further combined with rhBMP-2 and/or PEA coating as described before.

To assess protein organisation on PEA surfaces, FN and PRP coated PEA-functionalised glass coverslips were analysed by atomic force microscopy (AFM). Samples were washed, dried under nitrogen and imaged in tapping mode using a NanoWizard AFM system (JPK Instruments, Germany). Height images were acquired to evaluate the nanoscale organisation of adsorbed proteins.

### In vitro validation of coatings

Human adipose-derived stromal cells (hASCs) were provided by Histocell S.L. and used for all in vitro experiments. Cells were maintained in Dulbecco’s Modified Eagle Medium (DMEM) supplemented with 10% foetal bovine serum (FBS), 1% sodium pyruvate, 1% non-essential amino acids and 2% antibiotic solution (6.74 U mL□¹ penicillin-streptomycin, 0.2 μg mL□¹ fungizone and 2 mM L-glutamine). Cells were cultured at 37 °C in a humidified atmosphere containing 5% CO and used between passages 1 and 7.

Monetites were placed in non-tissue culture treated 48 well plates prior to cell seeding. 150000 cells were seeded onto each monetite in serum-free culture medium using a reduced medium volume to promote cell attachment. After 1.5 hours, an equal volume of complete medium containing 10% FBS was added, resulting in a final serum concentration of 5% FBS. Following 24 hours of culture, the medium was replaced with complete medium containing 10% FBS for the remainder of the experiment. Culture medium was refreshed twice weekly. For osteogenic differentiation studies, OM was used as positive control. OM was prepared by supplementing DMEM with 100 μM ascorbic acid, 100 nM dexamethasone and 3 mM sodium phosphate. All in vitro validation experiments were performed using the larger monetites (5 × 3 mm, diameter × height).

Cell viability on the monetites was assessed after 3 and 10 days of culture using a LIVE/DEAD Viability/Cytotoxicity Kit (Life Technologies, UK) according to the manufacturer’s instructions. Samples were imaged using fluorescence microscopy and qualitative assessment of cell viability was performed based on the distribution of live and dead cells on the monetite surface.

Osteogenic differentiation of hASCs cultured on the monetites was evaluated by quantitative real-time PCR (qRT-PCR). Total RNA was extracted using the RNeasy Micro Kit (Qiagen), including an on-column DNase treatment, according to the manufacturer’s instructions. RNA concentration was quantified and normalised prior to cDNA synthesis using the QuantiTect Reverse Transcription Kit (Qiagen). Gene expression analysis was performed using the QuantiFast SYBR Green PCR Kit (Qiagen) and gene-specific primers (Eurofins Genomics, Germany). Amplification was carried out using an Applied Biosystems 7500 Real-Time PCR System. Gene expression levels were normalised to *GAPDH* and relative expression was calculated using the 2 ΔΔCt method. Primers used for qRT-PCR can be found in Table S1.

### Rat critical-sized bone defect model

All experiments involving animals were approved by the Ethics Committee for Animal Experimentation of the University of Navarra (Comité de Ética para la Experimentación Animal, CEEA) and Navarra regional Government, CEEA E55-22(074-20). A 5-mm critical-sized defect was created in the femora of 10-12 weeks-old female Sprague-Dawley rats as previously reported ^[29]^. The void was filled either with one monetite (5 x 3 mm, diameter-height) or two (2.5 x 1.5 mm). Biodegradable plain or PEA-coated monetites were subjected to different surface treatments depending on the experimental group (Figure 1, Figure S1). These surface treatments included 20 μg/ml human fibronectin (FN, 1918-FN-02M, R&D systems, Minneapolis, MN), rhBMP-2 (InductOs 1,5 mg/ml, Medtronic, Dublin, Ireland) in a concentration of 10 or 25 µg/ml and PRP, following the procedure described in the section before. To avoid possible disparities derived from donors, the PRP from different donors was pooled and monetites were treated with the mix.

A total of 70 animals divided in 12 in vivo experimental groups (Figure 1a), including the plain monetite groups (n = 5-7 animals per experimental group) were analysed within this critical-sized bone defect model.

Ten weeks after implantation, rats were sacrificed and femora were fixed in 4% PFA (15713, Electron Microscopy Science, EMS, Morgantown, PA) to perform a multiscale characterization of non-decalcified bone specimens embedded in poly(methyl methacrylate) (PMMA, Technovit 9100, Morphisto GmbH, Germany) as described previously ^[40]^. Briefly, samples were processed for cold poly-methyl methacrylate (PMMA) embedding using the manufactureŕs protocol (Technovit 9100, Morphisto GmbH, Germany). Explants were dehydrated through an ascending ethanol series and stained with rhodamine solution, 0.417 g of Rhodamine B (Fisher Scientific SL, Spain) per 100 mL of ethanol. Following staining, the samples were cleared with xylene and subsequently embedded in polymerized PMMA.

### Ovine critical-sized bone defect model

All procedures were performed in accordance with institutional guidelines, with ethical approval, in accordance with the European regulations for experimental animals’ welfare. This study was approved by the University of Navarra Ethics Committee with reference code E40-24(074-20E3). Twelve female six-months-old sheep (Ovis aries), with average weight of 35 ± 5 kg were randomly assigned to two treatment groups PEA-coated monetites with 20 μg/ml of FN as the control on the one side (named PF) and PEA-coated monetites with FN and rhBMP-2 (25 μg/ml) as the experimental group (named PFB25) (Figure 7).

Sheep were group housed in a barn and had access to food and water for the duration of the study. Animals were examined to ensure good physical condition and were acclimatized to the indoor barn environment for a minimum of 14 days prior to surgery. Food was withheld for a minimum of 12 hours, and water was withheld for ∼ 2 hours prior to surgery to reduce the risk of regurgitation and aspiration. One hour prior to surgery, 12.5 µg/kg dexmedetomidina (Dexdomitor 0.5 mg/ml. Zoetis) was applied intramuscular in order to sedate the animals and transferred to the surgical facilities. The skin of the lateral antebrachium of one forelimb was shaved, cleaned and de-greased prior to application of Fentanyl cutaneous patches to deliver approximately 2.5 µg/kg/h Fentanyl (DUROGESIC MATRIX 75 µg/h, Janssen-Cilag, S. A. CN 848119) held by a bandage layer (Easifix® Cohesive, Leukoplast®) to provide pre-operative and up to 3 days post-operative analgesia, and 20 gauge intravenous catheter (BD Insyte, CN 381037) was placed in the contralateral forelimb. Prior induction, preoperative antibiotic therapy was administered, 22 mg/kg Cefazoline (Cefazolina 1G Normon, CN 769125.9). Intravenous induction of anaesthesia using 4-5 mg/kg Propofol (Propofol lipuro 10 mg/ml. BBRAUN. CN 855437.9 OH) and 0.05 mg fentanyl (Fentanest 0.05 mg/ml. Kern Pharma. CN 756650.2H) was performed, the solution was allowed to take action for one minute and intubation was performed. Anaesthesia was maintained with 0.3-0.6 mg/ml/min Propofol (Propofol lipuro 20 mg/ml, BBRAUN. CN 697791). Pre- and 48 h post-operative anti-inflammatory treatment was given by means of 4 mg/kg Carprofen (Rimadyl™ 50 mg/mL, Zoetis. CN 572100.2) subcutaneus injection. Prophylactic antibiotics therapy (Pen and Strep Suspension for Injection, 0.04 ml/kg, Exabiopen 250 ml, SYVA. CN 582615) was given post-operatively the following ten days. For surgery, the animals were placed in lateral recumbency to access the lateral aspect of selected hind leg. The stifle area was then clipped and prepared using chlorhexidine scrub solution (Hibiscrub 4% 500 ml, Mölnlycke) and povidone iodine (Povidone iodine 10%. Acofarma). The skin and underlying soft tissue over the lateral femoral condyle were incised and a single 8 mm diameter cylindrical drill hole/defect was created in the cancellous bone region of the lateral femoral condyle, with a 15 mm depth in the metaphysis. The defect was irrigated with sterile saline to remove any bone fragments and dried using a sterile swab to check for bleeding. The incision was filled with 8 monetites (Figure 7). For implantation, the tip of the delivery syringe was brought into position and its contents were deposited. After that, the incision was sutured in layers using absorbable suture material (Novosyn size 2/0, BBRAUN, C0068242). Hypoclorine hydro-gel (Hypoclorine Skin Care. JTPHARMA, CN 04300084) was then applied over the wound for protection from contamination. After surgery, animals were housed in habitual pen with the other sheep. Daily grimace and mobility scoring was performed to monitor for any signs of pain or complications, rescue analgesia (0.01 mg/kg IM Buprenorphine. Buprecare multidosis 0.3 mg/ml. Ecuphar, CN 578950) was administered to two animals and another sheep was humanely euthanized upon reaching an endpoint criteria. Animals were sacrificed after 13 weeks post-operatively by intravenous barbiturate (Tiobarbital 0.5G BBRAUN, CN 635573) overdose. The femoral condyles were retrieved immediately and fixed.

### High resolution laboratory microcomputed tomography (microCT) of rat explants

All explants embedded in PMMA were scanned using a RX Solutions X-ray microCT Easy Tom 160 system equipped with a micro-focus X-ray tube and a vertical flat-panel detector. A two-step scanning strategy was used with the 70 samples. First, a low-resolution (24 µm voxel size) scan was acquired to obtain an overview of the entire femur and to identify the defect region. For this overview scan, tomography was carried out in step-by-step mode, and the X-ray source was operated at 100 kV and 200 µA (20 W) in middle focal spot mode. The acquisition was performed with a frame rate of 5 and 4 averaged frames per projection, and an aluminium filter was used. Projection images were reconstructed using filtered back projection (tomography mode) with one iteration, a Tukey filter and sinus apodization.

Subsequently, a high-resolution (4.7 µm voxel size) scan was performed to analyse newly formed bone as described previously ^[40]^. Specimens were scanned at a at a voltage of 60 kV and a current of 166 µA (10 W) in small focal mode. Images were acquired with a frame rate of 1 and 7 averaged frames per projection, with an aluminium filter applied. Projection of images were reconstructed following the parameters established with the low-resolution scans.

Bridging of the samples was defined as a complete 3D closure of newly formed bone between the proximal and distal cortices across the defect, with no residual gap visible in the reconstructed volume. For non-bridged defects, the minimum gap distance between proximal and distal bone was measured as previously described ^[35]^ , by calculating the minimum straight line to join both bone parts. Newly formed bone volume (BV) was quantified both in (i) the entire 5 mm defect (defined as a 5 mm high cylinder covering the initial defect) and (ii) the immediate monetite vicinity (defined by morphological dilation of the segmented monetite volume by 10%). Next, the newly formed bone volume per each defined total volume (BV/TV) was calculated, using ORS Dragonfly version 2022.2 software. Deep-learning segmentation models were custom-made to automatically segment newly formed bone, monetite volume and porosity. Briefly, training ROIs were generated by manually labelling at least 15 representative 2D slices, which were used to train a U-Net model (100 epochs, stride ratio 0.25, batch size 512, Adadelta optimization^[64]^ with ORSDiceLoss used as the loss/validation metric). All segmentation models applied can be found in the following public repository [10.5281/zenodo.23324947]. In vivo monetite degradation and bone infiltration were quantified only for which the corresponding monetites had been scanned prior to implantation (n = 5-7 samples per experimental group).

### High resolution phase contrast synchrotron microCT, processing and analysis of ovine explants

Synchrotron microCT scans of sheep bone explants were acquired at the ANATOMIX beamline of the French synchrotron light source SOLEIL ^[65]^, under proposal #20250384. Previously fixed bone samples were scanned while immersed in 70% ethanol and mechanically stabilized to prevent movement artefacts. Scanning was conducted utilizing a polychromatic beam with a central photon energy of approximately 50 keV. The undulator gap was set to 6.1 mm, and the spectrum conditioned via a series of transmission filters including 0.6 mm of diamond, 26 µm of gold, and 1 mm of copper. Each tomographic scan comprised a total of 8000 radiographs spanning an angular range of 360° with an acquisition time per radiograph of 35 milliseconds. The distance between the sample and the detection plane, known as the propagation distance, was 1.9 m. The detector setup comprised a fluorescent screen (lutetium aluminum garnet, thickness 600 µm, supplier: Crytur, Czech Republic) coupled to a CMOS-based Orca Lightning camera (Hamamatsu Photonics, Japan) via a system of photo objectives in tandem geometry (Hasselblad, Gothenburg, Sweden). The set-up resulted in an isotropic pixel size of 5.4 µm. The camera was operated with a hardware region of interest (ROI) of 2944×2400 pixels (horizontal×vertical).

The tomographic volume data were reconstructed from the projection radiographs using the standard data processing pipeline of the beamline, with an inhouse-written Python script for pre-processing and the PyHST2 software (developed at ESRF, Grenoble, France ^[66]^ as the computing backend. A Paganin filter ^[67]^ as implemented in PyHST2, with a kernel width (“Paganin length”) of 40 pixels was applied in conjunction with the conventional filtered back-projection algorithm. The resulting reconstructed volumes are approximately 30.5 mm in diameter and 13.0 mm in height.

In cases where the relevant sample size exceeded the achieved field of view (FOV), individual datasets were fused resulting in one volume per sample. A precise segmentation separating mineralized bone tissue, monetite and background voxels was performed using a 3D U-net^[68]^ model implemented in Tensorflow/Keras. The network, of depth 5 (5 encoder stages, followed by a 4-stage decoding path) accepts a 3D sub-volume of 192x192x48 voxels as input, and was used to process full datasets as overlapping tilings of such subvolumes. Each stage in both the encoder and decoder consisted of two 3x3x3 convolutional layers with a dropout layer in between, followed by instance normalization. Max-pooling was used to in the encoder to reduce the data dimensions, and transposed convolution layers to increase the data dimensions in the decoder. The network was trained on manually segmented subvolumes containing target labels using Avizo Software (2025.1, Thermo Fisher Scientific, Waltham, MA, USA) and further refined based on corrected initial results. The final model is available at a public repository [10.5281/zenodo.23324947]. Segmentation masks were post-processed to remove artefacts. Individual monetites were manually classified according to experimental group and anatomical location in the bone at the periosteal or endosteal interface or within the cortical or bone marrow (periost., endost., cortical, marrow). Monetites cut during sample preparation or located outside of the mineralized bone compartment as well as monetite fragments were not included in the subsequent analysis.

Based on the post-processed masks, monetite volume and surface, and the bone-to-monetite surface (%) were calculated.

Additional masks were generated for the quantitative analysis. Monetite volumes were dilated by 96 voxels, approximately 500 µm. By intersecting with the bone label, bone volumes within each dilation were calculated (BV). To account for the different anatomical locations and the thereby naturally different amount of mineral tissue in the vicinity of the monetites, total volumes within each monetite dilation (TV) were created by closing the bone labelled volumes by 20 voxel and subtracting the monetite volumes. Finally, BV/TV was calculated within each monetite dilation. To assess the bone ingrowth into the monetites, monetite labels were closed by 10 voxels and ingrown bone was defined by intersecting the closed monetite with the bone mask. Individual infiltrations were size filtered with a minimum total size of 64 voxels.

Calculations were performed using IPSDK Explorer 4.2 (Reactiv’IP Smart Image Processing, Grenoble, France, 2026; https://www.reactivip.com/).

### Histology, histomorphometry and quantitative polarized light microscopy

In order to consistently expose the sagittal plate in all explants, a controlled-angle cutting procedure was used, guided by the low-resolution overview microCT scans, as described previously ^[40]^. Consecutive, numbered, longitudinal sections (6 μm in thickness) were cut from the PMMA-embedded blocks using a microtome (Model RM2255, Leica Biosystems). Serial histological sections were dried at 60 °C for 48 h and subjected to the following stains: Goldner’s Trichrome, Von Kossa + Movat Pentachrome, Tartrate-Resistant Acid Phosphatase (TRAP) and Alcian Blue + Picrosirius Red.

Stained slides were imaged using an AxioObserver 7 microscope (Zeiss) at 10x and 40x magnification. Imaging for all stains was performed using bright field microscopy in all samples per group (n = 5-7). Goldner’s Trichrome staining was used to visualize general tissue structure and morphology. The staining allows for the differentiations of collagen (green), osteoid (red), cell nuclei (black) and cell cytosol (bright red).

The Von Kossa + Movat Pentachrome staining was utilized to quantify mineralization and various connective tissue structures. Specific tissue components were differentiated as follows: mineralized cartilage and bone tissue (black), mineralization fronts (black, granulated appearance), collagen fibres (yellow), reticular fibres (light blue), muscle tissue (red), unmineralized osteoid (bright red), and cell nuclei (black). For histomorphometry analysis, we used Fiji (ImageJ version 1.54p ^[69]^) and a custom-made macro written in the ImageJ Macro Language (available on Zenodo: 10.5281/zenodo.23324947) and based in previous publications^[70,71]^, to automate the segmentation and area measurement of new bone, monetite and soft tissue. Prior to segmentation, images were calibrated using a scale bar reference provided by the software interface. The analysis followed a hierarchical segmentation protocol: first, a Total region of interest (ROI_Tt) was manually defined for each specimen to encompass the entire bone defect site, typically standardized to a length of 7 mm and a width of 3 mm. To assess spatial healing patterns, the software automatically subdivided the ROI_Tt into Proximal (ROI_Pr) and Distal (ROI_Dt) sub-regions for independent quantification. Tissue components were identified through a combination of color thresholding and manual refinement. The following tissues were segmented: monetite, new woven area, cartilage, cortical bone and void area (Figure S8). Each subsequent tissue mask was subtracted by the previous masks in the order of segmentation. Finally, soft tissue was defined as the remaining area within the ROI_Tt after subtracting the previous components listed above. Area measurements (mm^2^) were converted into relative volume fractions. In accordance with standard histomorphometric nomenclature, the % of new bone (TtB.Ar), monetite (TtSc.Ar) and soft tissue (TtSoT.Ar) were calculated relative to the available regeneration space, after subtracting the cortical and void area from the total area.

TRAP staining was employed to identify osteoclastic activity, which exhibited a bright red signal in tartrate-resistant acid phosphatase-positive cells and were quantified via manual cell counting.

Last, Alcian blue + Picrosirius Red staining was used to visualize the distribution and type of collagen and proteoglycans with both bright field and polarized light microscopy. Under bright field, cell nuclei (black), total collagen (red), and proteoglycan-rich cartilage matrix (blue) were observed. Under polarized light we take advantage of the enhanced birefringence as the elongated Picrosirius Red dye molecules attach to collagen fibres parallel to their long axis ^[72]^.

Quantitative polarized light microscopy was performed with a polarizer and analyser placed with 45° difference with respect to each other, to evaluate collagen fibre organization. Under polarized light, thick, mature collagen fibres appeared orange-reddish, while thinner, immature fibres appeared green. Polarized light images of three representative samples per group were acquired using six distinct polarizer-analyser angle (0-45°, 15-60°, 30-75°, 45-90°, 60-105° and 75-120°) for quantification, following the procedure described previously ^[11]^. Briefly, the six images per sample were superimposed using the Fiji Image J 1.54p plugin StackReg ^[73]^. The collagen intensity was measured in four regions of interest (ROI): “cortical” (original bone as a control), “bridged” (new bone bridging the defect), “interphase” (new bone adjacent to monetites) and “intermonetite” (new bone located between monetites) (Figure S10). Measurements were performed in squares of 200 x 200 μm^2^ area, in the different ROIs and through the six-image stacks. The predominant collagen angle orientation was identified as the polarizer-analyser angle with the maximum intensity per region. A total of n>6 random measurements per ROI were performed in each sample. Finally, the percentage of measurements with maximum intensity at polarization light 45° (analyser 90°) was calculated per experimental group and region. For final visualization all resulting six images from the polarizer-analyser angles were merged.

### Backscattered electron microscopy (BSE)

Bone infiltration and in vivo monetite degradation was further analysed with BSE mode using a field-emission scanning electron microscope (FE-SEM; JEOL JSM-IT800, JEOL Ltd., Japan). For this, the remaining PMMA-embedded blocks after histological cutting of three representative samples per chosen groups were imaged under high-vacuum conditions. BSE images were acquired at an accelerating voltage of 7 kV, a working distance of 4.4-6.5 mm, an emission current of 37.4 µA and a spot size of 4.0. Beam alignment and astigmatism were corrected prior to imaging. Once the mosaic of each sample was assembled, the quantification of bone-monetite integration was quantified using ORS Dragonfly segmentation tools.

### Nanoindentation measurements

PMMA-embedded samples were ground on both sides using a PM5 grinding system (Logitech, Glasgow, UK) to obtain parallel surfaces suitable for nanoindentation measurements. The surface designated for indentation was subsequently polished using diamond suspensions with particle sizes of 3, 1, and 0.25 μm to achieve a scratch-free finish.

Nanoindentation measurements were performed using a UB1 Hysitron nanoindenter (Hysitron Inc., Minneapolis, MN, USA) equipped with a tetrahedral diamond Berkovich indenter tip. The indenter tip was characterized by an elastic modulus of 1140 GPa and a Poisson’s ratio of 0.07. Indentation tests were conducted in load-function mode.

The indentation protocol was carried out with a peak load of 5000 μN, applied at a loading rate of 1000 μN/s. At maximum load, the force was maintained for 60 s before partial unloading to 1000 μN at 400 μN/s. A second hold segment of 10 s was then introduced, followed by complete unloading to 0 μN at a rate of 200 μN/s. An extended holding period at peak load was implemented to reduce the influence of viscoelastic deformation and creep on the measurements. Additionally, the intermediate hold during unloading, performed at approximately 90% of the maximum load, allowed correction for thermal drift in the displacement signal, including contributions from thermal expansion of both the indenter tip and the instrument ^[74]^.

The local elastic indentation modulus (*E_r_*) was determined according to the Oliver and Pharr method ^[75]^. Hardness (*H*) was calculated as the ratio between the maximum applied load (*P*) and the projected contact area (*A*), according to *H = P/A*. Three representative samples per chosen groups were used to map these parameters. In addition to the four ROIs defined in the polarized light microscopy analysis, another region representing the bone infiltrations into the monetite was analysed (named “infiltrations”) (Figure S10). A minimum of 5 measurements were obtained per predefined ROI and sample. In each sample, a grid of indents per ROI with a separation of 30 μm to 50 μm in both x and y direction was used. Before each grid measurement, the “quick approach” function was used to identify the surface and ensure safe contact.

### Scanning small-angle X-ray scattering (SAXS)

For small-angle X-ray scattering (SAXS) measurements, sections with an initial thickness of 200 μm to 240 μm were prepared using a low-speed diamond saw (Isomet, Buehler GmbH, Düsseldorf, Germany). The sections were subsequently ground using 1000 and 2000 grit silicon carbide papers to a final thickness of approximately 100 μm.

SAXS measurements were performed using a Bruker Nanostar system (Karlsruhe, Germany) equipped with a 2D VÅNTEC-2000 detector and a microfocus X-ray source operated with Cu Kα radiation (λ = 1.5418 Å). The beam size at the sample position was approximately 115 µm. The detector was a Xe-based gaseous avalanche detector with an active area and window size of (14 × 14) cm², comprising (2048 × 2048) pixels^2^ with a pixel size of (68 × 68) µm². Measurements were carried out at an accelerating voltage of 50 kV and a current of 600 µA. Detector calibration was performed using silver behenate as the calibration standard. All measurements were conducted under vacuum at approximately 0.02 mbar. Samples were first scanned briefly to obtain a transmission map for localization of different ROIs. Scattering patterns were then recorded with 60 min exposure per selected spot.

Data processing and analysis were performed using DPDAK ^[76]^ and employing custom T- and ρ-parameter plugins. Three representative samples per chosen groups were used for SAXS analysis. Same ROIs defined for nanoindentation measurements were used for this analysis (Figure S10). In each sample, several measurements per ROI were obtained (n > 10 in “bridged”; n > 4 in “cortical”, “interphase”, “intermonetite” and “infiltrations” when present).

### Statistical analysis

All statistical analyses were conducted using the statistical software GraphPad Prism 10.6.1. Prior to hypothesis testing, data were assessed for normality using the Shapiro–Wilk test and for homogeneity of variances using the F test. When variances were unequal, Welch’s correction was applied. Data are presented as mean ± standard deviation (SD) when criteria for parametric test was fulfilled, while boxplots showing median and interquartile range were used in the non-parametric cases. Statistical differences between all the experimental groups were assessed by analysis of variance (one-way ANOVA or its equivalent non-parametric Kruskal-Wallis test). Bonferroni post hoc analysis for multiple comparison was applied. When needed, comparisons between independent groups were performed using an unpaired t-test, or its equivalent non-parametric Mann-Whitney U test. A p-value of <0.05 was considered for statistical significance.

## Acknowledgements

The authors would like to thank Marco Moller from CIC biomaGUNE for the technical support with BSE imaging. We are grateful to former Cipitria Lab members at the Max Planck Institute, Daniela Garske and Jeanette Steffen, for initial protocol establishment and shipping of samples, as well as Hubert Taïeb for creating the 3D printed moulds to fabricate the smaller monetites. We thank the workshop at the Max Planck Institute for fabricating the wedges for controlled-angle cutting and Birgit Schonert for sample preparation. We thank the current Cipitria Lab manager, Iune Toledo, for helping with the general lab work and shipping of samples. We thank Mario Thiele from the Julius Wolff Institute at the Charité for providing the code for histomorphometry.

## Author contribution - CreDit section

• OMI: Methodology; Software; Validation; Formal analysis; Investigation; Data Curation; Writing - Original Draft; Writing - Review & Editing; Visualization

• EML: Methodology; Validation; Investigation; Data Curation; Writing - Review & Editing

• LI: Validation; Formal analysis; Investigation; Data Curation; Writing - Review & Editing; Visualization

• IAZ: Validation; Formal analysis; Investigation; Data Curation; Writing - Original Draft; Writing - Review & Editing; Visualization

• DW: Methodology; Data Curation

• JAW: Methodology; Data Curation; Writing - Review & Editing; Visualization

• VJ: Methodology; Validation; Investigation; Resources; Data Curation; Writing - Original Draft; Writing - Review & Editing; Visualization

• PMT: Methodology; Validation; Investigation; Resources; Data Curation; Writing - Review & Editing; Visualization

• TLM: Methodology; Validation; Data Curation

• RLB: Methodology; Validation; Data Curation

• OS: Methodology; Data Curation; Writing - Review & Editing

• LZ: Methodology; Data Curation

• WW: Methodology; Data Curation; Writing - Review & Editing

• SA: Methodology; Validation; Investigation; Data Curation; Writing - Original Draft; Visualization

• JPS: Methodology; Data Curation; Writing - Review & Editing; Visualization

• TW: Methodology; Writing - Review & Editing; Visualization

• BH: Methodology; Data Curation; Writing - Review & Editing; Visualization

• PGG: Project administration; Supervision; Writing - Review & Editing; Visualization

• SR: Project administration; Supervision

• EG: Project administration; Supervision

• DMR: Methodology; Validation; Investigation; Resources; Data Curation; Writing - Original Draft; Writing - Review & Editing; Visualization

• MJD: Conceptualization, Methodology, Validation, Resources, Writing - Review & Editing, Supervision, Project administration, Funding acquisition

• FGM: Conceptualization, Methodology, Validation, Resources, Writing - Original Draft, Writing - Review & Editing, Supervision, Project administration, Funding acquisition

• MSS: Conceptualization, Methodology, Validation, Resources, Writing - Original Draft, Writing - Review & Editing, Supervision, Project administration, Funding acquisition

• AC: Conceptualization, Methodology, Validation, Resources, Writing - Original Draft, Writing - Review & Editing, Supervision, Project administration, Funding acquisition

## Data Availability Statement

All raw and processed data, as well as custom-made deep learning models and Fiji macro for histomorphometry are available on Zenodo (10.5281/zenodo.23324947).

