## Supplementary Information for "Material-driven fibronectin assembly enables low-dose BMP-2 to regenerate structurally mature bone in critical-sized load-bearing defects"

Figure S1: Characterisation of PEA-coated monetites and in vitro biological evaluation.

Figure S2: Effect of monetite quantity in bone regeneration.

Figure S3: Characterisation of PRP functionalization and in vitro biological evaluation.

Figure S4: BSE images of selected groups with 10 µg/ml of rhBMP-2.

Figure S5: BSE images of selected groups with 25 µg/ml of rhBMP-2.

Figure S6: BSE images depicting the seamless interface between the monetite and new bone, with bone infiltrations showing osteocyte lacuna.

Figure S7: Goldner's trichrome staining images.

Figure S8: Specific tissue components obtained with Movat pentachron-von Kossa staining using a custom-designed automated macro for Fiji.

Figure S9: TRAP staining images with zoomed regions showing the osteoclasts.

Figure S10: Regions of interest defined for collagen fibre alignment, mineral particle thickness and alignment analysis.

Figure S11: Quantitative analysis of collagen fibre orientation presented per sample and ROI in group 2M.

Figure S12: Quantitative analysis of collagen fibre orientation presented per sample and ROI in group 2M_PFB10.

Figure S13: Quantitative analysis of collagen fibre orientation presented per sample and ROI in group 2M_PB25.

Figure S14: Collagen fibre orientation analysis performed with polarized light microscopy for sample 2M_PFB25_1

Figure S15: Collagen fibre orientation analysis performed with polarized light microscopy for sample 2M_PFB25_2

Figure S16: Collagen fibre orientation analysis performed with polarized light microscopy for sample 2M_PFB25_4

Figure S17: Image and quantification of the average mineral particle thickness (T) and the degree of alignment (ρ) obtained with SAXS measurements in the 2M_PFB10 experimental group.

Figure S18: Image and quantification of the average mineral particle thickness (T) and the degree of alignment (ρ) obtained with SAXS measurements in the 2M_PFB25 experimental group.

Figure S19: Image and quantification of the average mineral particle thickness (T) and the degree of alignment (ρ) obtained with SAXS measurements in the 2M_PB25 experimental group.

Figure S20. Quantification of local elastic indentation modulus (Er).

Figure S21. Quantification of hardness (H).

Figure S22. Representative cross-sections and 3D reconstructions of synchrotron microCT images with segmented data visualizing all monetites within sample PF_1.

Figure S23. Representative cross-sections and 3D reconstructions of synchrotron microCT images with segmented data visualizing all monetites within sample PF_2.

Figure S24. Representative cross-sections and 3D reconstructions of synchrotron microCT images with segmented data visualizing all monetites within sample PF_3.

Figure S25. Representative cross-sections and 3D reconstructions of synchrotron microCT images with segmented data visualizing all monetites within sample PF_4.

Figure S26. Representative cross-sections and 3D reconstructions of synchrotron microCT images with segmented data visualizing all monetites within sample PF_5.

Figure S27. Representative cross-sections and 3D reconstructions of synchrotron microCT images with segmented data visualizing all monetites within sample PF_6.

Figure S28. Representative cross-sections and 3D reconstructions of synchrotron microCT images with segmented data visualizing all monetites within sample PF_7.

Figure S29. Representative cross-sections and 3D reconstructions of synchrotron microCT images with segmented data visualizing all monetites within sample PFB25_1.

Figure S30. Representative cross-sections and 3D reconstructions of synchrotron microCT images with segmented data visualizing all monetites within sample PFB25_2.

Figure S31. Representative cross-sections and 3D reconstructions of synchrotron microCT images with segmented data visualizing all monetites within sample PFB25_3.

Figure S32. Representative cross-sections and 3D reconstructions of synchrotron microCT images with segmented data visualizing all monetites within sample PFB25_4.

Table S1. Primers used for qRT-PCR

Video S1. 3D reconstruction of synchrotron microCT images of an ovine sample.

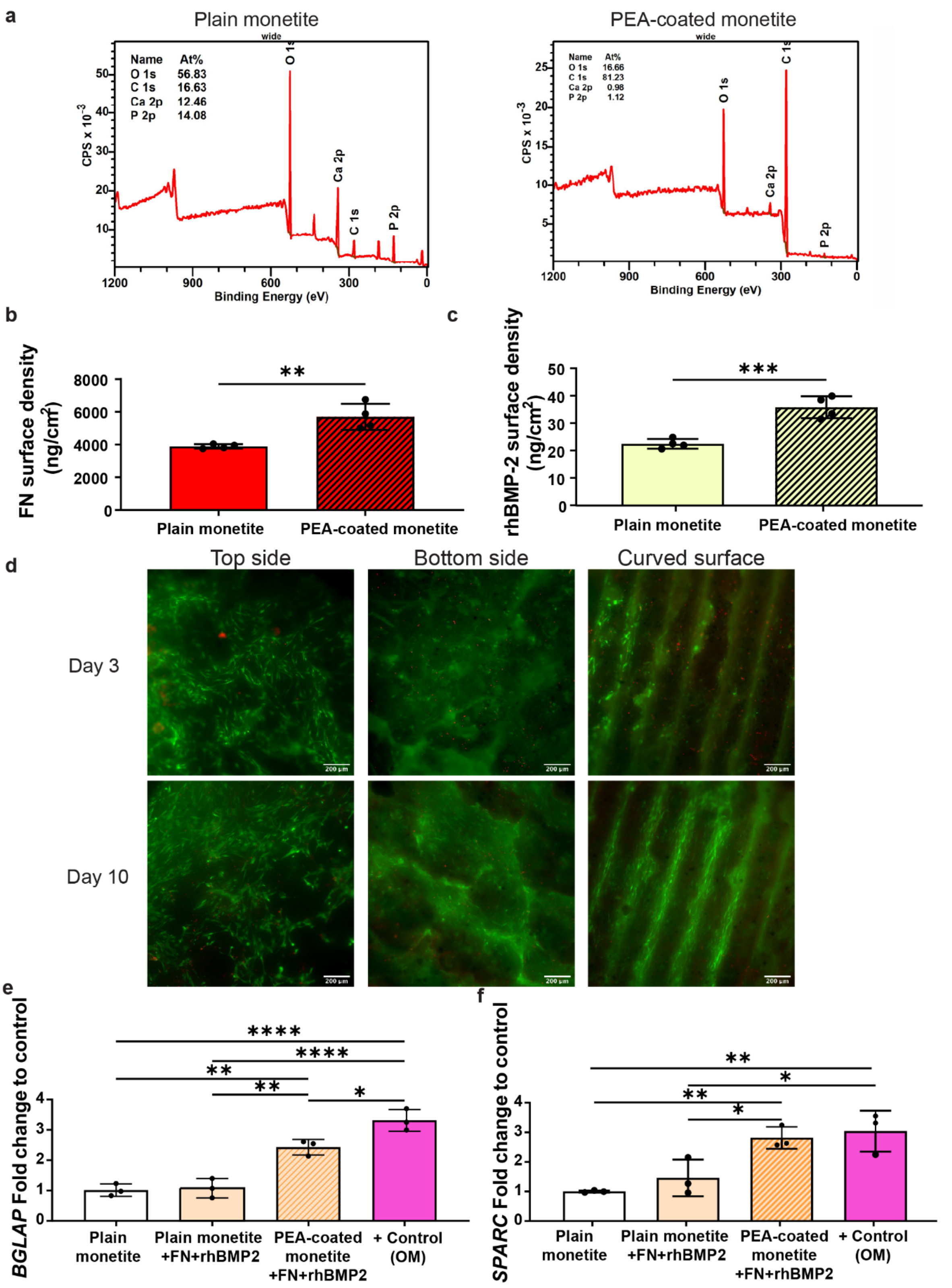


**Figure S1: Characterisation of PEA-coated monetites and in vitro biological evaluation.** a) X-ray photoelectron spectroscopy (XPS) spectra of uncoated and PEA-coated monetite scaffolds showing the surface elemental composition following plasma polymerisation. b) Quantification of FN adsorption by Bicinchoninic Acid Assay (BCA assay) on plain and PEA-coated monetites. c) Quantification of rhBMP-2 adsorption by ELISA on plain and PEA-coated monetites. d) Representative LIVE/DEAD fluorescence images of hASCs cultured on PEA-coated monetites functionalised with FN and rhBMP-2 after 3 and 10 days demonstrating high cell viability on the top, bottom and curved scaffold surfaces. e) Relative expression of osteogenic marker osteocalcin (*BGLAP*) after 28 days of culture. f) Relative expression of osteogenic marker osteonectin (*SPARC*) after 28 days of culture. Data are presented as mean ± standard deviation. Statistical significance is indicated as $p < 0.05$ (*), $p < 0.01$ (**), $p < 0.001$ (***) and $p < 0.0001$ (****).

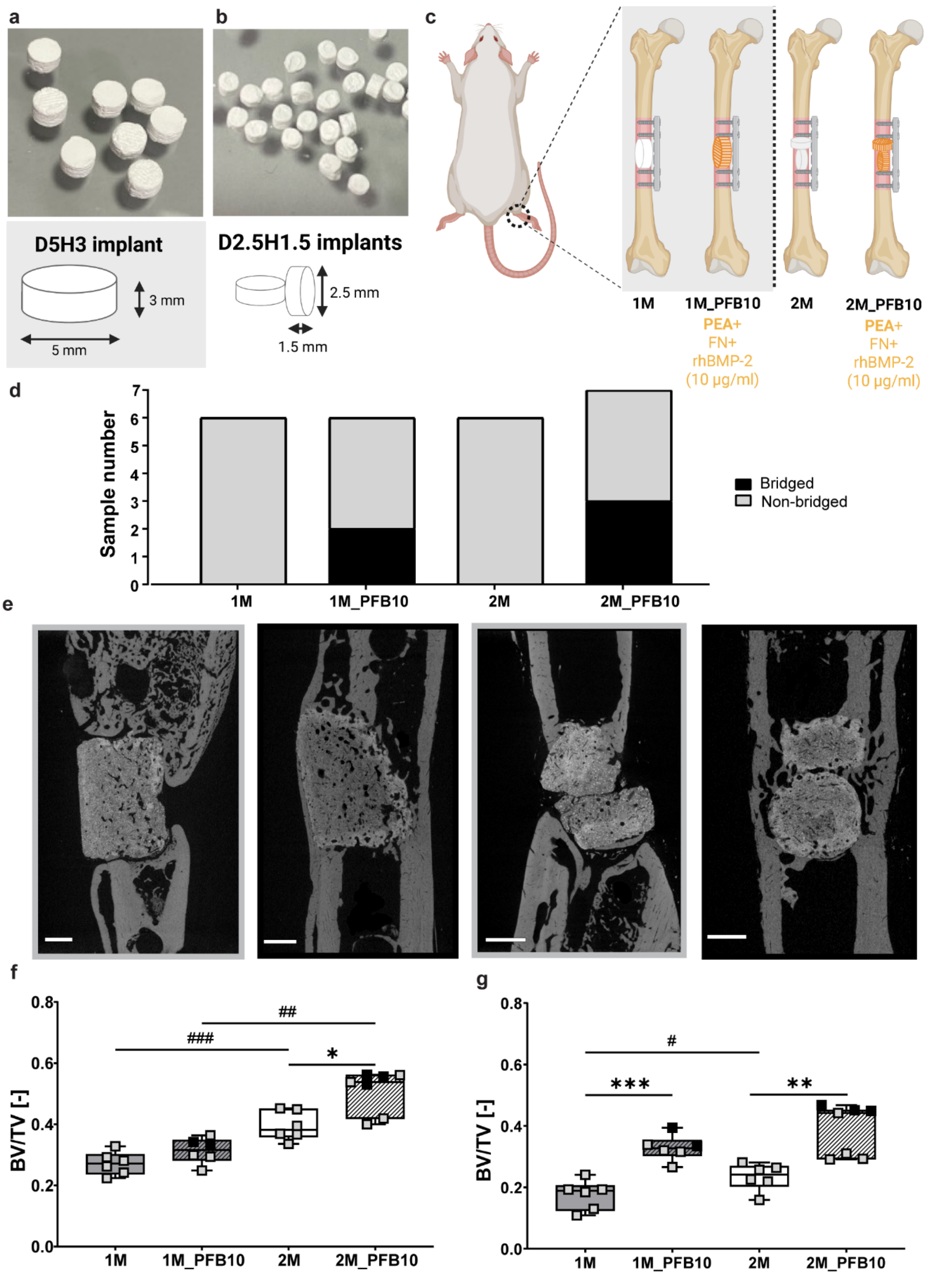


**Figure S2: Effect of monetite quantity in bone regeneration.** Representative images of the two monetite scaffold sizes fabricated for this study: a) large scaffolds (5 × 3 mm) used for in vitro experiments and the material intended for future human applications and b) small scaffolds (2.5 × 1.5 mm) used for in vivo rat implantation. c) Groups with the different coating combinations used in the analysis of the effect of the monetite quantity, represented each by a different colour or pattern: PEA (P, stripes), FN (F, red) and rhBMP-2 (10 µg/ml, B10, yellow). Figure created in BioRender.com. d) Summary of bone bridging analysis for the animals containing either one or two monetites. In this first attempt, either plain monetites or PEA-coated monetites functionalised with FN and 10 µg/ml rhBMP-2 were tested. e) Representative microCT images of each tested group, showing fully bridged (black border) and non-bridged (grey border) examples with only one or two monetites implanted. MicroCT images were analysed with Dragonfly to quantify the BV/TV in the whole defect volume (f) or in the vicinity of the monetite (g). Boxplots show the median and interquartile range, with all the data points (n=5-7) representing the bridged (black) or non-bridged (grey) samples. Statistical analyses have been performed to determine the effect of the monetite quantity (#) or the coating (*) using unpaired Mann-Whitney U test. Statistical significance is indicated as $p < 0.05$ (*/#), $p < 0.01$ (**/##) and $p < 0.001$ (***/###).

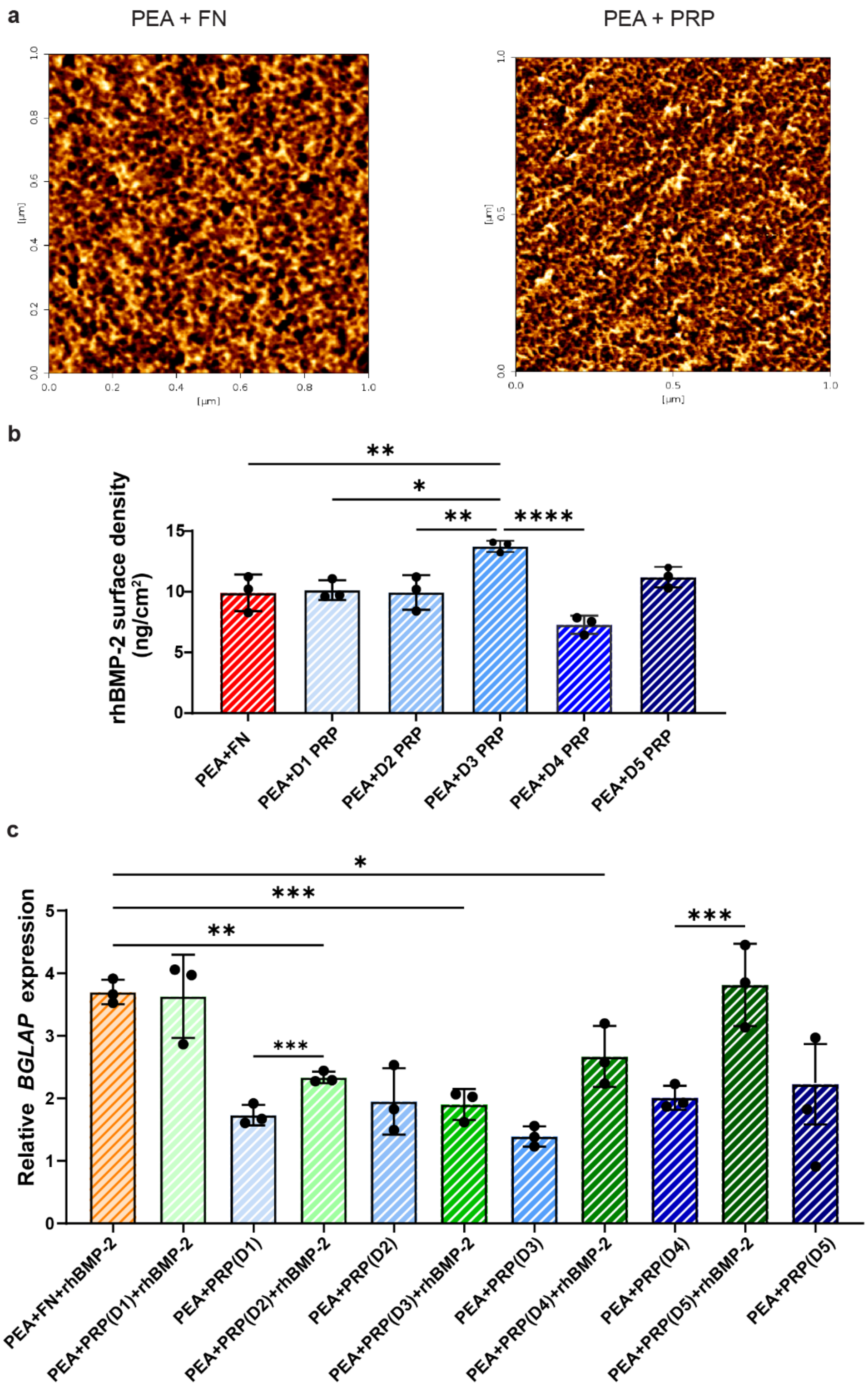


**Figure S3: Characterisation of PRP functionalization and in vitro biological evaluation.** A) Representative AFM height images of FN and PRP coated PEA surfaces illustrating the nanoscale organisation of adsorbed proteins. B) rhBMP-2 adsorption on PEA-coated monetites functionalised with FN or PRP obtained from five different donors. C) Relative osteocalcin (*BGLAP*) gene expression after 28 days for hASCs cultured on PEA-coated monetites functionalised with FN+BMP-2 or PRP-based mixtures.

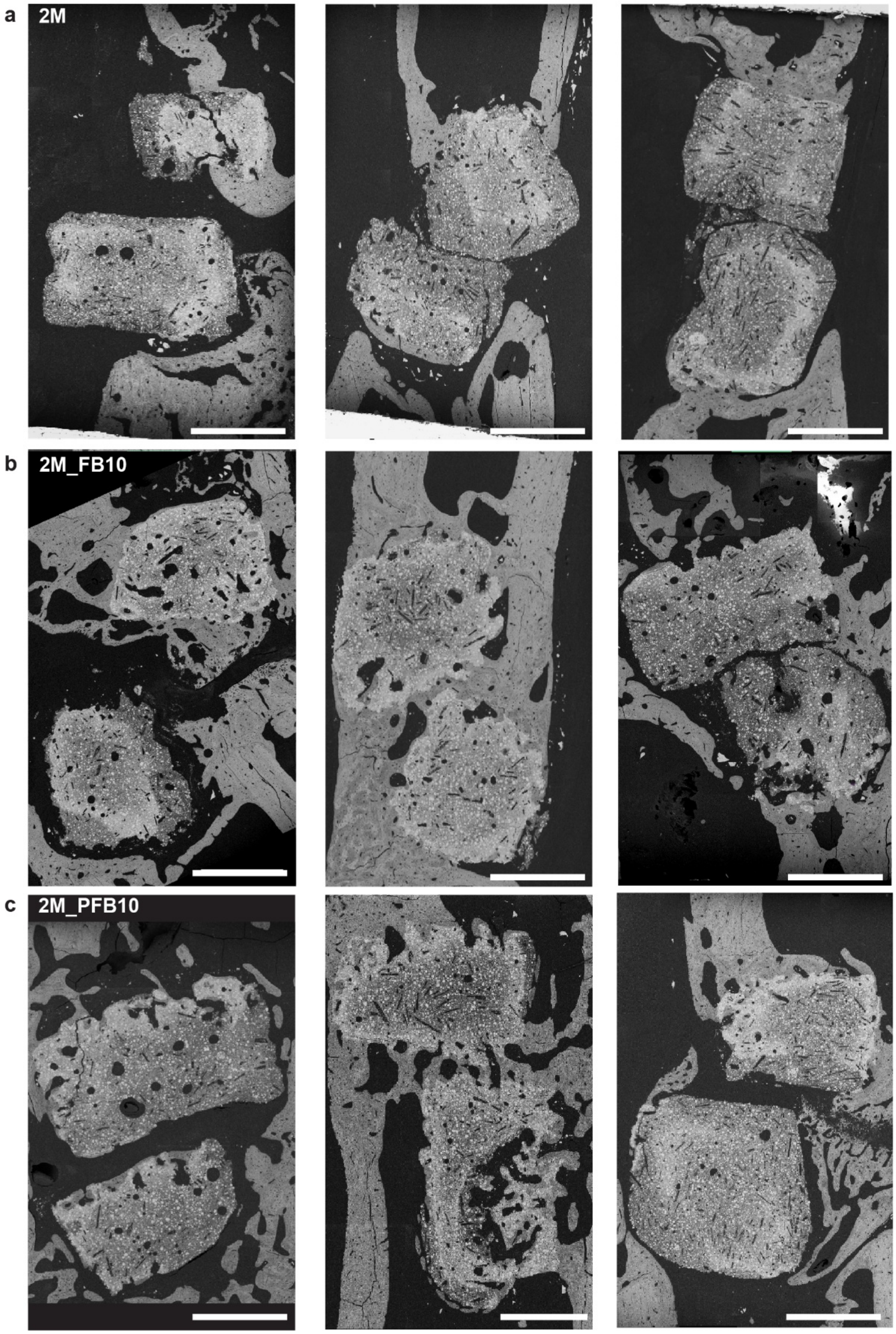


**Figure S4: BSE images of selected groups with 10 μg/ml of rhBMP-2.** BSE images of three samples per selected group, including plain monetite (2M), FN+rhBMP-2 10 μg/ml (2M_FB10) and PEA+FN+rhBMP-2 10 μg/ml (2M_PFB10). Scale bar: 1 mm.

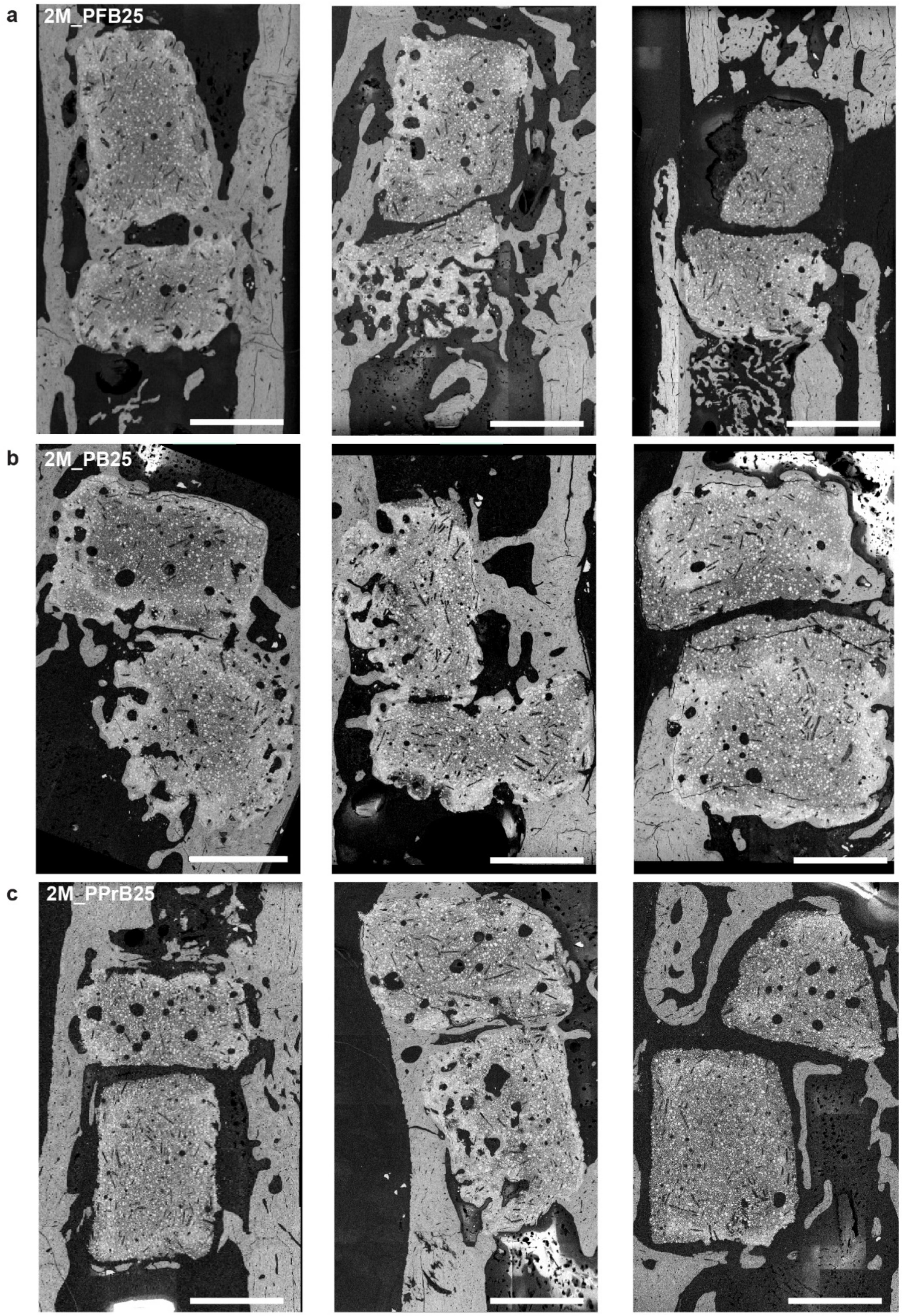


**Figure S5: BSE images of selected groups with 25 µg/ml of rhBMP-2.** BSE images of three samples per selected group, including PEA+FN+rhBMP-2 25 µg/ml (2M_PFB25), PEA+rhBMP-2 25 µg/ml (2M_PB25) and PEA+PRP+rhBMP-2 25 µg/ml (2M_PPrB25) groups. Scale bar: 1 mm.

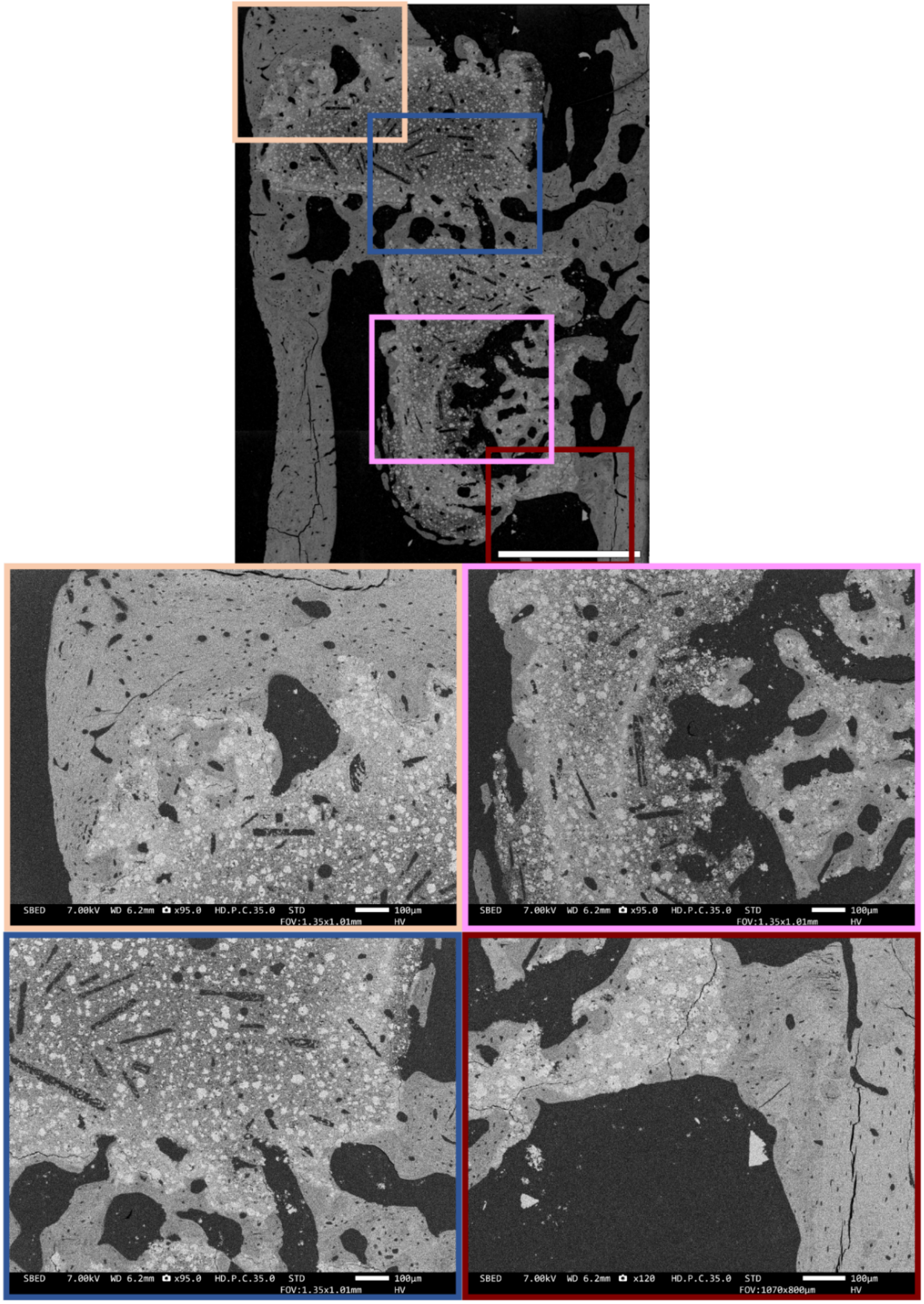


**Figure S6: BSE images depicting the seamless interface between the monetite and new bone, with bone infiltrations showing osteocyte lacuna.** BSE image mosaic with four zoom-in regions, showing the monetite integrated in the bone of a sample of the group PEA+FN+rhBMP-2 10 µg/ml (2M_PFB10). The osteocyte lacunae seen in the cortical bone could be clearly identified inside the new bone infiltrations observed through the monetite, demonstrating new bone in-growth through the ceramic materials. Scale bar of the mosaic: 1 mm, zoom-in regions: 100 µm.

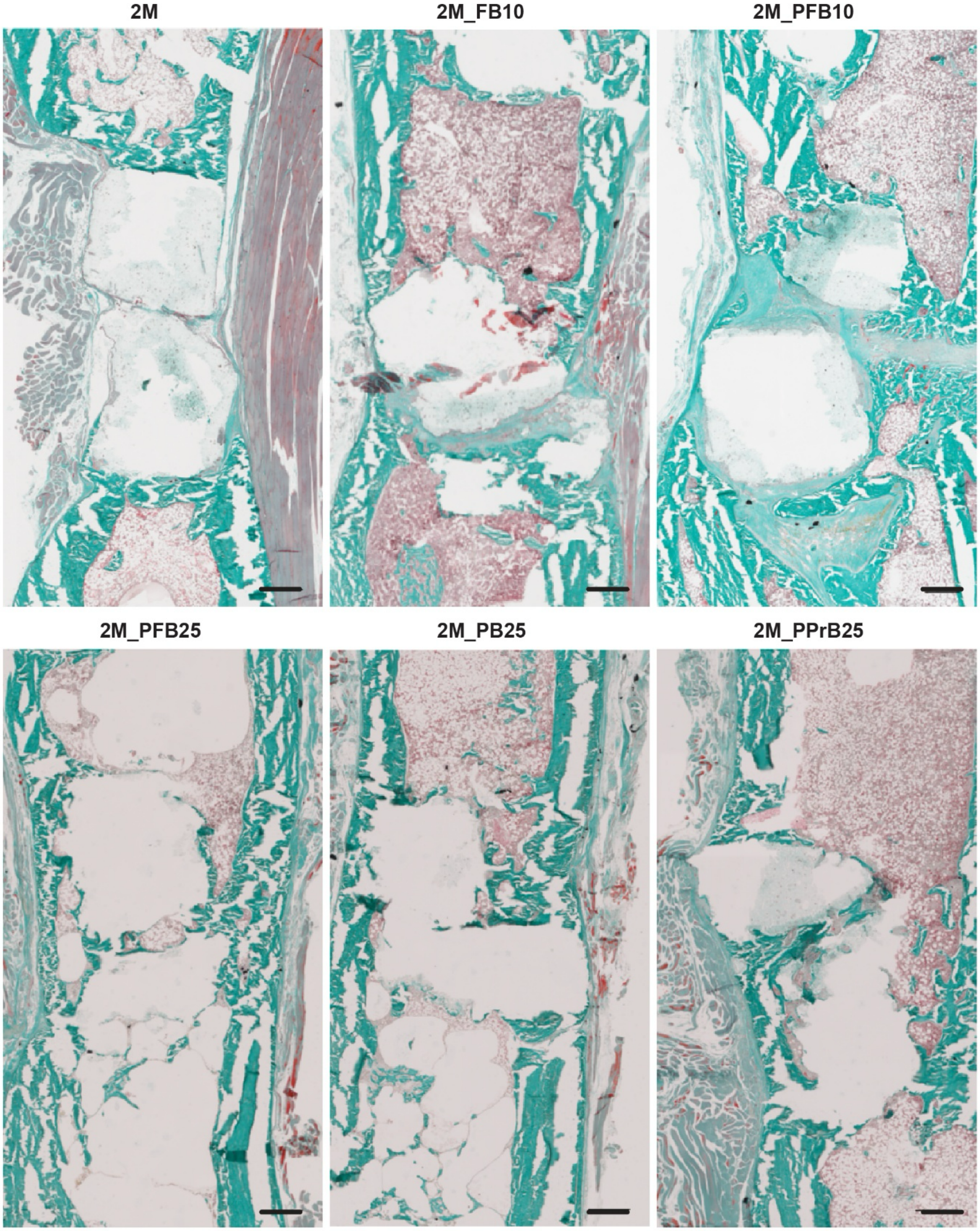


**Figure S7: Goldner's trichrome staining images.** Images of one representative sample of the selected experimental groups, including plain monetite (2M), FN+rhBMP-2 10 µg/ml (2M_FB10), PEA+FN+rhBMP-2 10 µg/ml (2M_PFB10), PEA+FN+rhBMP-2 25 µg/ml (2M_PFB25), PEA+rhBMP-2 25 µg/ml (2M_PB25) and PEA+PRP+rhBMP-2 25 µg/ml (2M_PPrB25) groups. Scale bar: 500 µm.

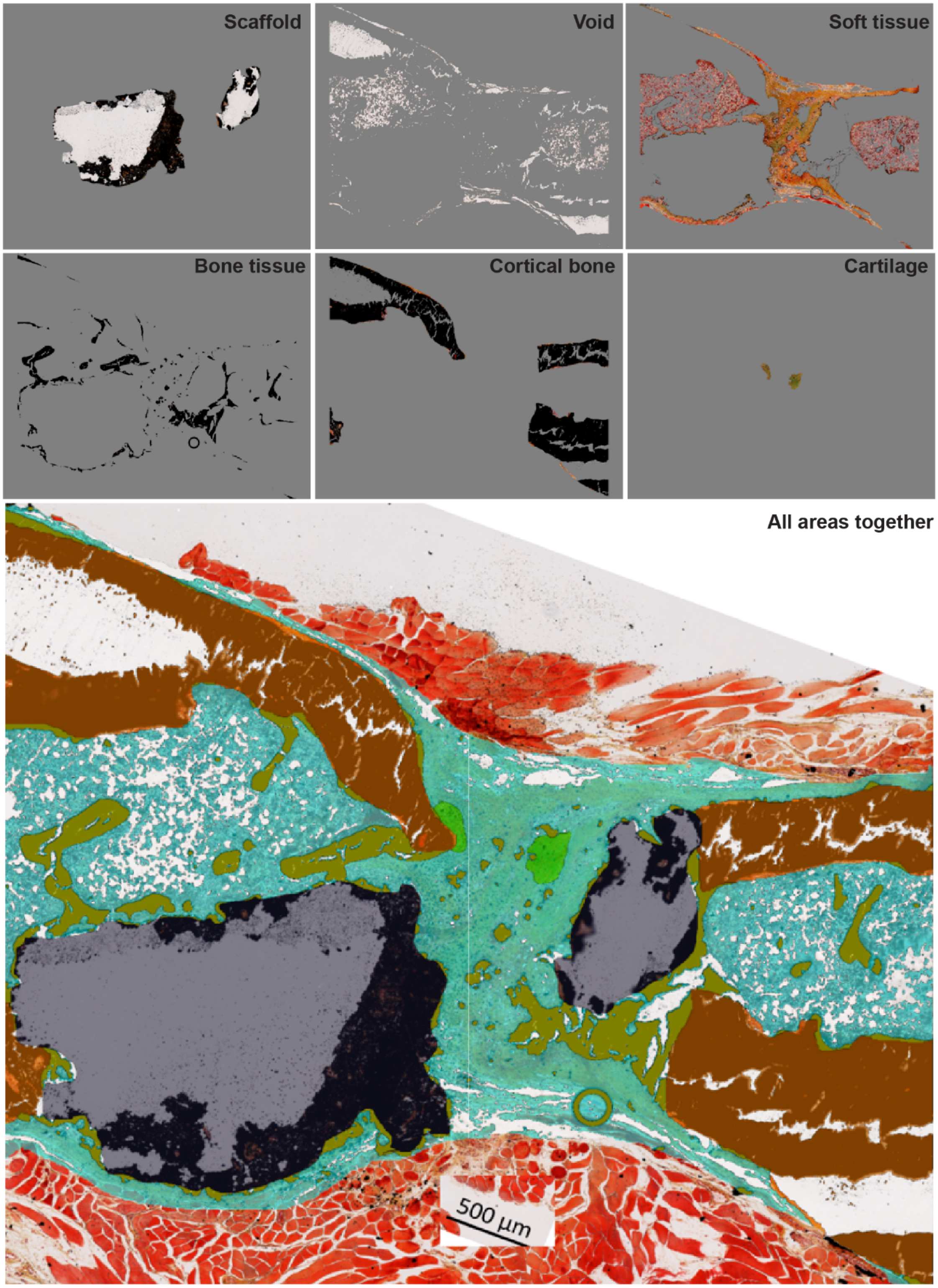


**Figure S8: Specific tissue components obtained with Movat pentachrome-von Kossa staining using a custom-designed automated macro for Fiji.** Specific tissue components are segmented as follows (stating the colour in the merged image): scaffold [greys], void [white], soft tissue [cyan], new trabecular bone tissue [olive green], new cortical bone tissue [brown], cartilage [bright green] and muscle [red]. Last image shows all regions merged.

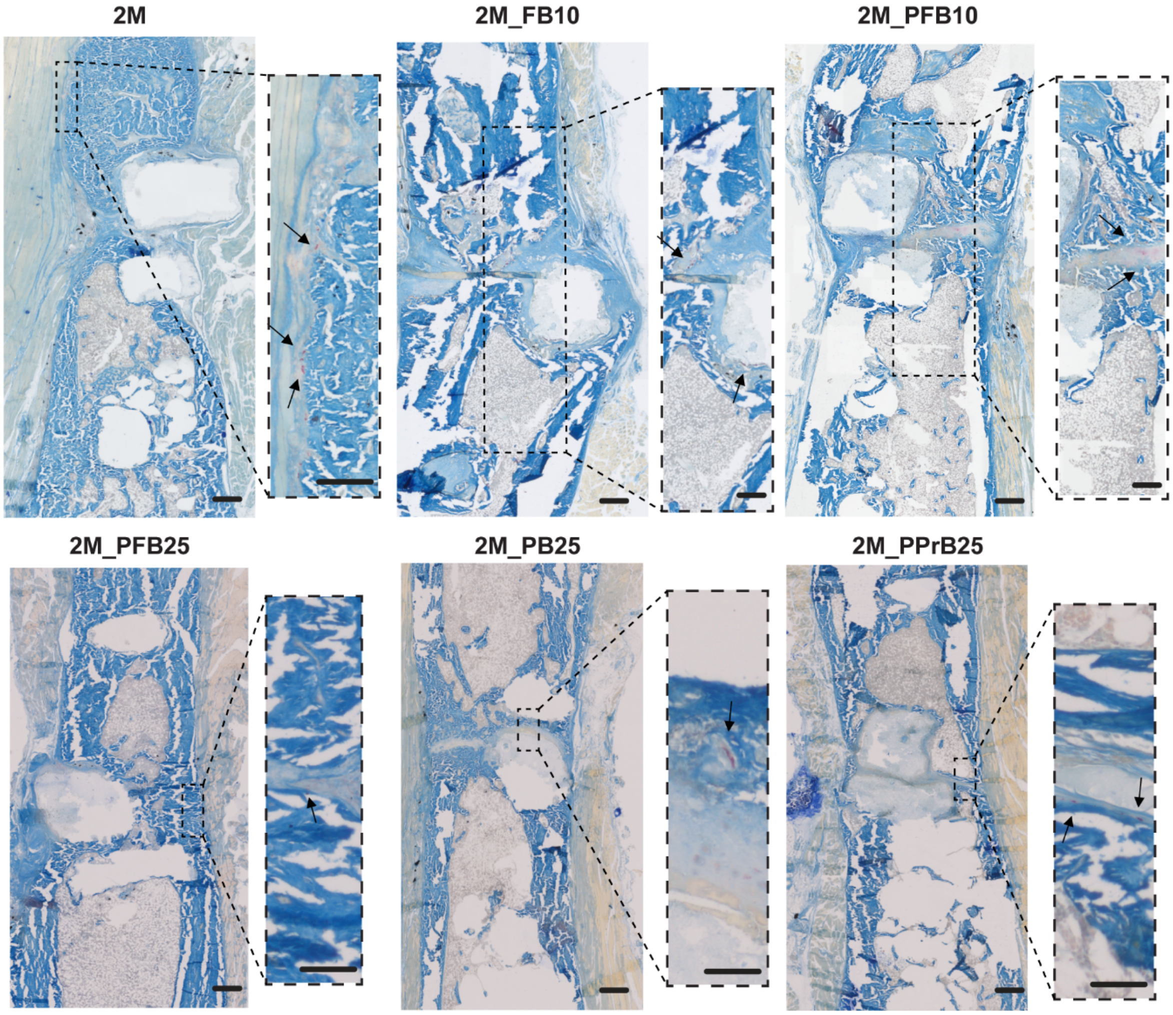


**Figure S9: TRAP staining images with zoomed in regions showing the osteoclasts.** Images of one representative sample of the selected experimental groups, including plain monetite (2M), FN+rhBMP-2 10 µg/ml (2M_FB10), PEA+FN+rhBMP-2 10 µg/ml (2M_PFB10), PEA+FN+rhBMP-2 25 µg/ml (2M_PFB25), PEA+rhBMP-2 25 µg/ml (2M_PB25) and PEA+PRP+rhBMP-2 25 µg/ml (2M_PPrB25) groups. Scale bar of big images: 500 µm. Scale bar of zoomed in images: 200 µm.

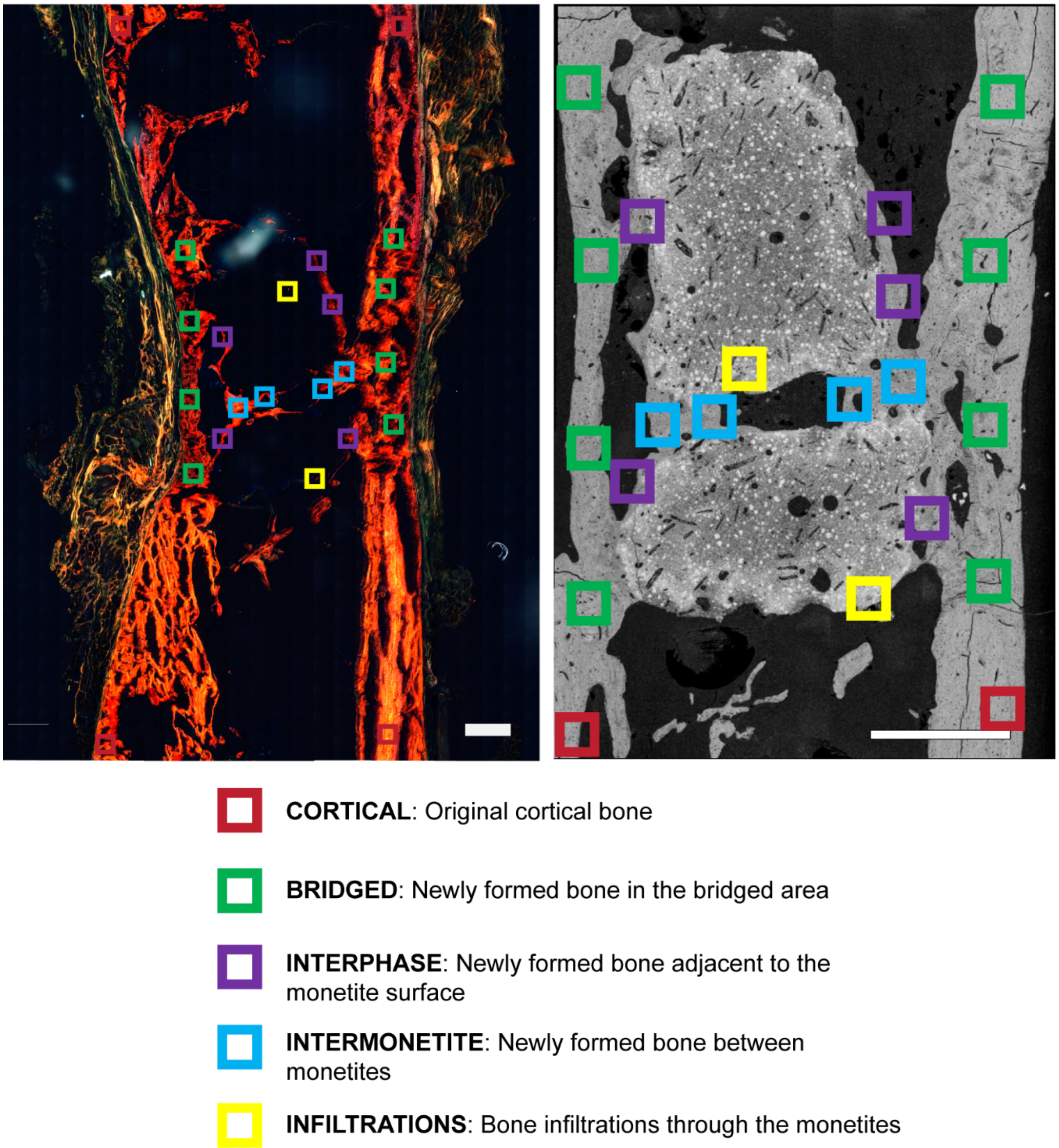


**Figure S10: Regions of interest defined for collagen fibre alignment, mineral particle thickness and alignment analysis**. In the polarized light microscopy analysis, the collagen intensity was measured in four regions of interest (ROI): “cortical” (native cortical bone, red), “bridged” (newly formed bone in the bridging area, green), “interphase” (newly formed bone adjacent to the monetite surface, purple) and “intermonetite” (newly formed bone between monetites, blue). In addition to these four ROIs, another region representing the bone infiltrations into the monetite (named “infiltrations”) was analysed in nanoindentation and SAXS studies. Scale bar of polarized light image: 500 µm. Scale bar of SEM image: 1 mm.

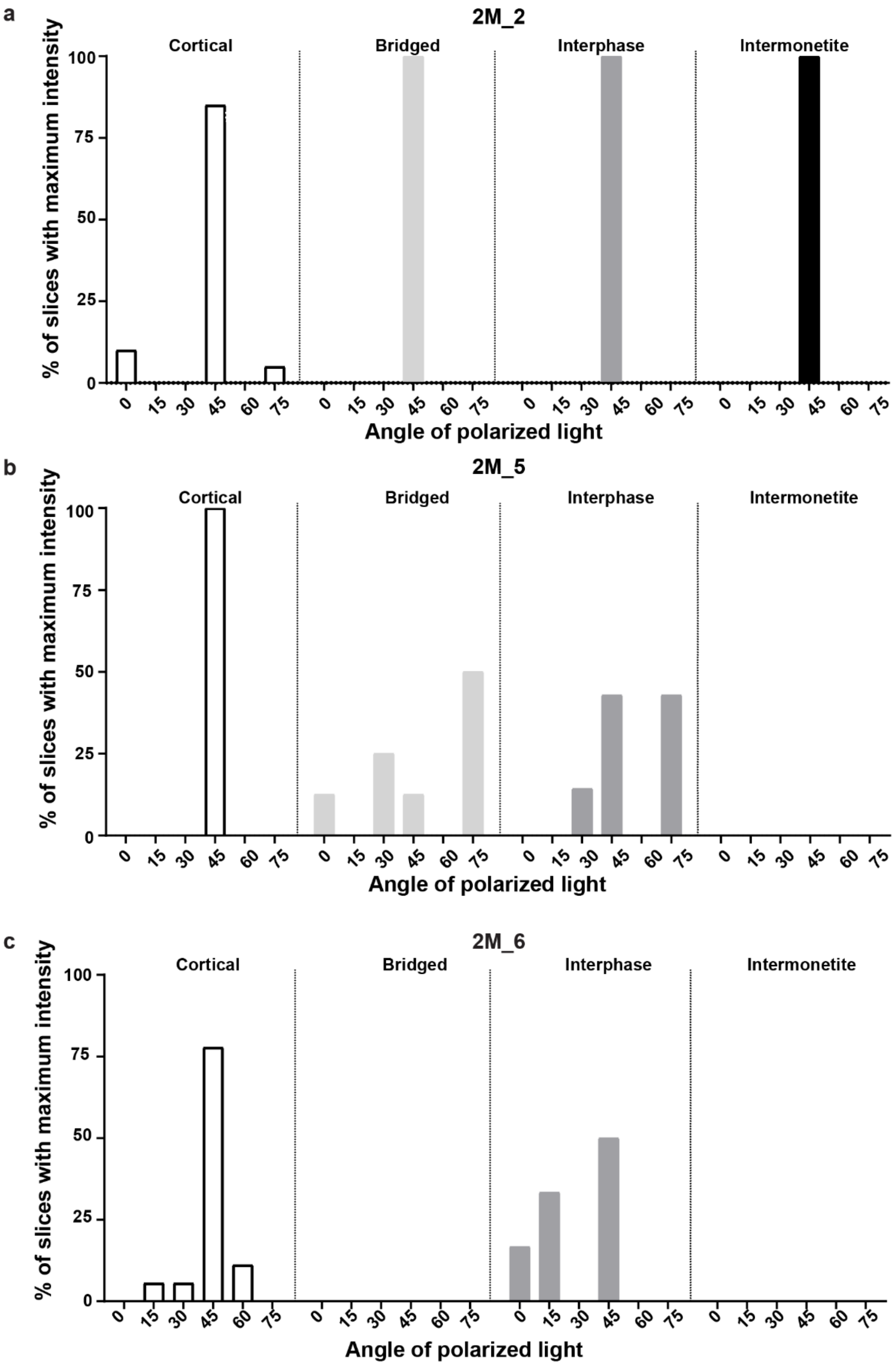


**Figure S11: Quantitative analysis of collagen fibre orientation presented per sample and ROI in group 2M.** Quantification performed by the number of measurements per sample and per ROI (n>6), in 3 samples of 2M group: a) #2, b) #5 and c) #6. Measurements show the percentage of counts that exhibit a maximum intensity in an area of 200 x 200 µm$^2$ for an angle of polarizer/analyzer equal to 0-45º, 15-60°, 30-75°, 45-90°, 60-105° and 75-120°. Samples are characterized in the four ROIs presented in Figure S10: cortical (white), bridged (light grey), interphase (dark grey) and intermonetite (black). The percentage of measurements with maximum intensity per polarizer angle is presented by region and sample.

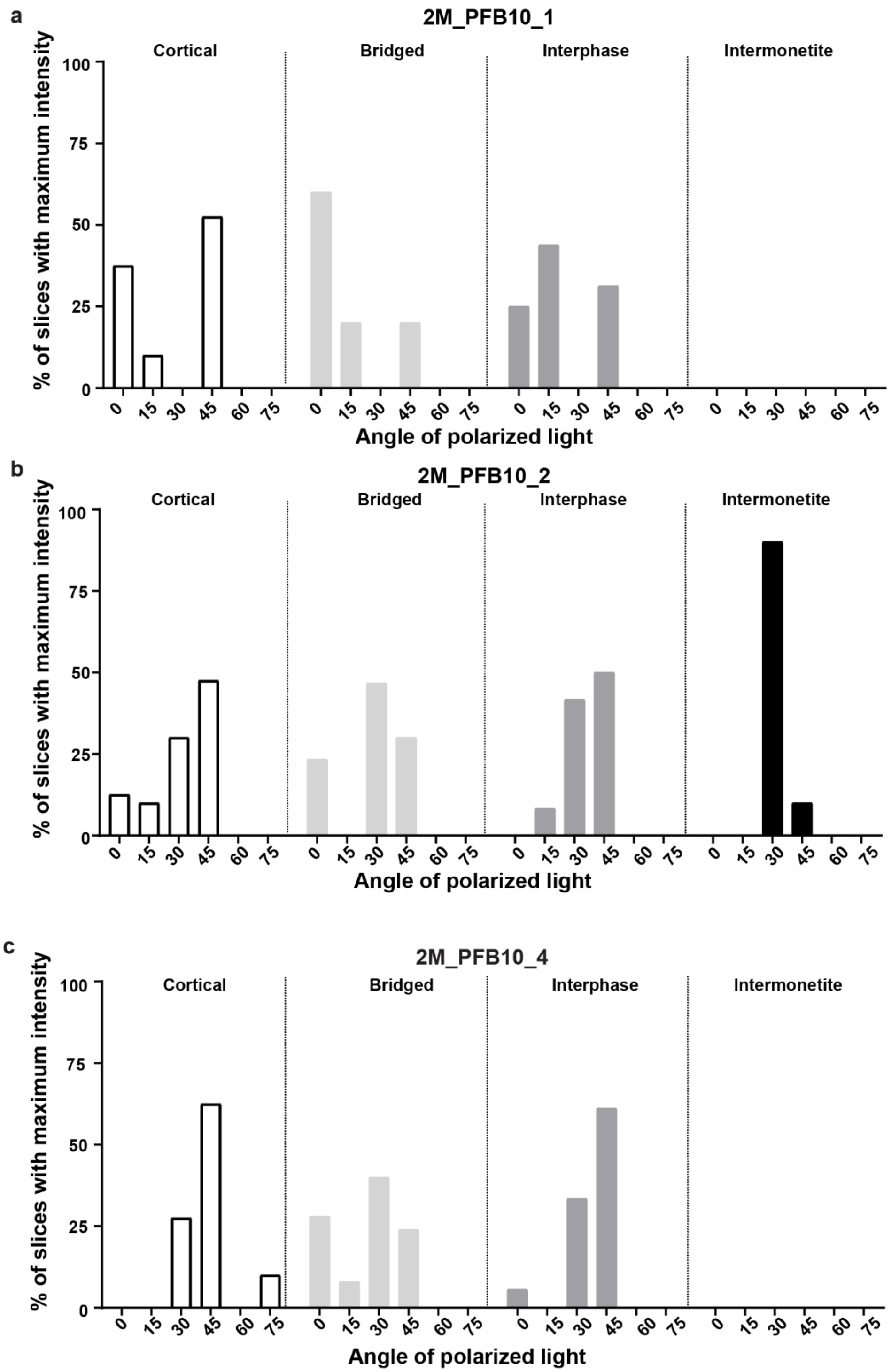


**Figure S12: Quantitative analysis of collagen fibre orientation presented per sample and ROI in group 2M_PFB10.** Quantification performed by the number of measurements per sample and per ROI (n>6), in 3 samples of 2M_PFB10 group: a) #1, b) #2 and c) #4. Measurements show the percentage of counts that exhibit a maximum intensity in an area of 200 x 200 µm$^2$ for an angle of polarizer/analyzer equal to 0-45º, 15-60°, 30-75°, 45-90°, 60-105° and 75-120°. Samples are characterized in the four ROIs presented in Figure S10: cortical (white), bridged (light grey), interphase (dark grey) and intermonetite (black). The percentage of measurements with maximum intensity per polarizer angle is presented by region and sample.

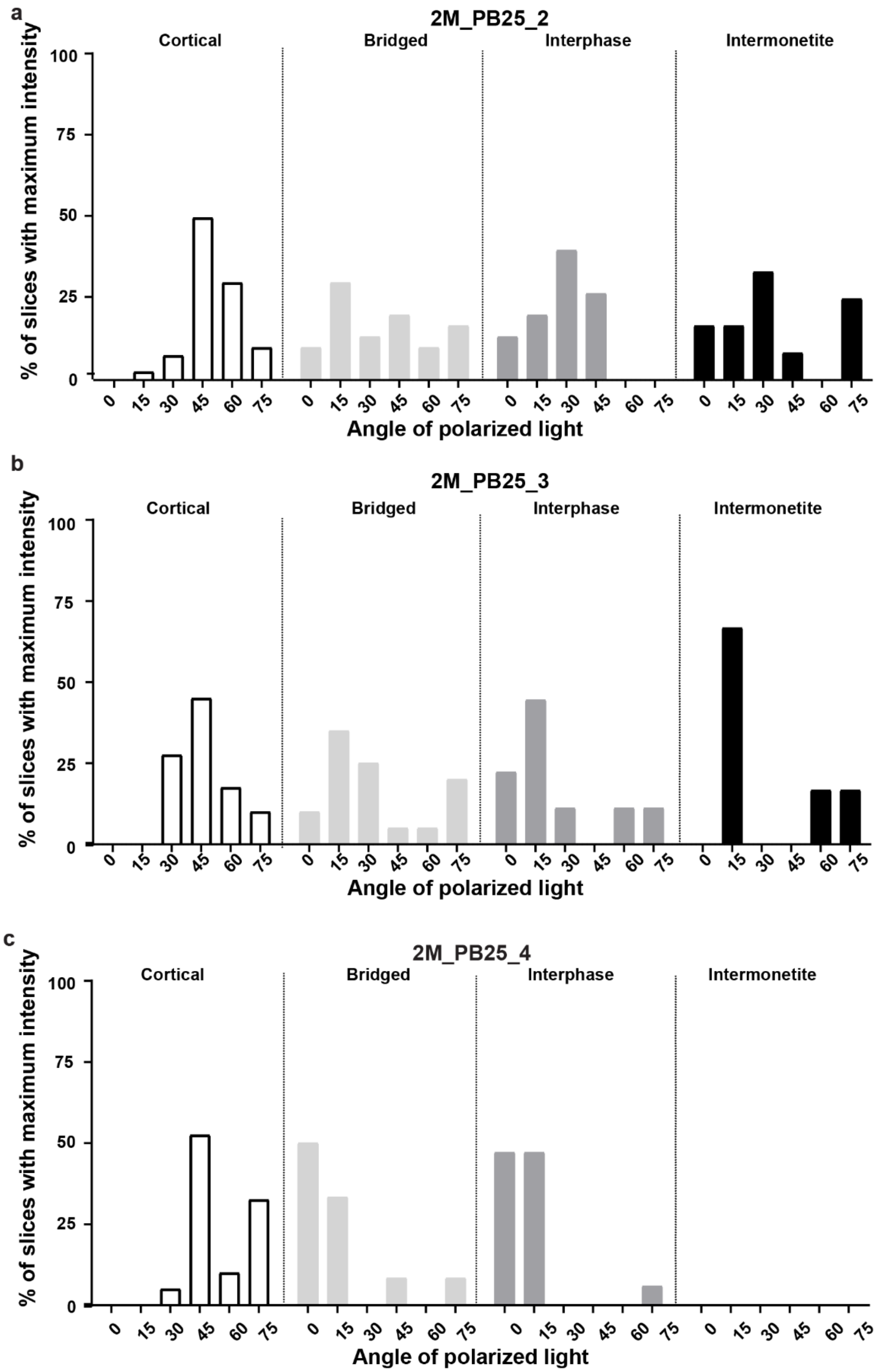


**Figure S13: Quantitative analysis of collagen fibre orientation presented per sample and ROI in group 2M_PB25.** Quantification performed by the number of measurements per sample and per ROI (n>6), in 3 samples of 2M_PB25 group: a) #2, b) #3 and c) #4. Measurements show the percentage of counts that exhibit a maximum intensity in an area of 200 x 200 μm² for an angle of polarizer/analyzer equal to 0-45º, 15-60°, 30-75°, 45-90°, 60-105° and 75-120°. Samples are characterized in the four ROIs presented in Figure S10: cortical (white), bridged (light grey), interphase (dark grey) and intermonetite (black). The percentage of measurements with maximum intensity per polarizer angle is presented by region and sample.

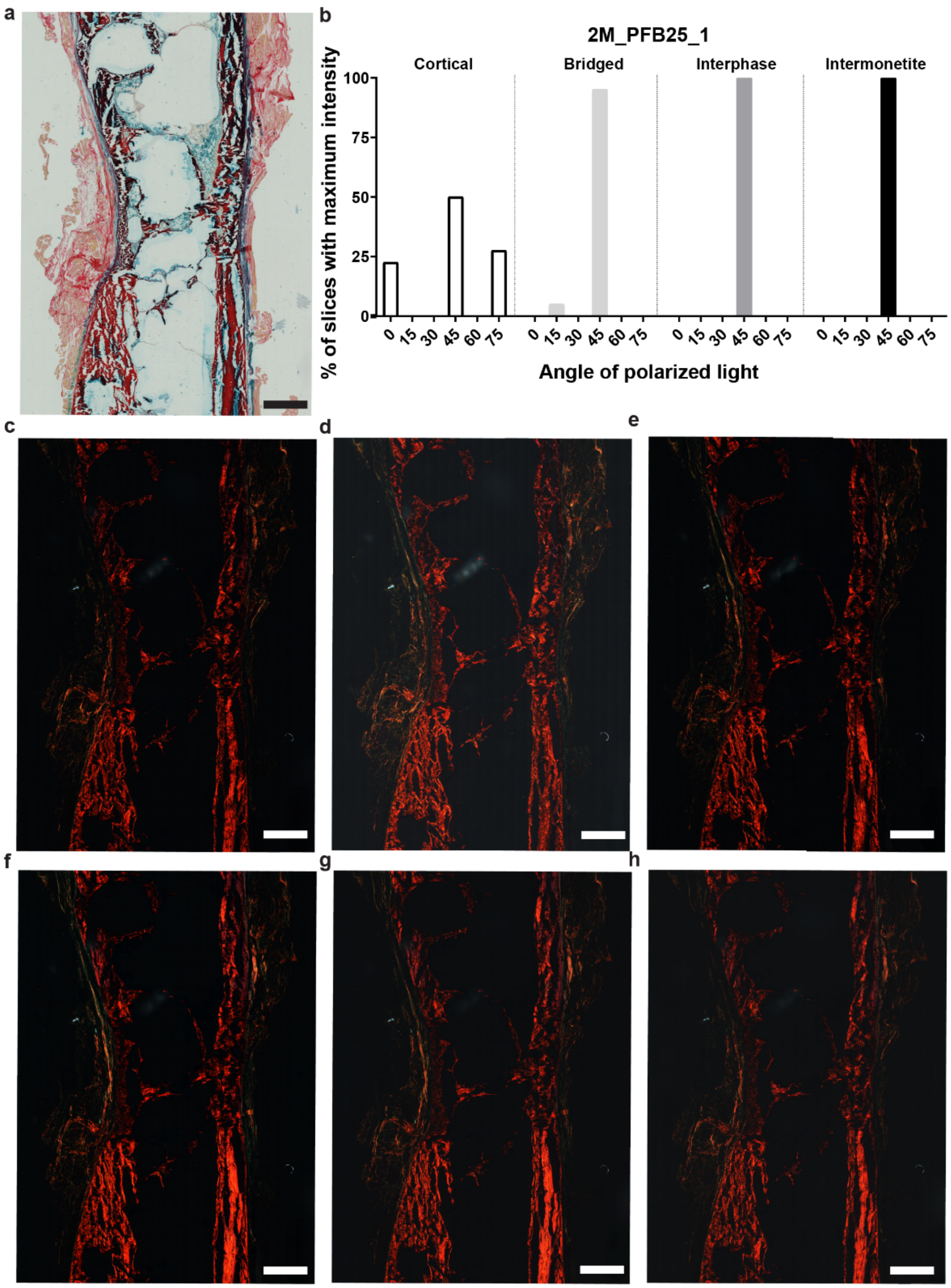


**Figure S14: Collagen fibre orientation analysis performed with polarized light microscopy for sample 2M_PFB25_1.** a) Alcian blue and picrosirius red staining bright field image of sample 2M_PFB25_1. b) Quantitative analysis of collagen fibre orientation as measured by the number of measurements per ROI (n>6), that exhibit a maximum intensity in an area of 200 x 200 μm$^2$ for different polarizer angles. Samples are characterized in the four ROIs presented in Figure S10: cortical (white), bridged (light grey), interphase (dark grey) and intermonetite (black). The percentage of measurements with maximum intensity per polarizer angle is presented by region and sample. Images of alcian blue and picrosirius red staining of 2M_PFB25_1 sample under six angles of polarized light for each of the corresponding polarizer-analyser angles: 0°-45º, 15°-60º, 30°-75º, 45°-90º, 60°-105º and 75°-120º. Scale bars: 1000 μm.

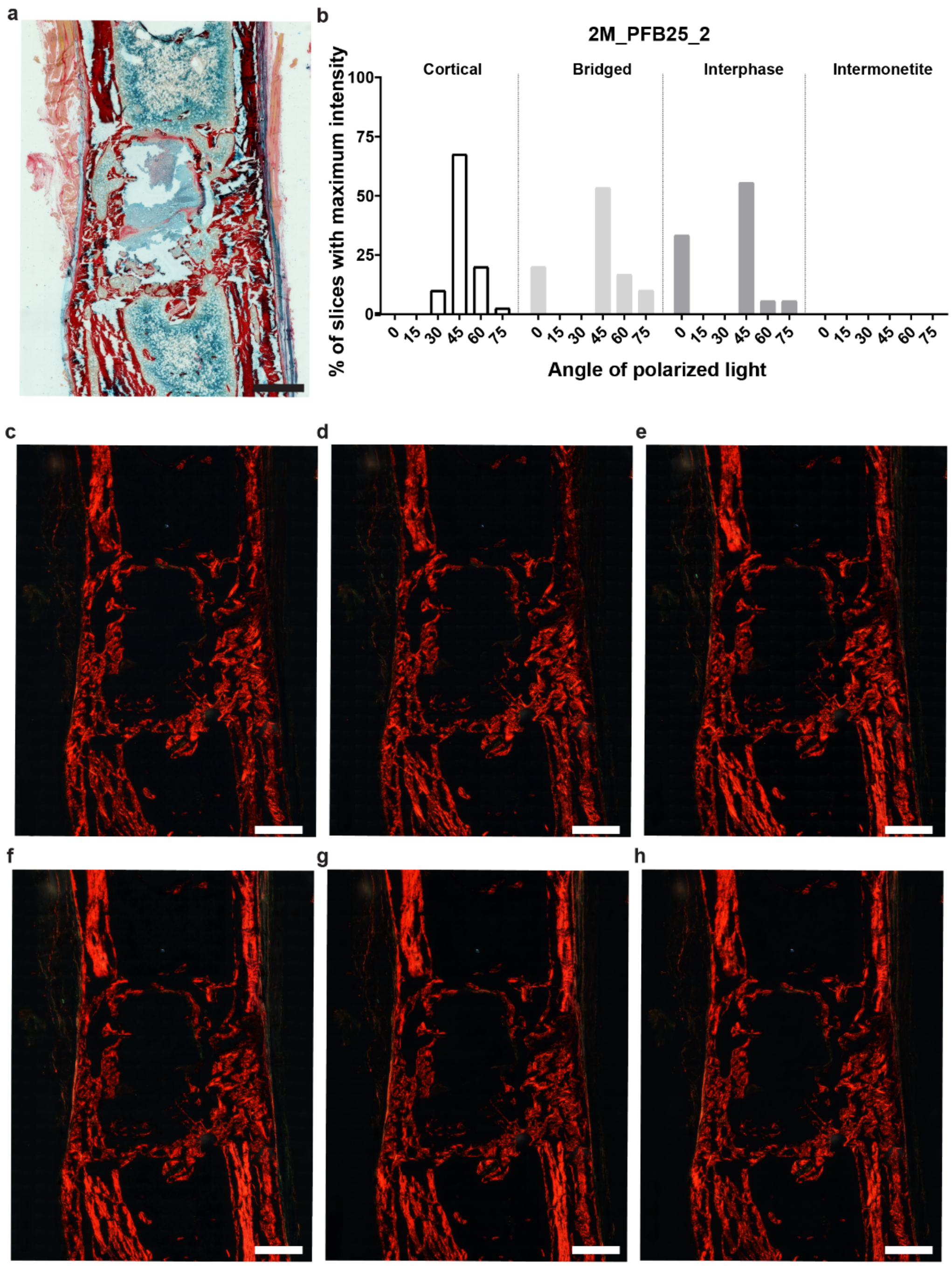


**Figure S15: Collagen fibre orientation analysis performed with polarized light microscopy for sample 2M_PFB25_2.** a) Alcian blue and picrosirius red staining bright field image of sample 2M_PFB25_2. b) Quantitative analysis of collagen fibre orientation as measured by the number of measurements per ROI (n>6), that exhibit a maximum intensity in an area of 200 x 200 μm$^2$ for different polarizer angles. Samples are characterized in the four ROIs presented in Figure S10: cortical (white), bridged (light grey), interphase (dark grey) and intermonetite (black). The percentage of measurements with maximum intensity per polarizer angle is presented by region and sample. Images of alcian blue and picrosirius red staining of 2M_PFB25_2 sample under six angles of polarized light for each of the corresponding polarizer-analyser angles: 0°-45º, 15°-60º, 30°-75º, 45°-90º, 60°-105º and 75°-120º. Scale bars: 1000 μm.

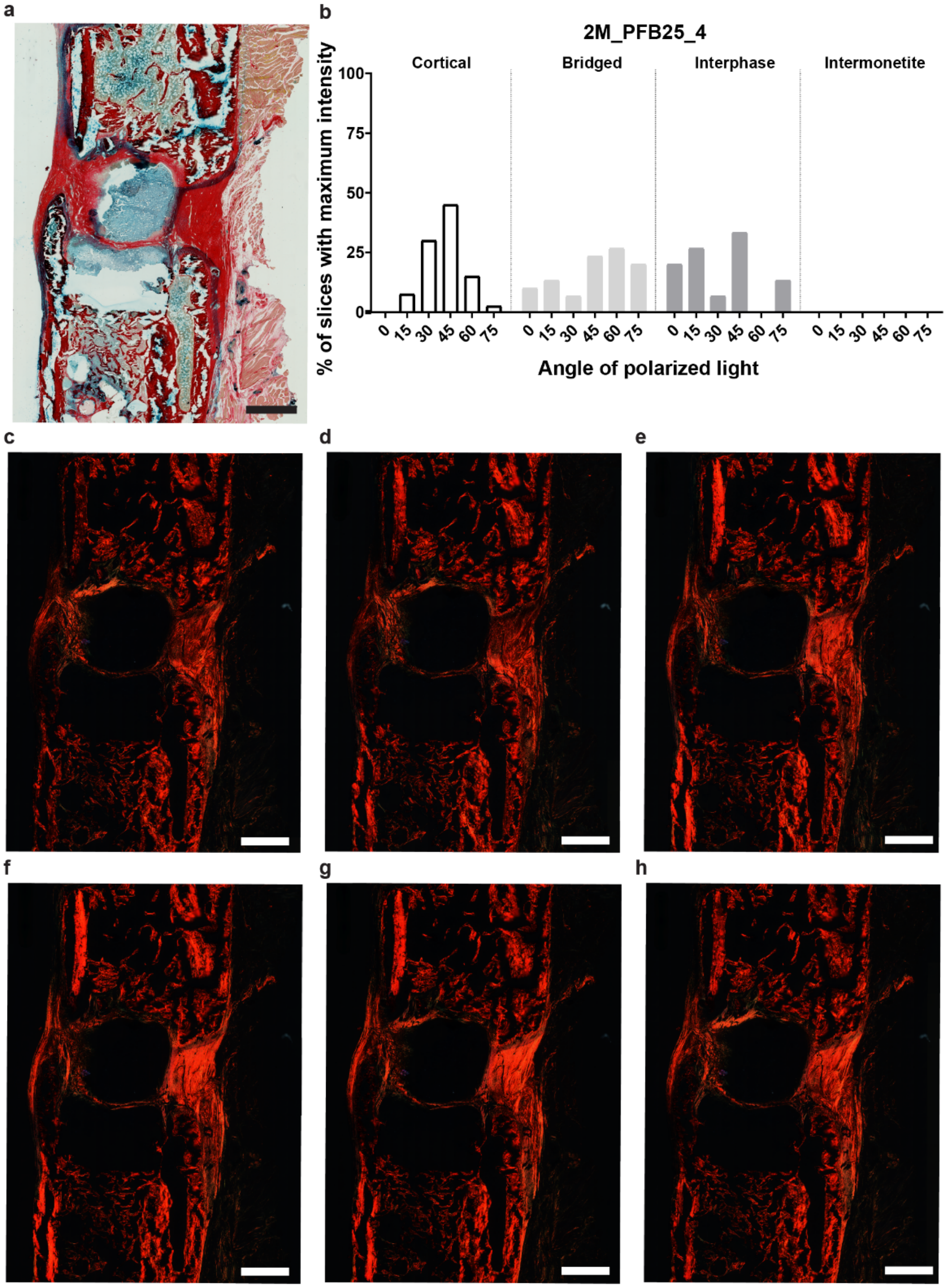


**Figure S16: Collagen fibre orientation analysis performed with polarized light microscopy for sample 2M_PFB25_4.** a) Alcian blue and picrosirius red staining bright field image of sample 2M_PFB25_4. b) Quantitative analysis of collagen fibre orientation as measured by the number of measurements per ROI (n>6), that exhibit a maximum intensity in an area of 200 x 200 μm$^2$ for different polarizer angles. Samples are characterized in the four ROIs presented in Figure S10: cortical (white), bridged (light grey), interphase (dark grey) and intermonetite (black). The percentage of measurements with maximum intensity per polarizer angle is presented by region and sample. Images of alcian blue and picrosirius red staining of 2M_PFB25_4 sample under six angles of polarized light for each of the corresponding polarizer-analyser angles: 0°-45º, 15°-60º, 30°-75º, 45°-90º, 60°-105º and 75°-120º. Scale bars: 1000 μm.

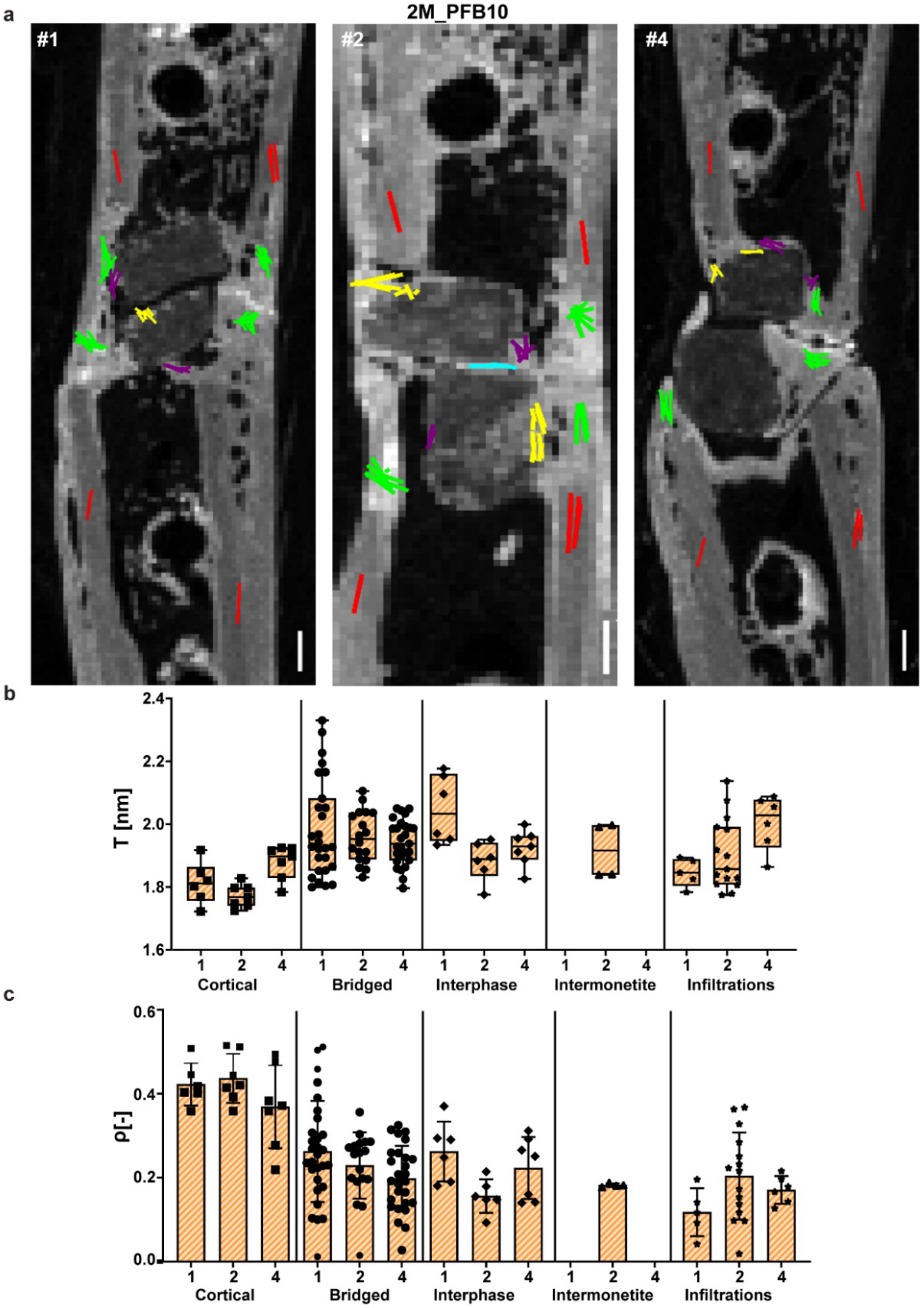


**Figure S17: Image and quantification of the average mineral particle thickness (T) and the degree of alignment (ρ) obtained with SAXS measurements in the 2M_PFB10 experimental group.** Data obtained from samples #1, #2 and #4 in the predefined regions of interest presented in Figure S10. Symbols represent each of the measurements obtained in each region (n>4 per region). Boxplots indicate the mean and standard deviation of all measurements per region, where every dot represents one SAXS or nanoindentation measurement. Bar plots show the median value and the interquartile range per region.

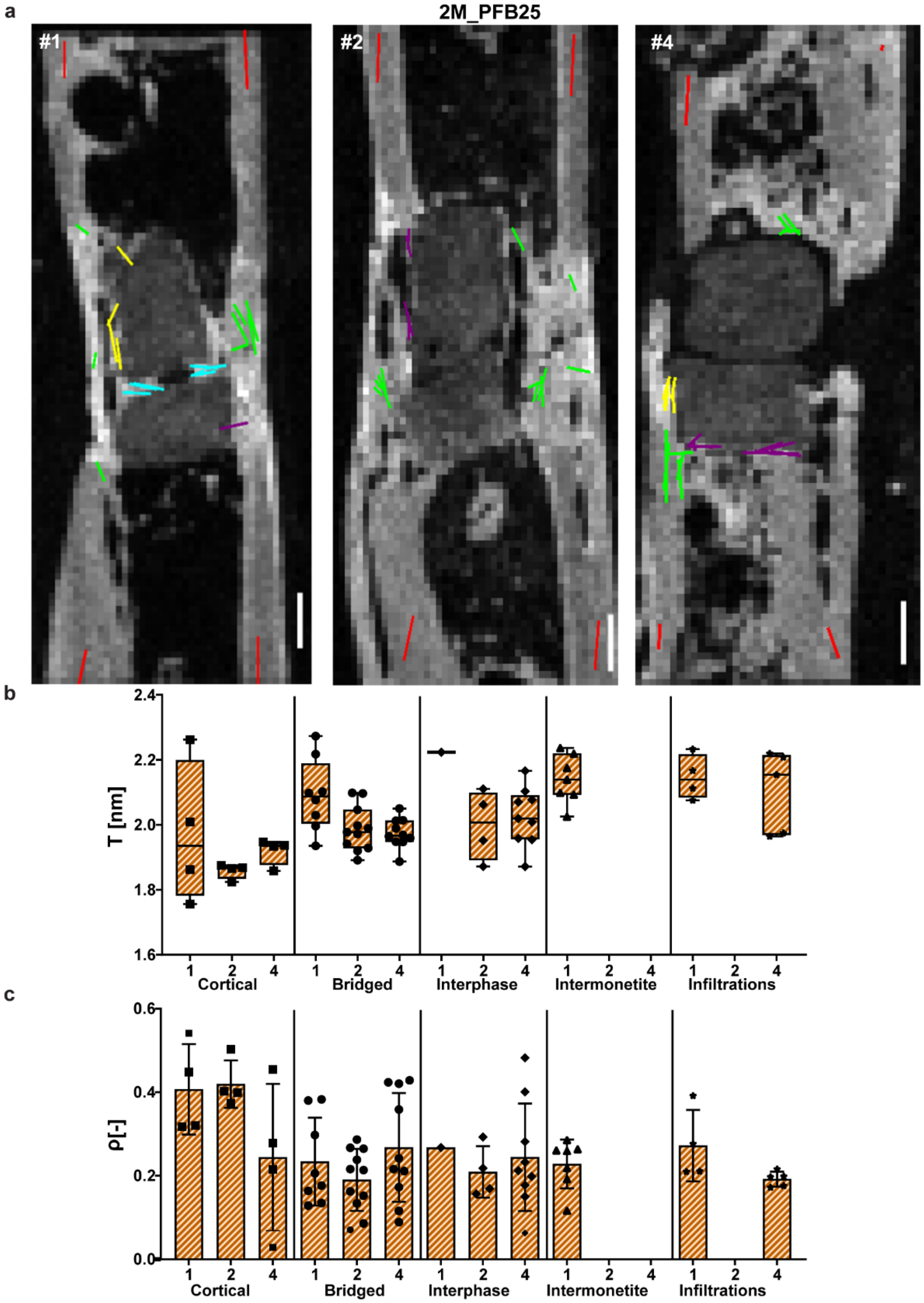


**Figure S18: Image and quantification of the average mineral particle thickness (T) and the degree of alignment (ρ) obtained with SAXS measurements in the 2M_PFB25 experimental group.** Data obtained from samples #1, #2 and #4 in the predefined regions of interest presented in Figure S10. Symbols represent each of the measurements obtained in each region (n>4 per region). Boxplots indicate the mean and standard deviation of all measurements per region, where every dot represents one SAXS or nanoindentation measurement. Bar plots show the median value and the interquartile range per region.

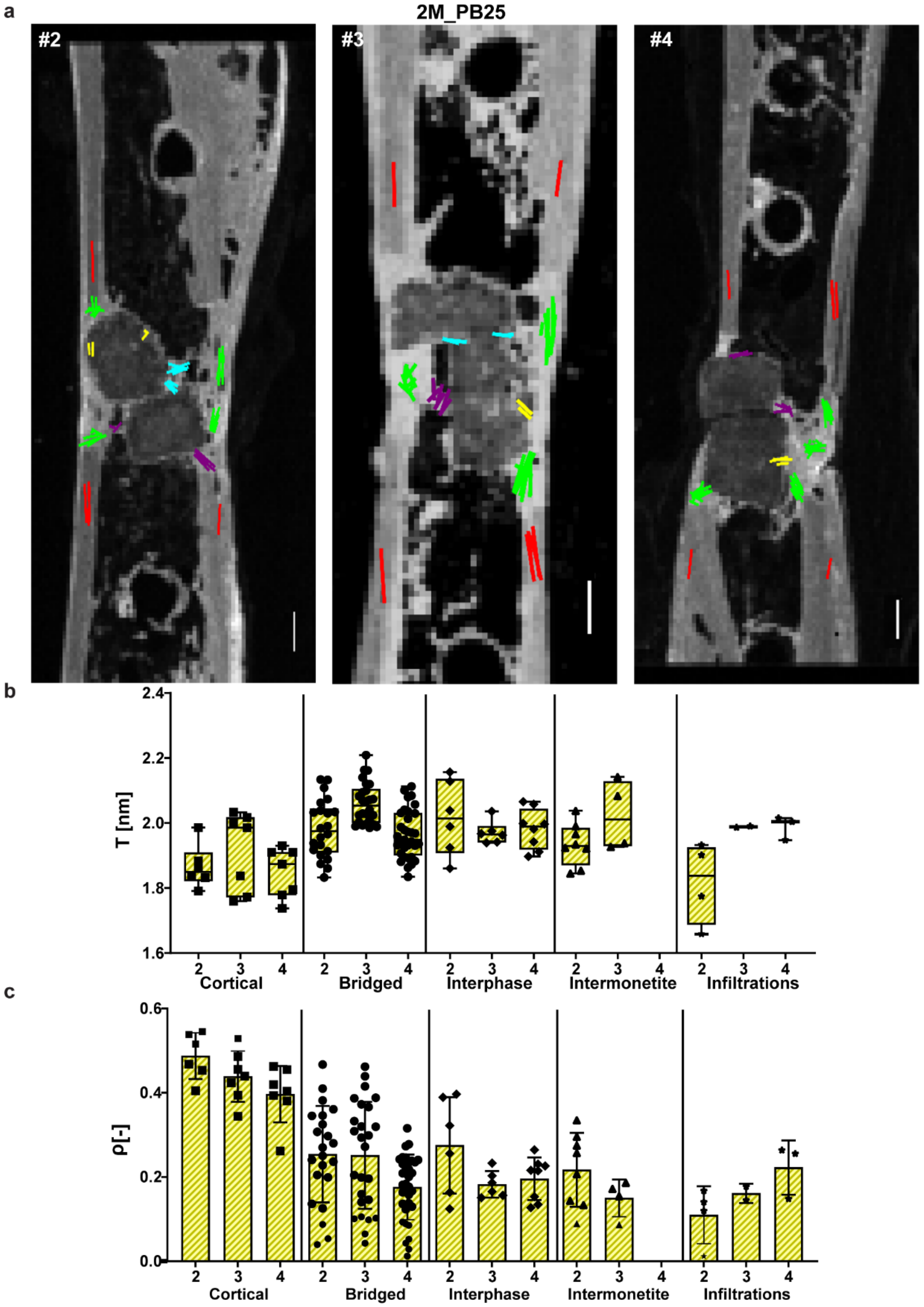


**Figure S19: Image and quantification of the average mineral particle thickness (T) and the degree of alignment (ρ) obtained with SAXS measurements in the 2M_PB25 experimental group.** Data obtained from samples #2, #3 and #4 in the predefined regions of interest presented in Figure S10. Symbols represent each of the measurements obtained in each region (n>4 per region). Boxplots indicate the mean and standard deviation of all measurements per region, where every dot represents one SAXS or nanoindentation measurement. Bar plots show the median value and the interquartile range per region.

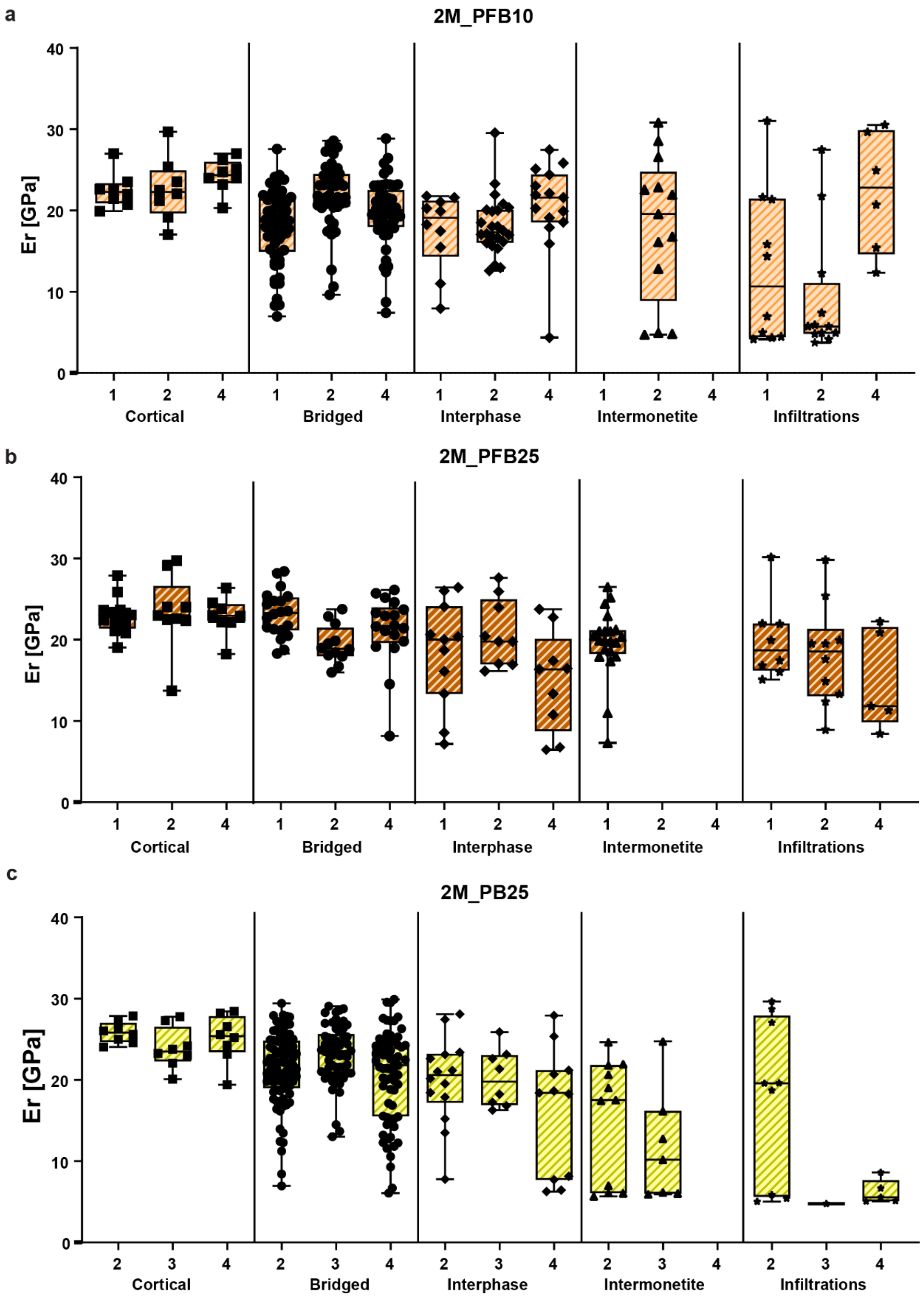


**Figure S20. Quantification of local elastic indentation modulus (Er)**. Measurements obtained in the predefined regions of interest presented in Figure S10 in the selected experimental groups, including a) 2M_PFB10, b) 2M_PFB25 and c) 2M_PB25. Symbols represent each of the measurements obtained in each region (n>5 per region) and sample. Boxplots indicate the median value and the interquartile range of all measurements per region and sample.

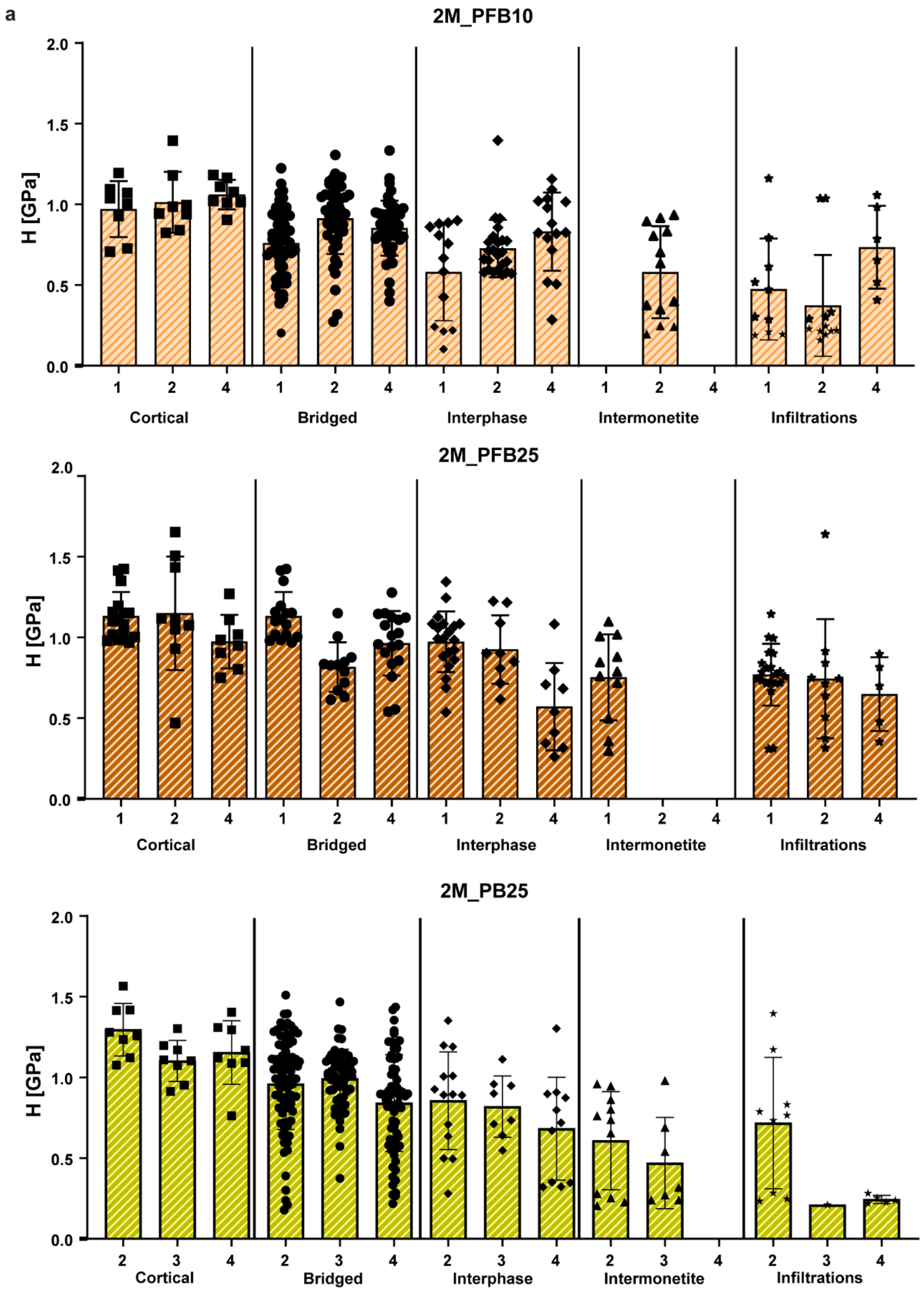


**Figure S21. Quantification of hardness (H).** Measurements obtained in the predefined regions of interest presented in Figure S10 in the selected experimental groups, including a) 2M_PFB10, b) 2M_PFB25 and c) 2M_PB25. Symbols represent each of the measurements obtained in each region (n>5 per region) and sample. Bar plots indicate the mean ± standard deviation of all measurements per region and sample.

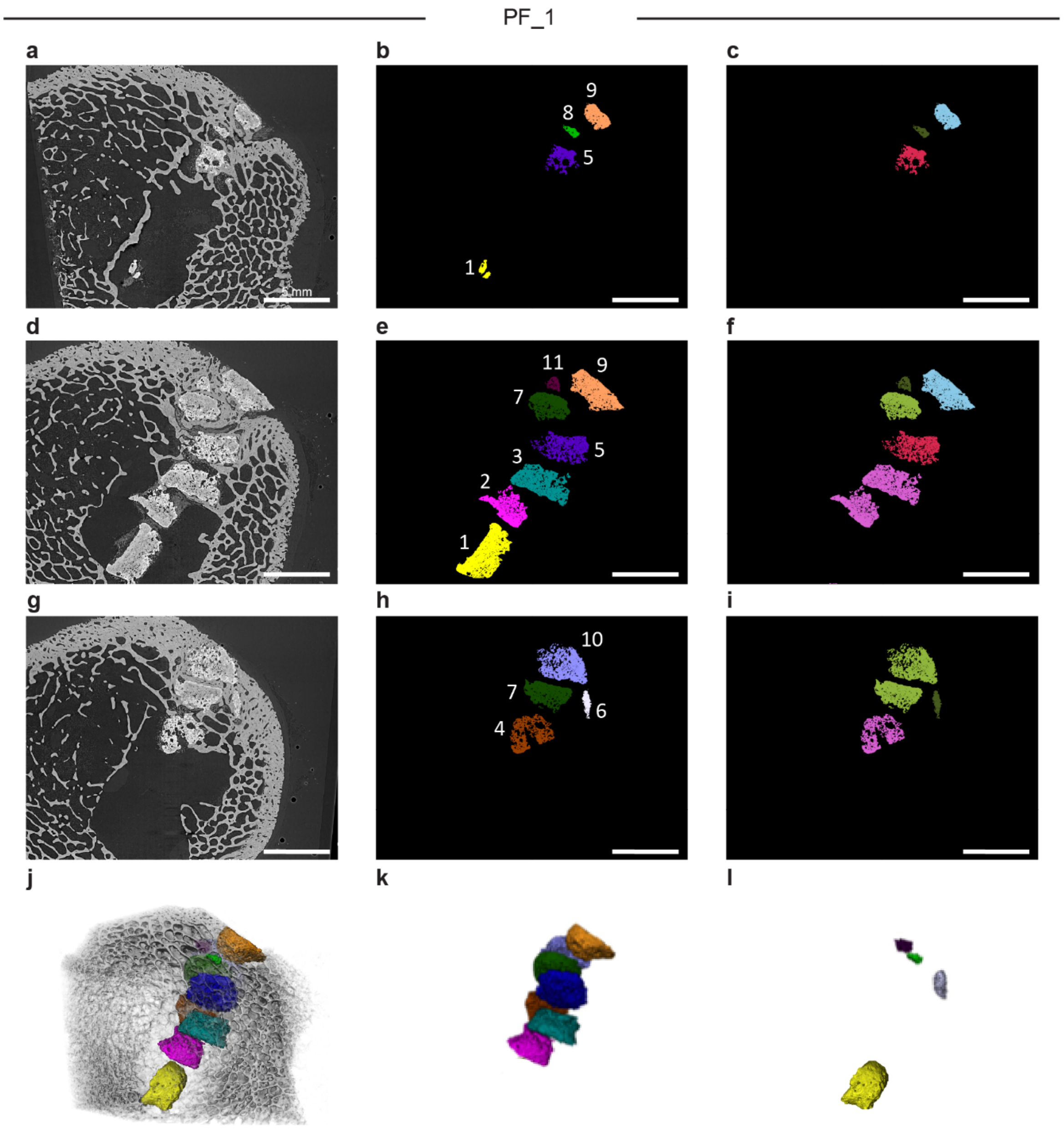


**Figure S22. Representative cross-sections and 3D reconstructions of synchrotron microCT images with segmented data visualizing all monetites within sample PF_1.** a-i) Representative cross-sections of synchrotron microCT and segmented data visualizing all monetites within sample: a), d), g) tomographic images; b), e), h) individual monetites, with individual color and number by label; c), f), i) monetites assigned to group by anatomical location (blue = periost, green = cortical, red = endost, pink = marrow, dark green = discarded monetite). j) 3D rendering of entire sample including all individual monetites; k) visualization of monetites included in the analysis, l) visualization of monetite fragments not included in the analysis. Scale bars = 5 mm.

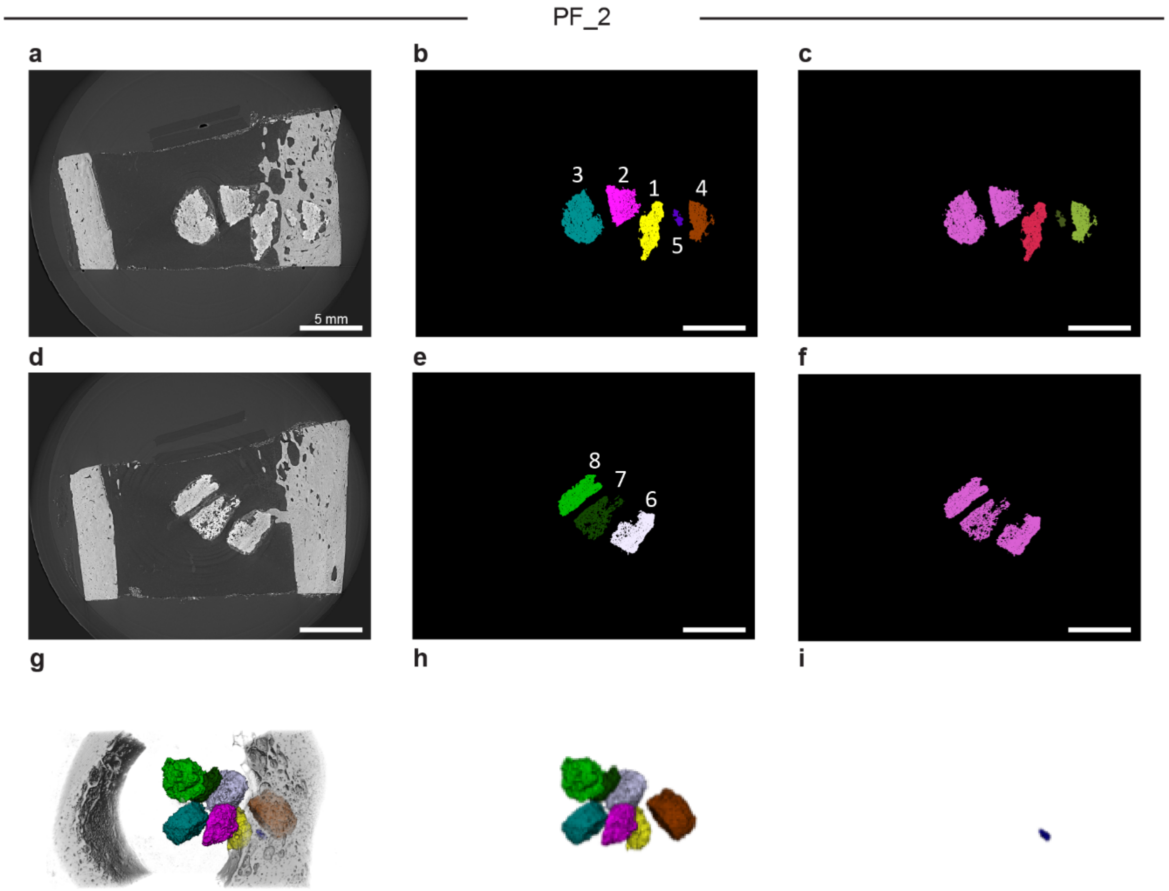


**Figure S23. Representative cross-sections and 3D reconstructions of synchrotron microCT images with segmented data visualizing all monetites within sample PF_2.** a-f) Representative cross-sections of synchrotron microCT and segmented data visualizing all monetites within sample: a), d) tomographic images; b), e) individual monetites, with individual color and number by label; c), f) monetites assigned to group by anatomical location (green = cortical, red = endost, pink = marrow, dark green = discarded monetite). g) 3D rendering of entire sample including all individual monetites; h) visualization of monetites included in the analysis, i) visualization of monetite fragments not included in the analysis. Scale bars = 5 mm.

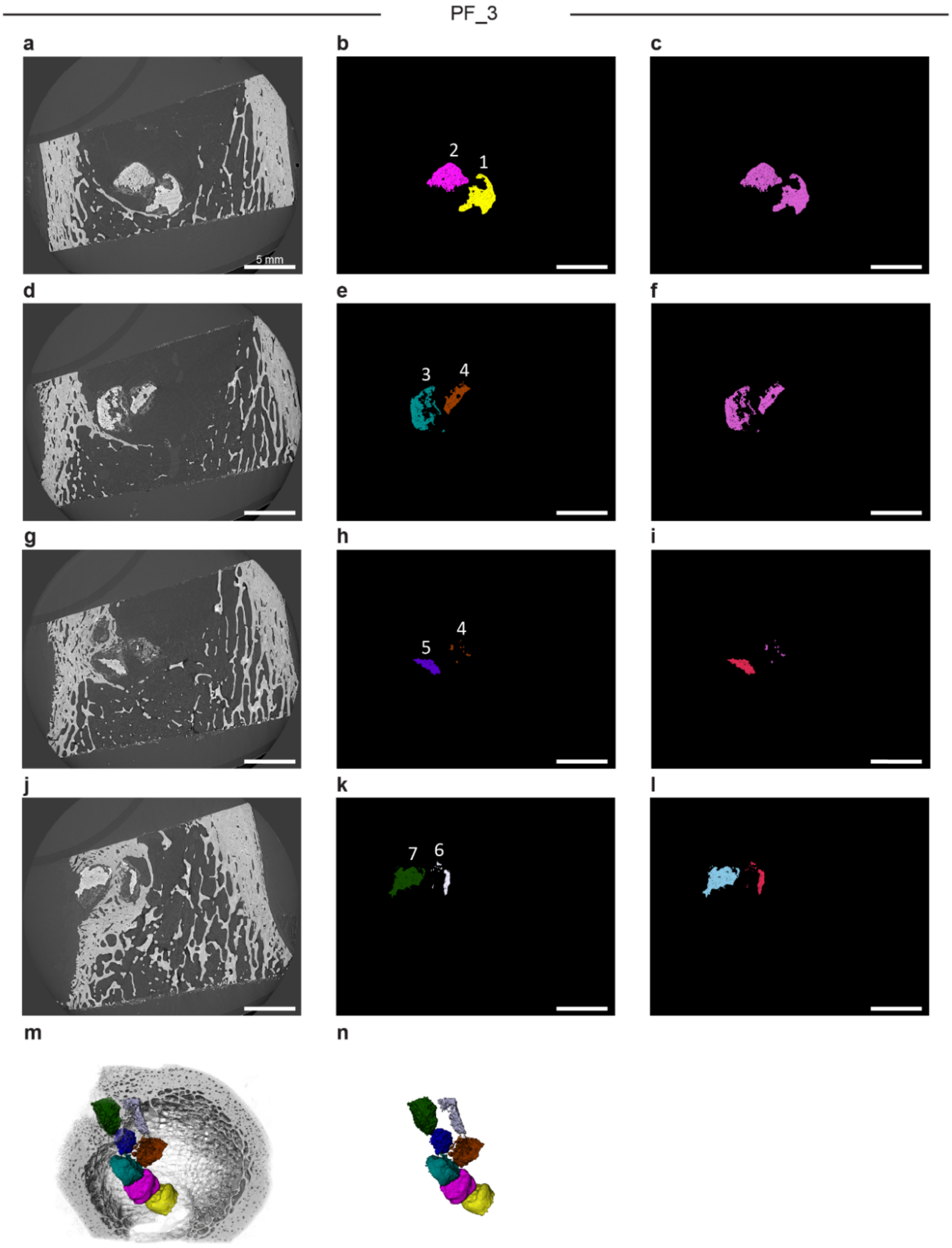


**Figure S24. Representative cross-sections and 3D reconstructions of synchrotron microCT images with segmented data visualizing all monetites within sample PF_3.** a-l) Representative cross-sections of synchrotron microCT and segmented data visualizing all monetites within sample: a), d), g), j) tomographic images; b), e), h), k) individual monetites, with individual color and number by label; c), f), i), l) monetites assigned to group by anatomical location (red = endost, pink = marrow). m) 3D rendering of entire sample including all individual monetites; n) visualization of monetites included in the analysis (all of them). Scale bars = 5 mm.

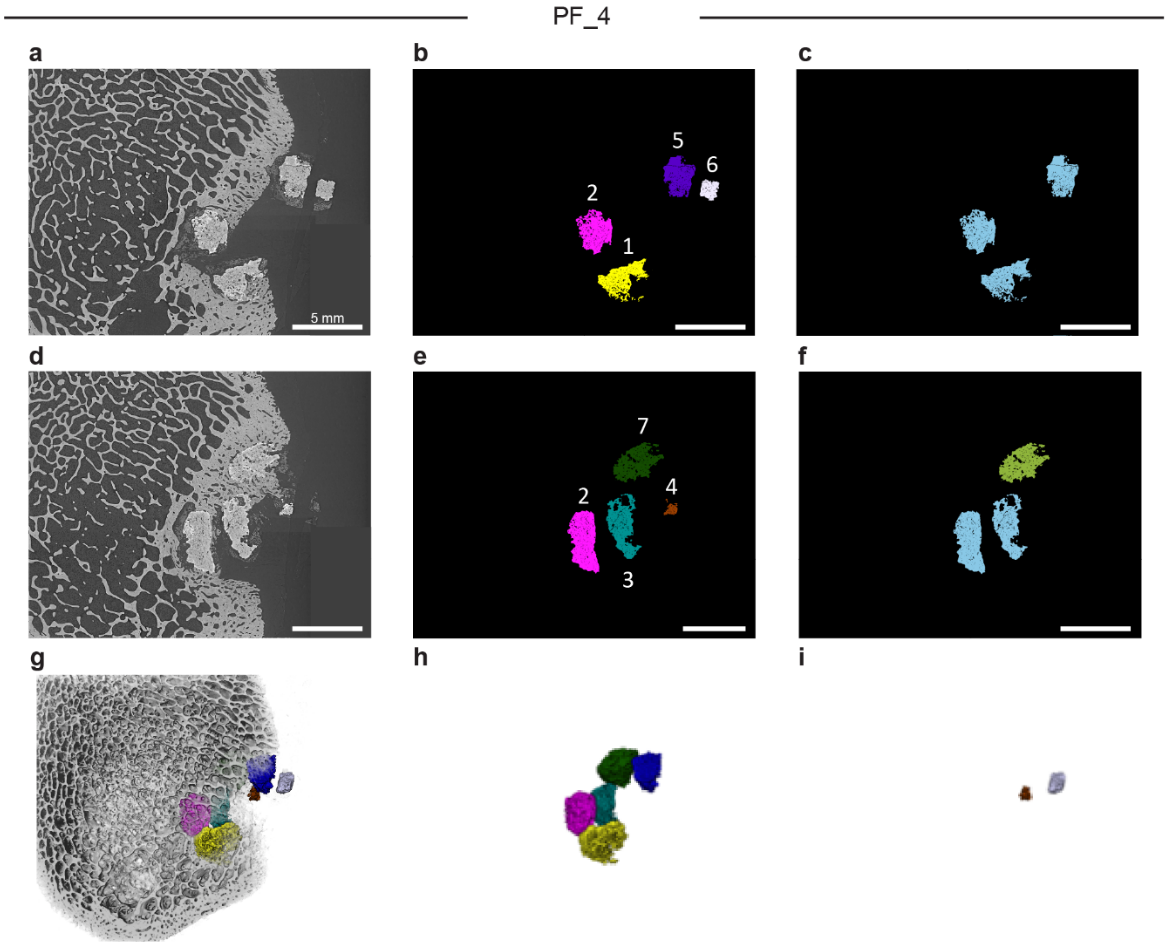


**Figure S25. Representative cross-sections and 3D reconstructions of synchrotron microCT images with segmented data visualizing all monetites within sample PF_4.** a-f) Representative cross-sections of synchrotron microCT and segmented data visualizing all monetites within sample: a), d) tomographic images; b), e) individual monetites, with individual color and number by label; c), f) monetites assigned to group by anatomical location (blue = periost, green = cortical, discarded monetite fragments are deleted). g) 3D rendering of entire sample including all individual monetites; h) visualization of monetites included in the analysis, i) visualization of monetite fragments not included in the analysis. Scale bars = 5 mm.

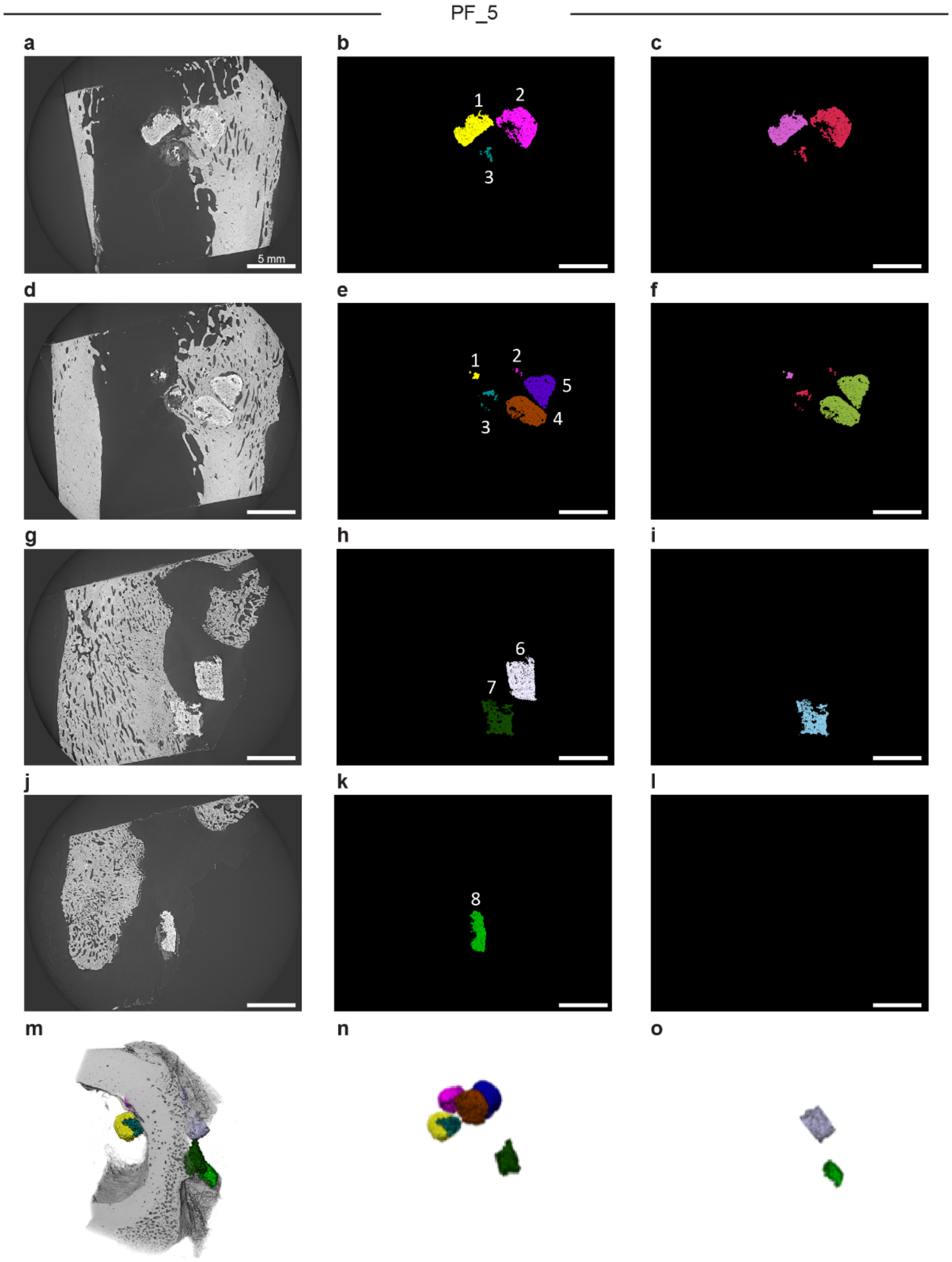


**Figure S26. Representative cross-sections and 3D reconstructions of synchrotron microCT images with segmented data visualizing all monetites within sample PF_5.** a-l) Representative cross-sections of synchrotron microCT and segmented data visualizing all monetites within sample: a), d), g), j) tomographic images; b), e), h), k) individual monetites, with individual color and number by label; c), f), i), l) monetites assigned to group by anatomical location (blue = periost, green = cortical, red = endost, pink = marrow, discarded monetite fragments are deleted). m) 3D rendering of entire sample including all individual monetites; n) visualization of monetites included in the analysis, o) visualization of monetite fragments not included in the analysis. Scale bars = 5 mm.

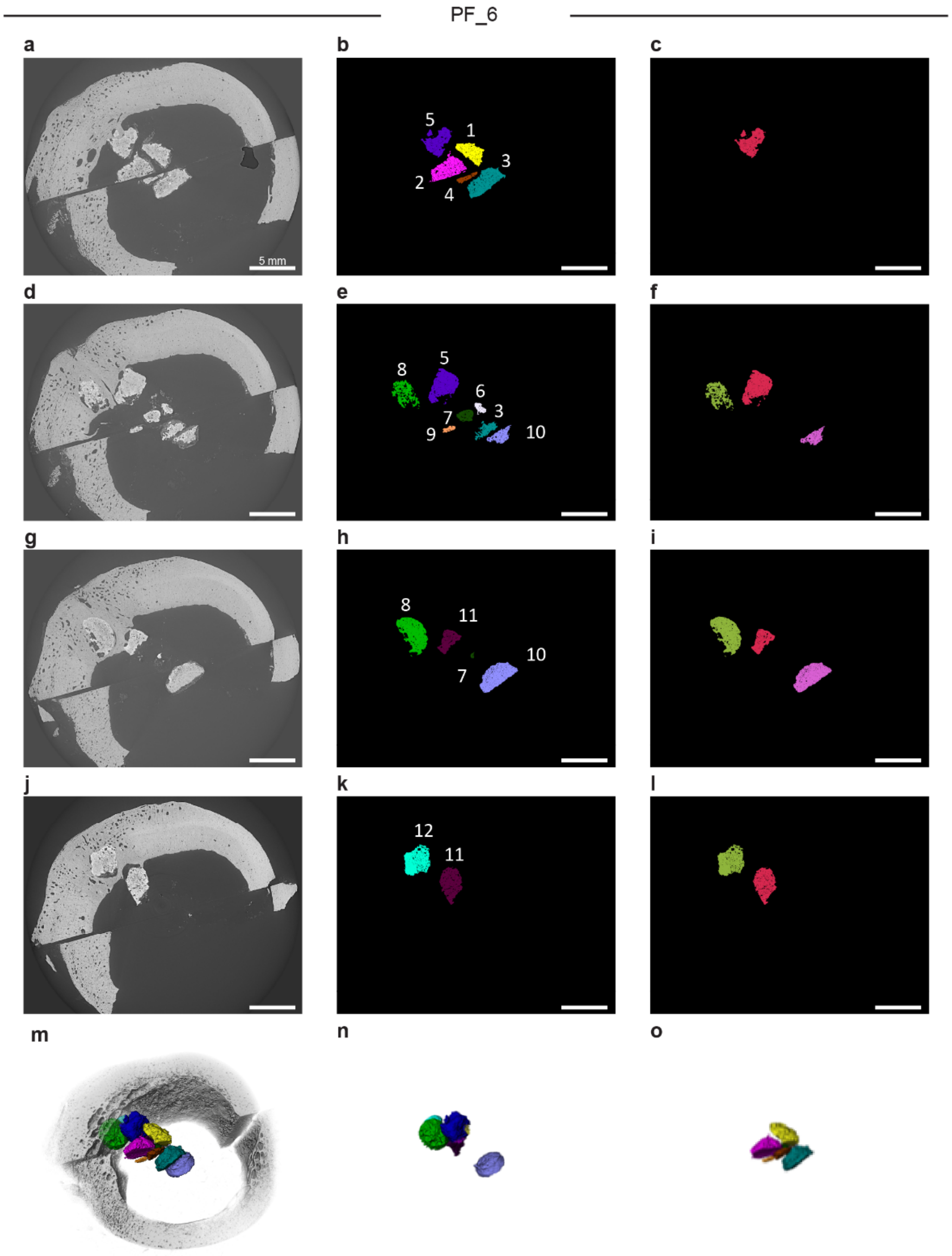


**Figure S27. Representative cross-sections and 3D reconstructions of synchrotron microCT images with segmented data visualizing all monetites within sample PF_6.** a-l) Representative cross-sections of synchrotron microCT and segmented data visualizing all monetites within sample: a), d), g), j) tomographic images; b), e), h), k) individual monetites, with individual color and number by label; c), f), i), l) monetites assigned to group by anatomical location (green = cortical, red = endost, pink = marrow, discarded monetite fragments are deleted). m) 3D rendering of entire sample including all individual monetites; n) visualization of monetites included in the analysis, o) visualization of monetite fragments not included in the analysis. Scale bars = 5 mm.

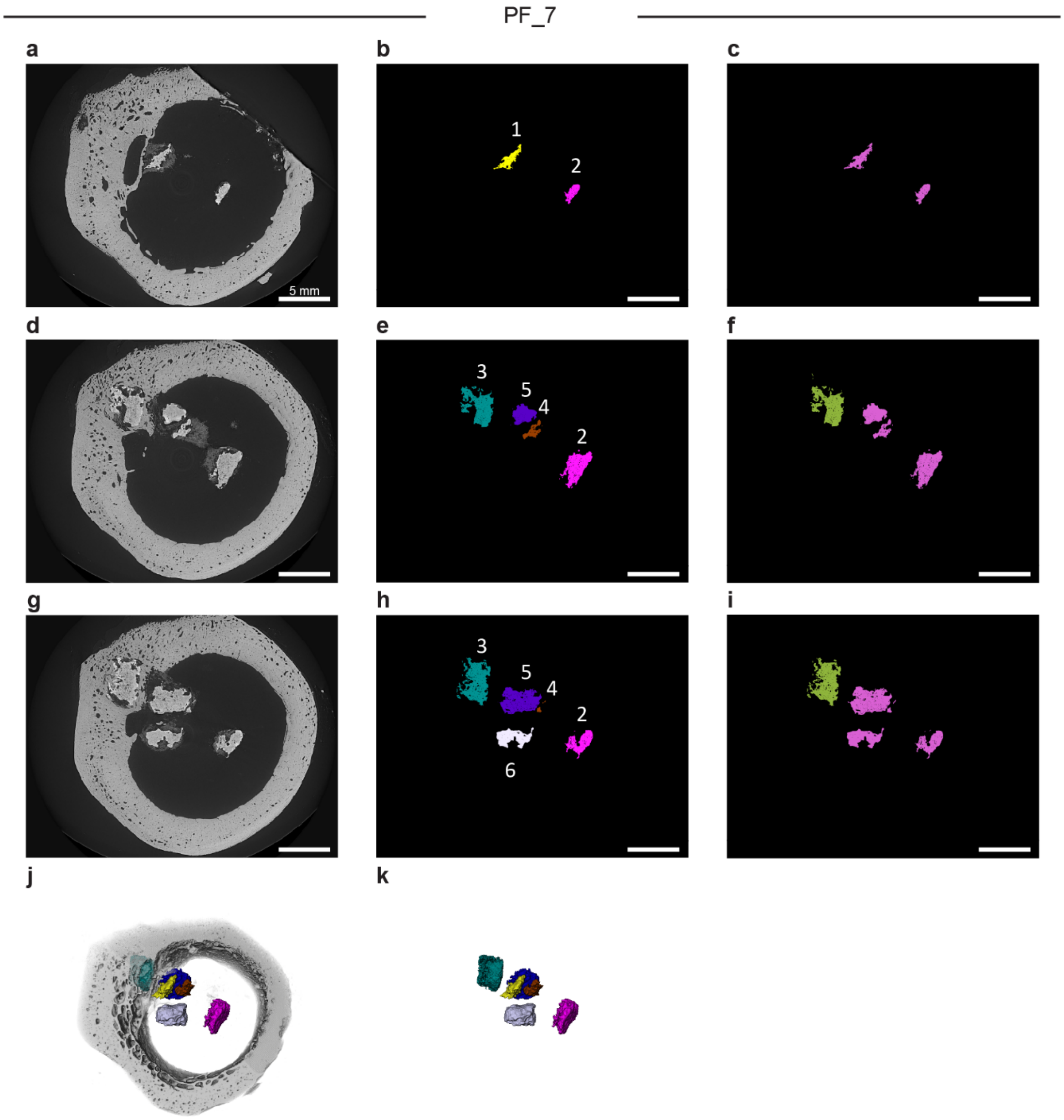


**Figure S28. Representative cross-sections and 3D reconstructions of synchrotron microCT images with segmented data visualizing all monetites within sample PF_7.** a-i) Representative cross-sections of synchrotron microCT and segmented data visualizing all monetites within sample: a), d), g) tomographic images; b), e), h) individual monetites, with individual color and number by label; c), f), i) monetites assigned to group by anatomical location (green = cortical, pink = marrow). j) 3D rendering of entire sample including all individual monetites; k) visualization of monetites included in the analysis (all of them). Scale bars = 5 mm.

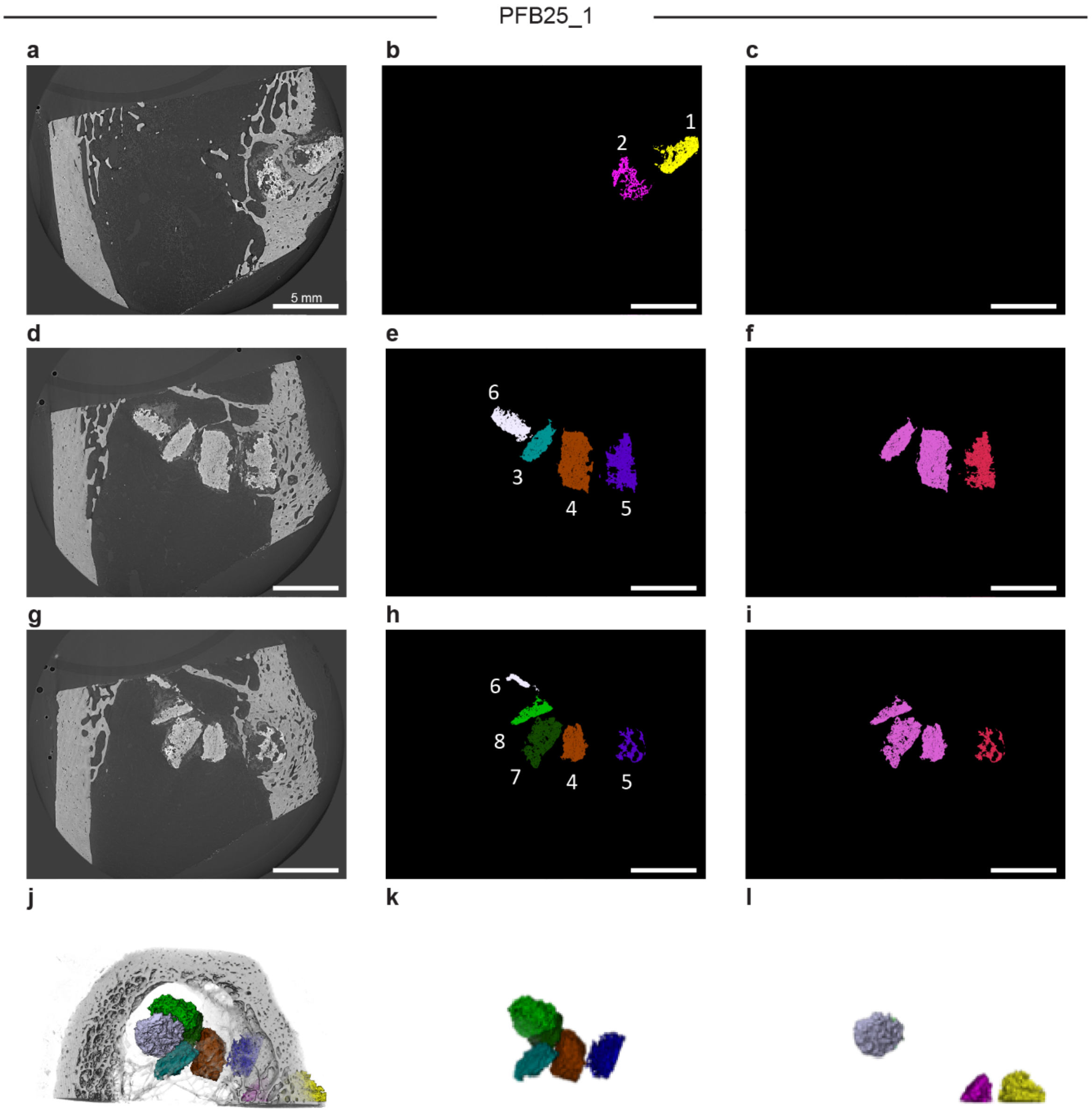


**Figure S29. Representative cross-sections and 3D reconstructions of synchrotron microCT images with segmented data visualizing all monetites within sample PFB25_1.** a-i) Representative cross-sections of synchrotron microCT and segmented data visualizing all monetites within sample: a), d), g) tomographic images; b), e), h) individual monetites, with individual color and number by label; c), f), i) monetites assigned to group by anatomical location (red = endost, pink = marrow, discarded monetite fragments are deleted). j) 3D rendering of entire sample including all individual monetites; k) visualization of monetites included in the analysis, l) visualization of monetite fragments not included in the analysis. Scale bars = 5 mm.

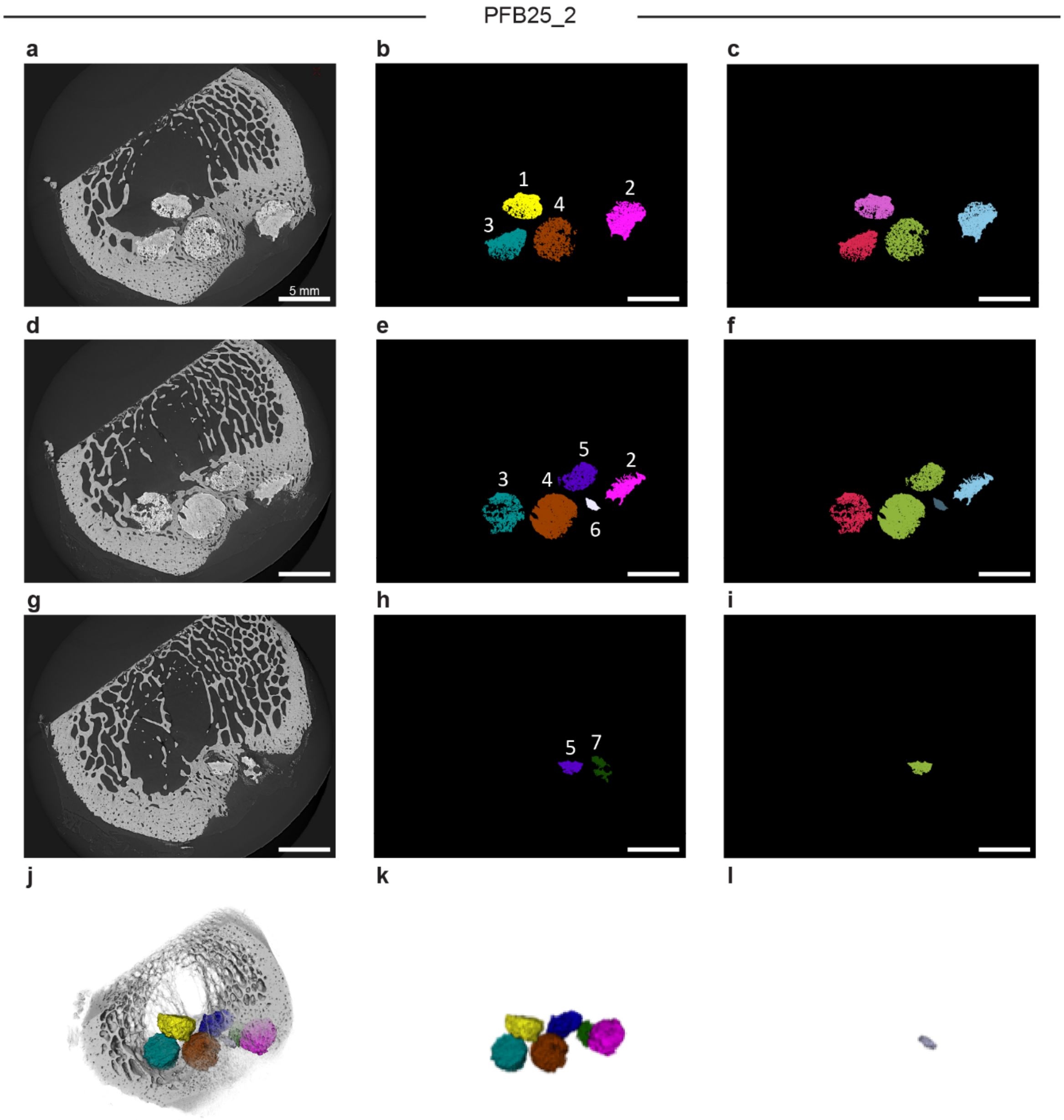


**Figure S30. Representative cross-sections and 3D reconstructions of synchrotron microCT images with segmented data visualizing all monetites within sample PFB25_2.** a-i) Representative cross-sections of synchrotron microCT and segmented data visualizing all monetites within sample: a), d), g) tomographic images; b), e), h) individual monetites, with individual color and number by label; c), f), i) monetites assigned to group by anatomical location (blue = periost, green = cortical, red = endost, pink = marrow, dark green = discarded monetite). j) 3D rendering of entire sample including all individual monetites; k) visualization of monetites included in the analysis, l) visualization of monetite fragments not included in the analysis. Scale bars = 5 mm.

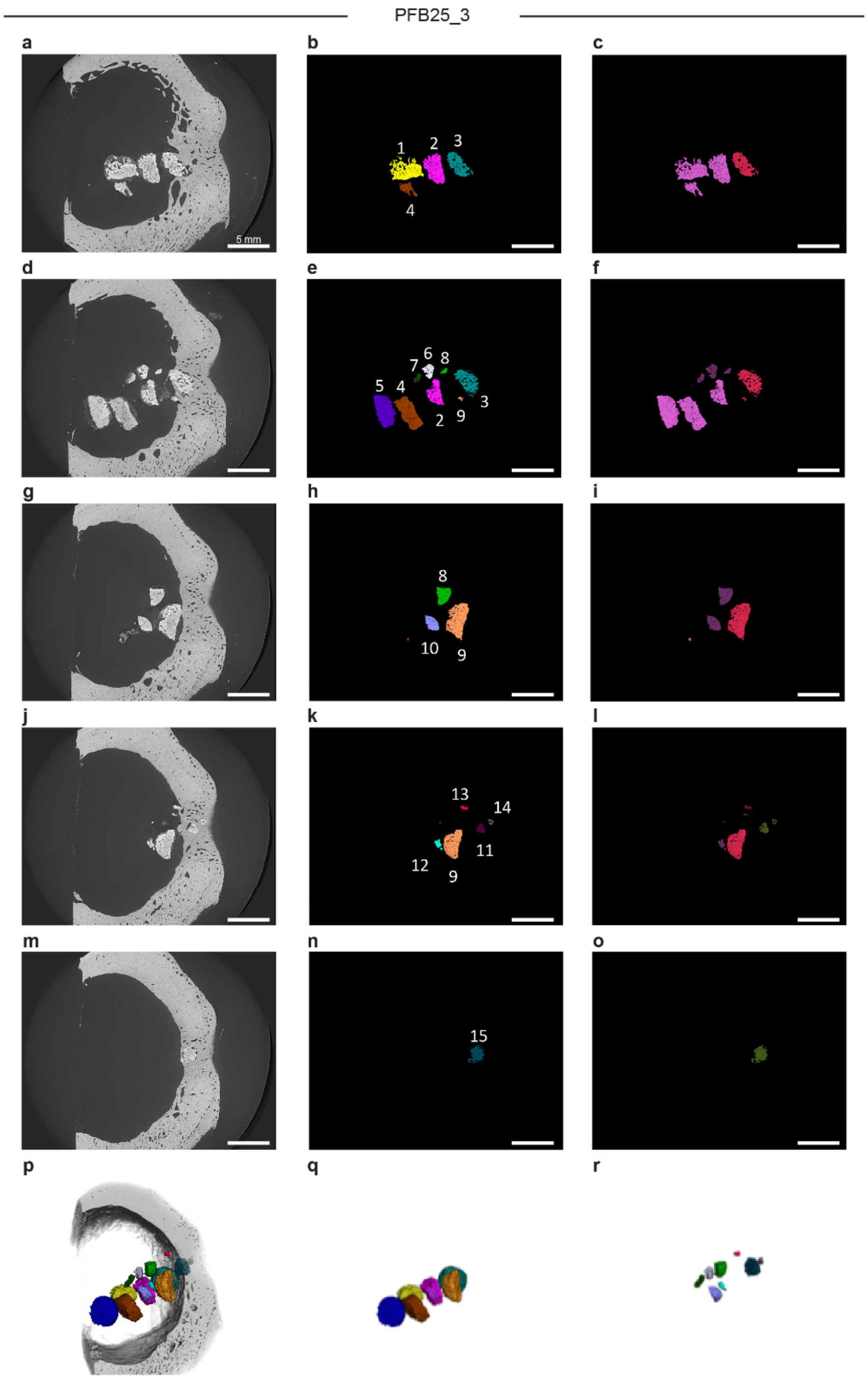


**Figure S31. Representative cross-sections and 3D reconstructions of synchrotron microCT images with segmented data visualizing all monetites within sample PFB25_3.** a-o) Representative cross-sections of synchrotron microCT and segmented data visualizing all monetites within sample: a), d), g), j), m) tomographic images; b), e), h), k), n) individual monetites, with individual color and number by label; c), f), i), l), o) monetites assigned to group by anatomical location (green = cortical, red = endost, pink = marrow, purple = discarded monetite fragments). p) 3D rendering of entire sample including all individual monetites; q) visualization of monetites included in the analysis, r) visualization of monetite fragments not included in the analysis. Scale bars = 5 mm.

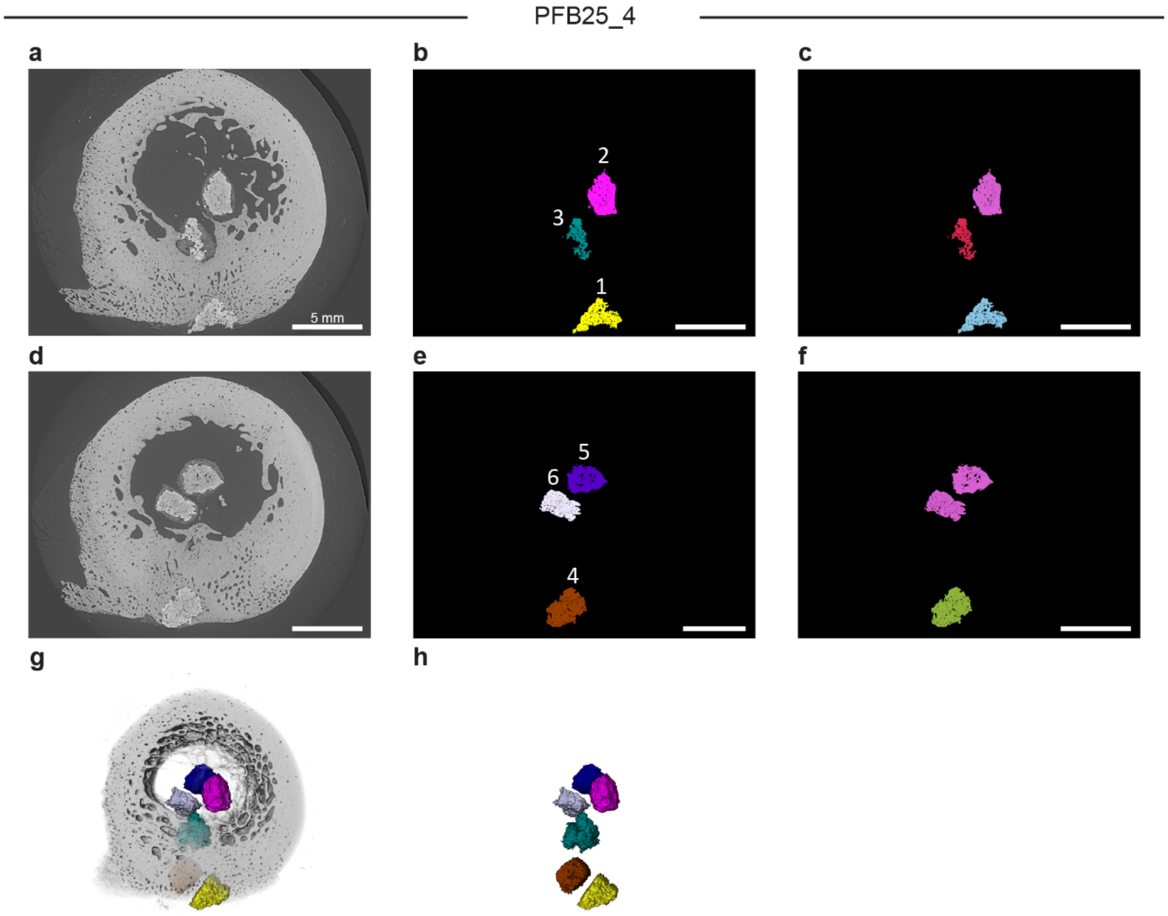


**Figure S32. Representative cross-sections and 3D reconstructions of synchrotron microCT images with segmented data visualizing all monetites within sample PFB25_4.** a-f) Representative cross-sections of synchrotron microCT and segmented data visualizing all monetites within sample: a), d) tomographic images; b), e) individual monetites, with individual color and number by label; c), f) monetites assigned to group by anatomical location (blue = periost, green = cortical, red = endost, pink = marrow). g) 3D rendering of entire sample including all individual monetites; h) visualization of monetites included in the analysis (all of them). Scale bars = 5 mm.

**Table S1. Primers used for qRT-PCR**

| Gene Name | Primer sequence 5'→3' |
|---|---|
| *GAPDH* | Forward: TCAAGGCTGAGAACGGGAA |
| | Reverse: TGGGTGGCAGTGATGGCA |
| *BGLAP* | Forward: CAGCGAGGTAGTGAAGAGACC |
| | Reverse: TCTGGAGTTTATTTGGGAGCAG |
| *SPARC* | Forward: AGAATGAGAAGCGCCTGGAG |
| | Reverse: CTGCCAGTGTACAGGGAAGA |

**Video S1. 3D reconstruction of synchrotron microCT images of an ovine sample.** The sequence of the segmentation process can be observed identifying all monetites by anatomical location, bone volume in the vicinity of the monetite and bone infiltrations.

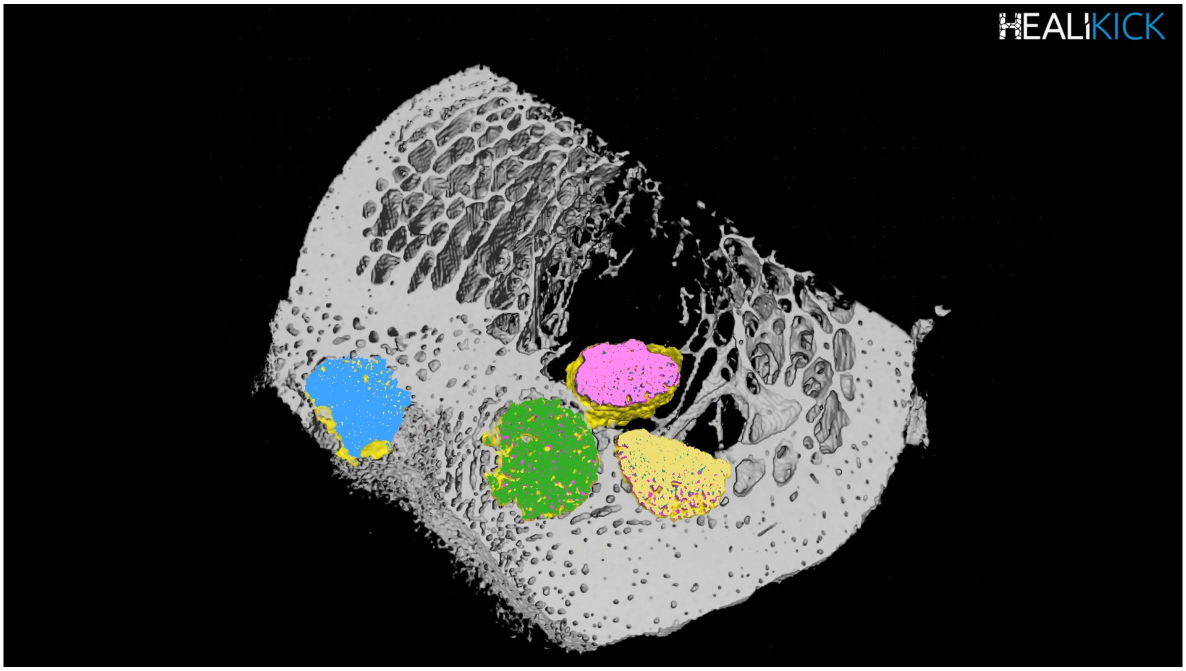
